# Epigenetic engineering of yeast reveals dynamic molecular adaptation to methylation stress and genetic modulators of specific DNMT3 family members

**DOI:** 10.1101/2020.01.26.919936

**Authors:** Alex I. Finnegan, Somang Kim, Hu Jin, Michael Gapinske, Wendy S. Woods, Pablo Perez-Pinera, Jun S. Song

**Author notes:** Correspondence: JSS and PPP. equal contributions (Alex I. Finnegan, Somang Kim).

## Abstract

Cytosine methylation is a ubiquitous modification in mammalian DNA generated and maintained by several DNA methyltransferases (DNMTs) with partially overlapping functions and genomic targets. To systematically dissect the factors specifying each DNMT’s activity, we engineered combinatorial knock-in of human DNMT genes in *Komagataella phaffii*, a yeast species lacking endogenous DNA methylation. Time-course expression measurements captured dynamic network-level adaptation of cells to DNMT3B1-induced DNA methylation stress and showed that coordinately modulating the availability of S-adenosyl methionine (SAM), the essential metabolite for DNMT-catalyzed methylation, is an evolutionarily conserved epigenetic stress response, also implicated in several human diseases. Convolutional neural networks trained on genome-wide CpG-methylation data learned distinct sequence preferences of DNMT3 family members. A simulated annealing interpretation method resolved these preferences into individual flanking nucleotides and periodic poly(A) tracts that rotationally position highly methylated cytosines relative to phased nucleosomes. Furthermore, the nucleosome repeat length defined the spatial unit of methylation spreading. Gene methylation patterns were similar to those in mammals, and hypo- and hypermethylation were predictive of increased and decreased transcription relative to control, respectively, in the absence of mammalian readers of DNA methylation. Introducing controlled epigenetic perturbations in yeast thus enabled characterization of fundamental genomic features directing specific DNMT3 proteins.

## INTRODUCTION

The formation and removal of 5-methylcytosines (5mCs) at CpG dinucleotides constitute critical steps in mammalian development, pathogenesis and aging (1–4). In addition to being associated with various aspects of transcriptional activities (5–7), DNA methylation also provides epigenetic memory of cell identity (8). Moreover, aberrant DNA methylation is frequently found in multiple diseases, highlighting the importance of studying this epigenetic modification to help improve disease detection, diagnosis and treatment (9, 10). A key problem that needs to be addressed to improve our understanding is how DNA methylation is established and maintained at specific genomic loci by catalytic enzymes.

It is known that DNA methylation is accomplished by a set of DNA methyltransferases (DNMTs). For example, DNMT1 recognizes hemi-methylated CpG sites and methylates the opposite unmethylated cytosine during cell division. Likewise, DNMT3A and DNMT3B can establish *de novo* methylation of unmethylated CpG dinucleotides, while DNMT3L enhances the catalytic activity of these enzymes across the genome (11). The partially overlapping functions of the DNMTs and the omnipresence of these enzymes in mammalian cells pose a challenge in systematically dissecting the binding specificity and functional consequences of individual DNMTs. To overcome these difficulties, we engineered a combination of human DNMT genes into *Komagataella phaffii* yeast cells that do not possess endogenous DNA methylation and studied the high-throughput epigenetic and transcriptomic profiles of these perturbed cells via whole-genome bisulfite sequencing and RNA-seq. We demonstrate that this controlled epigenetic perturbation helps address several outstanding questions: (1) What is the pattern of *de novo* DNA methylation established by a particular set of DNMTs? (2) To what extent does genetic information encoded in DNA influence CpG methylation? (3) What is the functional consequence of DNA methylation on gene expression? (4) Which molecular stress responses do cells utilize to adapt to ectopic DNA methylation?

Applying our statistical interpretation method (12) to convolutional deep neural networks trained on whole-genome bisulfite sequencing data, we show that different DNMTs have distinct patterns of sequence preference and aversion, similar to those previously found using an episomal DNA methylation assay in human HEK293c18 cells (13). Importantly, our time-course measurements of epigenetic and transcriptomic states allow us to disentangle direct and indirect effects of DNA methylation on gene expression changes in *K. phaffii*; in addition to the genes transcriptionally altered by direct gene-body DNA methylation, cells transiently modulate their expression pattern to counter the stress of exogenous DNA methylation. Specifically, biosynthesis of S-adenosylmethionine (SAM), the principal methyl donor for DNA methylation, is significantly impaired in *K. phaffii* at a network level by multiple genes that coordinately change expression during early days post DNMT induction. Intriguingly, the cellular level of SAM has been previously implicated in regulating the methylome dynamics of Schwann cells during peripheral nerve myelination in mouse, with decreased and increased SAM levels being associated with demethylation and hypermethylation, respectively (14). Our work thus shows that modulating the SAM level is an ancient molecular mechanism, conserved across species for controlling DNA methylation, that is also available to yeast as a means to adapt to exogenous epigenetic stress. By integrating our data with local chromatin conformation information, we further find evidence for either rotational positioning of nucleosomes flanking methylated CpG sites or rotational positioning of CpG itself on a single nucleosome, suggesting that the geometric orientation of accessible CpGs with respect to histones may play a role in facilitating DNMT activities.

Several previous studies have also examined DNA methylation from different perspectives. At the level of individual cytosines, efforts to identify DNA sequence determinants of *de novo* methylation have produced consensus motifs from the sequences flanking CpGs with high and low levels of methylation and characterized the methylation preferences of individual DNMTs *in vitro* (13, 15). Meanwhile, at the level of histone modification, knockout of DNMTs in mouse embryonic stem cells followed by reintroduction of individual *de novo* DNMTs found differential localization patterns between the isoforms DNMT3A2 and DNMT3B1, demonstrating the recruitment of DNMT3B1 but not DNMT3A2 by H3K36 trimethylation (H3K36me3) (16). The relationship between DNMT3B1 and H3K36me3 was also confirmed by knocking in the DNMT in *S. cerevisiae* (17); this study in *S. cerevisiae* also demonstrated exclusion of DNMT3B1 from regions of H3K4 trimethylation (H3K4me3) and concluded that while the DNMT3B1 introduction produced patterns of DNA methylation similar to those in mammals, the resulting *de novo* DNA methylation did not produce large changes in transcriptional output, nor did it associate with differentially expressed genes (17).

Our approach of introducing controlled combinatorial epigenetic perturbations represents a systematic dissection of DNMT activities and sheds light on the genetic determinants of *de novo* DNA methylation and the functional consequences of methylation on gene expression. From an evolutionary perspective, our findings also suggest that the fundamental architecture of metabolic and epigenetic regulatory networks is broadly shared between yeast and mammals, to the extent that it can readily incorporate feedback from exogenous DNMTs and sense DNA methylation in the ordinarily unmethylated *K. phaffii* genome.

## MATERIAL AND METHODS

### Yeast strains

Wild-type *Komagataella phaffii* NRRL Y-11430, ATCC 76273 was previously used to construct a strain harboring a recombinase landing pad in the Trp2 locus in the *K. phaffii* genome (18). All plasmids utilized in this work were transformed into this strain at the GAP locus.

### Cloning of DNMTs in *K. phaffii* expression plasmids

The open reading frames for DNMT1A, DNMT3A1, DNMT3A2, DNMT3B1, and DNMT3L were purchased from Addgene (plasmid #36939 (DNMT1A), plasmid #35521 (DNMT3A1), plasmid #36941 (DNMT3A2), plasmid #35522 (DNMT3B1) and plasmid #35523 (DNMT3L), and cloned using Gibson Assembly into the *K. phaffii* expression plasmid PP162 (Addgene plasmid #78995). The cloning primers added an SV40 nuclear localization signal (NLS) at both the 5’ and 3’ end of each DNMT to ensure proper nuclear localization and access to genomic DNA. For construction of plasmids expressing combinations of DNMTs, we first cloned the DNMT3L gene using Gibson Assembly into the *K. phaffii* expression plasmid PP164 (Addgene plasmid #78988); the resulting DNMT3L expression cassette driven by a ppTEF1 promoter was then cloned using Gibson Assembly into each of the constructs expressing single DNMTs described above (Supplementary Table S1). A control plasmid containing the selectable marker ZeoR, but no *DNMT* gene, was similarly constructed (Supplementary Table S1). All plasmids were sequenced at the University of Illinois Core Sequencing Facility.

### Media composition

<u>YPD</u> media was prepared with 1% yeast extract (VWR #90004-092), 2% peptone (VWR #90000-264) and 2% dextrose (VWR # BDH0230). <u>BMGY</u> medium contained 10 g /l (1% (w/v)) yeast extract, 20 g/ l (2% (w/v)) peptone, 100 mM potassium phosphate monobasic (VWR #MK710012), 100 mM potassium phosphate dibasic (VWR #97061-588), 4 × 10^−5^ % biotin (Life Technologies #B1595), 13.4 g/ l (1.34% (w/v)) Yeast Nitrogen Base (Sunrise Science #1501-500) and 2% glycerol (VWR #AA36646-K7).

### Plasmid transformation into *K. phaffii*

Competent cells were prepared by first growing one single colony of *K. phaffii* in 5 ml of YPD at 30 °C overnight. Fifty μL of the resulting culture were inoculated into 100 ml of YPD and grown at 30 °C overnight until they reached OD600 ∼1.3 - 1.5. The cells were then centrifuged at 1,500 *g* for 5 min at 4 °C and resuspended in 40 ml of ice-cold sterile water, centrifuged at 1,500 *g* for 5 min at 4 °C and resuspended with 20 ml of ice-cold sterile water, centrifuged at 1,500 *g* for 5 min at 4 °C and resuspended in 20 ml of ice-cold 1 M sorbitol, and centrifuged at 1,500 *g* for 5 min at 4 °C and resuspended in 0.5 ml of ice-cold 1 M sorbitol. Plasmids were linearized with the AvrII restriction enzyme and purified by ethanol precipitation. We mixed 80 μl of competent cells with 5-20 μg of linearized DNA and transferred them to an ice-cold 0.2 cm cuvette for 5 min before electroporation using exponential decay (Pulse parameters were 1500 V, 200 Ω and 25 μF). Immediately after pulsing, 1 ml of ice-cold 1 M sorbitol was added to the cuvette, and the cuvette content was transferred to a sterile culture tube containing 1 ml of 2x YPD. The culture tubes were incubated for 2 hours at 30 °C with shaking, and 50-100 μl of the culture was spread on YPDS plates (1% yeast extract, 2% peptone, 1 M sorbitol, 1% dextrose and 2% agar) with zeocin at 100 μg/ml. Control strains contained the control plasmid described above.

### PCR verification of DNMT and control plasmid integration

Integration of the plasmids at the correct target site of the *K. phaffii* genome was verified using PCR. Genomic DNA was isolated by resuspending single colonies in 20 mM NaOH and heating at 95°C for 10 minutes to lyse the cells. PCRs were performed using KAPA2G Robust PCR kits (KAPA Biosystems). A typical 25 μl reaction used 20–100 ng of genomic DNA, Buffer A (5 μl), Enhancer (5 μl), dNTPs (0.5 μl), 10μM forward primer (1.25 μl), 10 μM reverse primer (1.25 μl), KAPA2G Robust DNA Polymerase (0.5 U) and water (up to 25 μl). The DNA sequence of the primers for each target are provided in Supplementary Table S1: Sheet 2. The PCR products were visualized in 2% agarose gels, and images were captured using a ChemiDoc-It^2^ (UVP).

### Growth Conditions

Clones were inoculated into 5 ml YPD media and grown overnight at 30 °C. Five μl of the overnight cultures were transferred to BMGY. Five μl of the resulting cultures were transferred every day to 5 ml of fresh BMGY for 5 days, and at the end point, cells were collected for RNA or genomic DNA isolation. The time-course experiments followed the same procedure, except that cells were collected on days 1-4 after transferring to BMGY.

### Isolation of RNA and sequencing

Spheroblasts were obtained by incubating 20 million cells in buffer YT (1 M Sorbitol, 0.1M EDTA, pH 7.4) in the presence of 0.1% β-mercaptoethanol and 25 Units of Zymolase (Zymo Research). Spheroblasts were lysed by centrifugation in Qiashredder columns and total RNA was isolated using the RNEasy Mini Plus kit (Qiagen) according to the manufacturer’s instructions. RNA quality and concentration was assessed using a 2100 Bioanalyzer (Agilent). All paired-end RNA-seq experiments were performed by the BGI Americas Corporation using Illumina HiSeq 4000 at read length of 100 bp.

### Isolation of genomic DNA and bisulfite sequencing

Genomic DNA from 300 million cells was purified using the Master Pure Yeast DNA Purification kit (Epicentre) according to the manufacturer’s instructions. The resulting precipitated DNA was resuspended in Tris-EDTA (10 mM Tris-HCl pH 8.0, 1 mM EDTA) at ∼20 ng/μl. All paired-end whole-genome bisulfite sequencing (WGBS) experiments were performed by the BGI Americas Corporation using Illumina HiSeq X Ten at read length of 150bp.

### Data Accession

*K. phaffii* genome sequence and genome feature files were obtained from ftp://ftp.ncbi.nlm.nih.gov/genomes/all/GCA/001/708/085/GCA_001708085.1_ASM170808v1.

Sequences for knock-in genes DNMT3A1, DNMT3A2, DNMT1A, DNMT3B1, DNMT3L and for KanR and ZeoR (Supplementary Table S1) were added to the genome fasta file with corresponding entries added to the genome feature file. Sequences from MNase digestion-based nucleosome profiling performed by (19) were obtained from the Sequence Read Archive (https://www.ncbi.nlm.nih.gov/sra/) under the accession number SRP031651 and used to calculate nucleosome occupancy and dyad positions (Supplementary Methods: “MNase-seq data processing” and “Calculation of nucleosome occupancy and dyad calling”).

### Quantification of gene expression

Paired end RNA-seq reads were mapped to the *K. phaffii* genome using TopHat2 and condition-specific reference genomes to prevent multimapping to highly similar DNMT isoforms (20) (version 2.1.1) (Supplementary Methods: “RNA-seq analysis”). Mapped reads were filtered with samtools view (version 1.7) with options -h -f 3 -F 3596 -q 13, and read pairs mapping to different chromosomes were removed (21). Gene expression values, in units of FPKM (fragments per kilobase of transcript per million mapped reads), were calculated using cuffnorm (version 2.2.1) (Figure S1) (22). Alignment rates and sequencing depths for each experiment are given in Table S2.

### Differential expression

We tested for differential gene expression between conditions using RNA-seq read count tables. These tables were generated from filtered bam/sam files with HTSeq-count (version 0.9.1) using options -s no -I ID -t gene (23). Differential expression analysis used DEseq2 (version 1.14.1) (24). The DEseq2 model was fit on data from all read count tables using covariates for each of our 13 experimental conditions, as well as for the three batches in which RNA-seq experiments were performed. After fitting, we tested for differential expression of genes in each knock-in condition relative to cells containing a control plasmid, with independent filtering parameter (alpha) 0.05 and all other options set to default. We used a threshold of 0.05 for Benjamini-Hochberg adjusted p-values reported by DEseq2 to identify differentially expressed genes.

### Quantification of DNA methylation

We used Bismark (version 0.18.0) to quantify methylation at cytosines genome wide (25). Supplementary Methods: “WGBS analysis” details our method which yielded files describing the observed methylation status of CpG for each read and report files which contained counts of methylated and unmethylated observations pooled across reads covering a single CpG. For each experiment, alignment rates, coverage, and bisulfite conversion rates based of lambda phage DNA are given in Table S3.

The CpG report files were used for checking reproducibility among WGBS experiments and for calculation of mCpG rates using Empirical Bayes (Figure S1A, Supplementary Methods: “Clustering experiments by window average methylation” and “Empirical Bayes approach for estimating mCpG rate”). We summarized methylation patterns across genomic regions aligned at features of interest (e.g. TSS) by calculating sliding widow averages of mCpG rates and estimated credible intervals with a bootstrap approach (Supplementary Methods: “Calculation of feature-aligned mCpG rates”).

### GO and pathway analysis

We used the R package RDAVIDWebService to programmatically perform DAVID GO analysis on sets of DE up and DE down genes relative to control (26, 27). Details are given in Supplementary Methods: “Gene ontology analysis and GO term clustering.” We used the R package Pathview to visualize the time-course differential expression of genes composing KEGG pathways (28).

### Predicting differential expression from metagene methylation

Logistic regression classifiers for the “DE down vs. rest” and “DE up vs. rest” tasks were evaluated in a 10-fold cross validation scheme. Gene differential expression status, metagene methylation and time-course day information were pooled across time-course days, and the pooled data was randomly split into 10-folds. For each gene, model inputs were a one-hot encoding of time-course day and tanh transformed standardized metagene methylation data. Standardization parameters were calculated separately for each time-course day by applying a sliding widow to metagene mCpG data in the training set. The tanh transform served to reduce the effect of outlier methylation rates. Target outputs were 0,1 encodings of class labels. Details of dataset construction are given in Supplementary Methods: “Description of logistic regression task and input.”

We wanted our classification model to avoid a binning approach that imposed broad metagene intervals over which the effect of CpG methylation was assumed constant. Our logistic regression model therefore included 30 regression coefficients, each of which multiplied standardized methylation inputs in one of the 30 bins that finely partitioned the metagene interval used for prediction. To prevent overfitting, we penalized the square of the second derivative of these regression coefficients with respect to metagene position. This penalty introduced a regularization parameter whose value was chosen based on cross validation performance (Figure S2-S4, Supplementary Methods: “Logistic regression model” and “Choice of logistic regression smoothness penalty”)

### CNN training and interpretation

The dataset used for CNN training and evaluation was constructed from cytosines in the CpG context, lying in the metagene interval [-0.25,1.25] for at least one of the annotated *K. phaffii* genes. When a cytosine corresponded to more than one metagene interval, it was assigned to the gene containing the cytosine between its transcription start and termination sites and then, if this did not resolve the assignment, to the gene with nearest TSS in absolute genomic coordinates. This full data set contained 510,698 cytosines which were split into training, validation and test sets as follows: to insure balance of highly and lowly methylated cytosines in each set, we divided the full data set into quintiles according to maximum likelihood estimates of methylation rate. Pairs of cytosines corresponding to a single CpG site were then assigned to one of 25 groups according to the quintile of methylation rates on the plus and minus strand. For each group, we randomly assigned 80%, 10%, and 10% of the elements to the training, validation and test sets, respectively. CNN’s were trained using the python packages Keras https://keras.io. The encoding of CNN inputs and details of CNN training and architecture are given in Supplementary Methods: “Description of CNN inputs” and “Architecture and training of CNN”.

The simulated annealing (SA) interpretation method sampled sequences that approximately maximize or minimize the pre-activation of output neurons of the CNNs trained on each condition. SA was initialized at a condition-specific initial temperature and was cooled for 8 × 10^6^ iterations using a logarithmic cooling schedule with a set of 2 × 10^4^ sequences collected for each initialization. A detailed description of the algorithm and choice of initial temperatures is given in Supplementary Methods: “Simulated annealing” and “Choice of the temperature parameter *d* in simulated annealing.” Validation of SA motifs constructed from sequences in clustered sets of SA samples used FIMO to scan the *K. phaffii* genome, as described in greater detail in Supplementary Methods: “Validation of SA motifs” (29).

## RESULTS

### Exogenously expressed DNMT3A1, 3A2 and 3B1 methylate the *K. phaffii* genome

To study DNA methylation in *K. phaffii* which lacks endogenous DNMTs, we created clones with a single knock-in of genes encoding human DNMT enzymes, as well as clones with a double knock-in of these genes with *DNMT3L*, which has been shown to stimulate the function of other DNMTs in mammals (30) (Figure 1A, Methods). For all single-knock-in clones and the double-knock-in clones containing *DNMT3A1* (3A1-3L), *DNTM3A2* (3A2-3L), and *DNMT1A* (1A-3L), we measured the genome-wide levels of 5mC and gene expression five days after generating and validating the knock-in strains using duplicate whole-genome bisulfite sequencing (WGBS) and triplicate RNA-seq experiments, respectively (Methods). Because *de novo* methylation by murine DNMT3B1 has previously been studied in the yeast *S. cerevisiae*, where no significant transcriptional changes could be attributed to DNA methylation, and because DNMT3B is often aberrantly overexpressed in human cancers (31, 32), we chose to study this particular *de novo* DNMT in greater detail in *K. phaffii*. We thus performed a time-course experiment for the *DNMT3B1* and *DNMT3L* double-knock-in clones (3B1-3L), measuring 5mC and gene expression on each of the four days following validation of knock-in expression (3B1-3L d1 through 3B1-3L d4). In each condition, RNA-seq confirmed the expected patterns of expression for knock-in genes and for the selection genes KanR and ZeoR (Figure 1B, Figure S5A,B, Supplementary Methods: “DNMT Western blot”).

**Figure 1:**
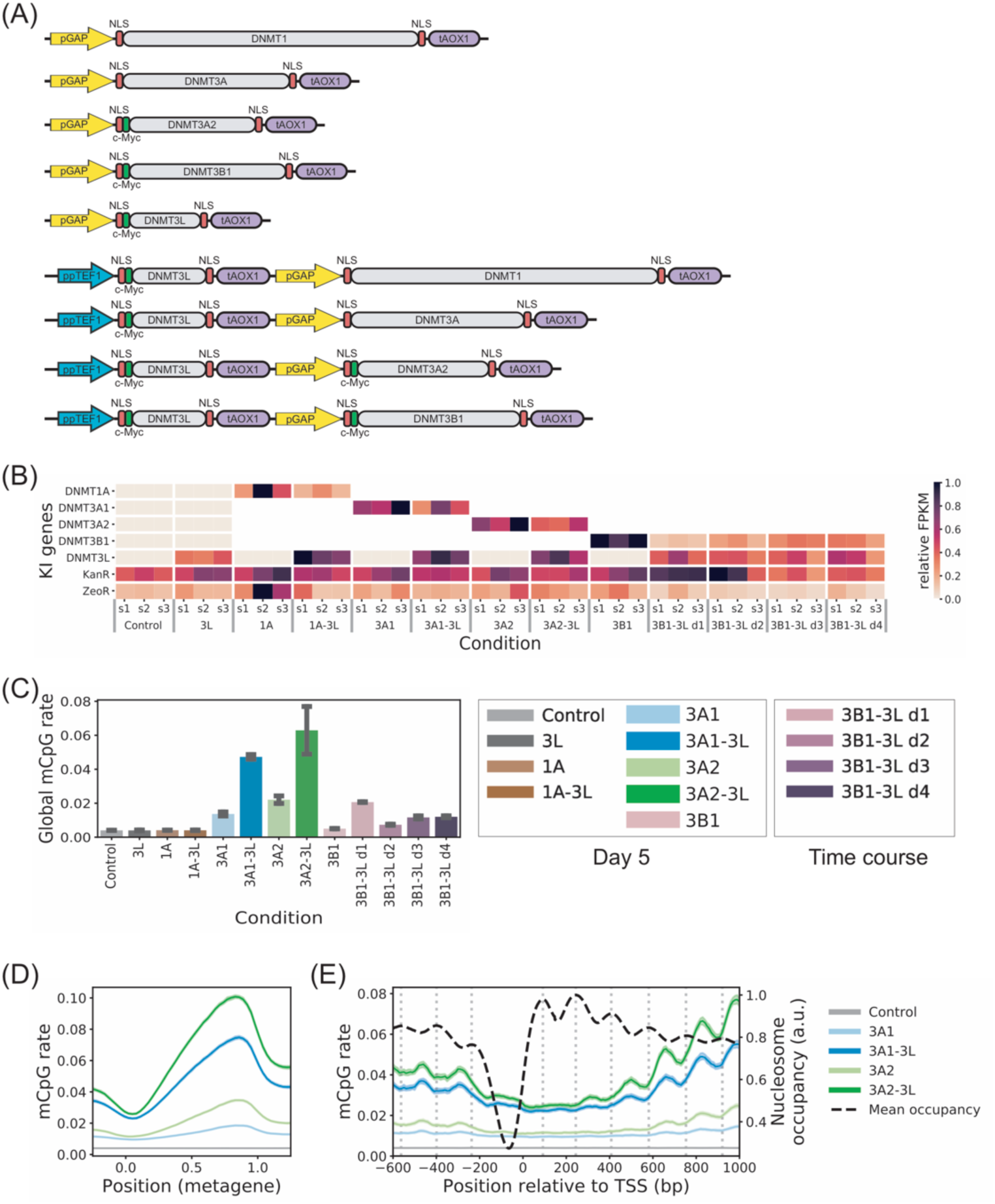
Combinatorial knock-in of DNMT genes methylates the *K. phaffii* genome. **(A)** Schematic representation of expression cassettes for knock-in of DNMT genes. **(B)** Expression levels (FPKM) of knocked-in genes in each experiment scaled by maximum knock-in expression in each row. White cells in the matrix indicate that a particular DNMT isoform was excluded from alignment to prevent multi-mapping (Methods). **(C)** Genome-wide 5mC fraction in CpG context averaged over two replicates. Error bars show results for the minimum and maximum replicate. **(D)** Metagene plot of mCpG rates averaged across *K. phaffii* genes. CpG-context cytosines were assigned to a metagene coordinate, with the distance downstream of the TSS scaled by gene length. Solid lines indicate average mCpG rate of cytosines in sliding metagene coordinate windows. Shading indicates 95% Bayesian credible interval (Methods). **(E)** Profiles of mCpG rates and nucleosome occupancy aligned at TSS and averaged across *K. phaffii* genes. Lines and shaded credible intervals were calculated using a sliding window method similar to (D).

To begin our analysis, we compared the global 5mC levels among experimental conditions as well as among sequence contexts, and observed several patterns of methylation similar to those in mammals (4, 33). First, only the DNMTs capable of *de novo* methylation in humans — 3A1, 3A2 and 3B1 — produced 5mC levels substantially greater than control (Figure 1C, Table S3). Second, this *de novo* methylation was largely restricted to the CpG dinucleotide context, with methylation in other sequence contexts comparable to or only slightly greater than control (Figure S5C,D). Third, the catalytically inactive DNMT3L alone did not produce 5mC levels above control, but co-expression of 3L with any of the *de novo* DNMTs increased global CpG methylation by 1-3 fold compared to the corresponding single knock-in. Based on these observations, we chose to focus our subsequent analysis primarily on CpG methylation by single and double knock-in of the established *de novo* DNMTs. CpG methylation patterns for these active DNMT conditions were highly reproducible among replicates (Figure S1A); we therefore pooled the raw methylated and unmethylated CpG count data across replicates and estimated site-specific methylation rates using the Bayes posterior mean calculated from the count data and an empirical Bayes beta prior (Methods). Below, we refer to these final methylation rate estimates as mCpG rates.

The variation of mCpG rate along genes agreed with the established methylation profile of mammalian genes (4), where the mCpG rate consistently dipped near the transcription start site (TSS) and then increased along the gene body to peak around 80% of the gene length (Figure 1D, Figure S6A). Moreover, near the TSS, mCpG rate was anti-phased with statistically positioned nucleosome occupancy (34) (Figure 1E, Figure S6B). This combination of depleted methylation and anti-phasing with nucleosome occupancy at TSS agreed with the previous observations in *S. cerevisiae* where murine *DNMT3B1* was knocked in (17). Furthermore, anti-phasing of methylation and nucleosome occupancy was also reported at well-positioned nucleosomes adjacent to CTCF binding sites in mouse embryonic stem (ES) cells (16), suggesting that local chromatin structure may influence DNMT activities. Overall, these results indicated that exogenously expressed DNMT3A1, 3A2 and 3B1 can methylate their CpG targets, function synergistically with DNMT3L and produce patterns of gene methylation resembling those in mammals in the absence of co-evolved cellular machinery.

### Exogenous expression of DNMT yields broad transcriptional changes that decay with time

We investigated transcriptional changes following DNMT knock-in both at the broad level of covarying gene sets and at the granular level of differential gene expression in each condition. At the broad level, principal components analysis (PCA) identified the first principal component (PC1) capturing the time-course progression of transcriptomic states, starting from an out-group on day 1 and progressing monotonically towards the bulk of the remaining samples (Figure 2A, Supplementary Methods: “Gene expression PCA”). PC1 explained nearly 50% of the observed variance across experimental conditions, showing that the difference between early days (time-course days 1 and 2) and later days – reflecting the immediate and prolonged effects of methylation stress, respectively – was the dominant structure in our expression data (Figure S7A).

**Figure 2:**
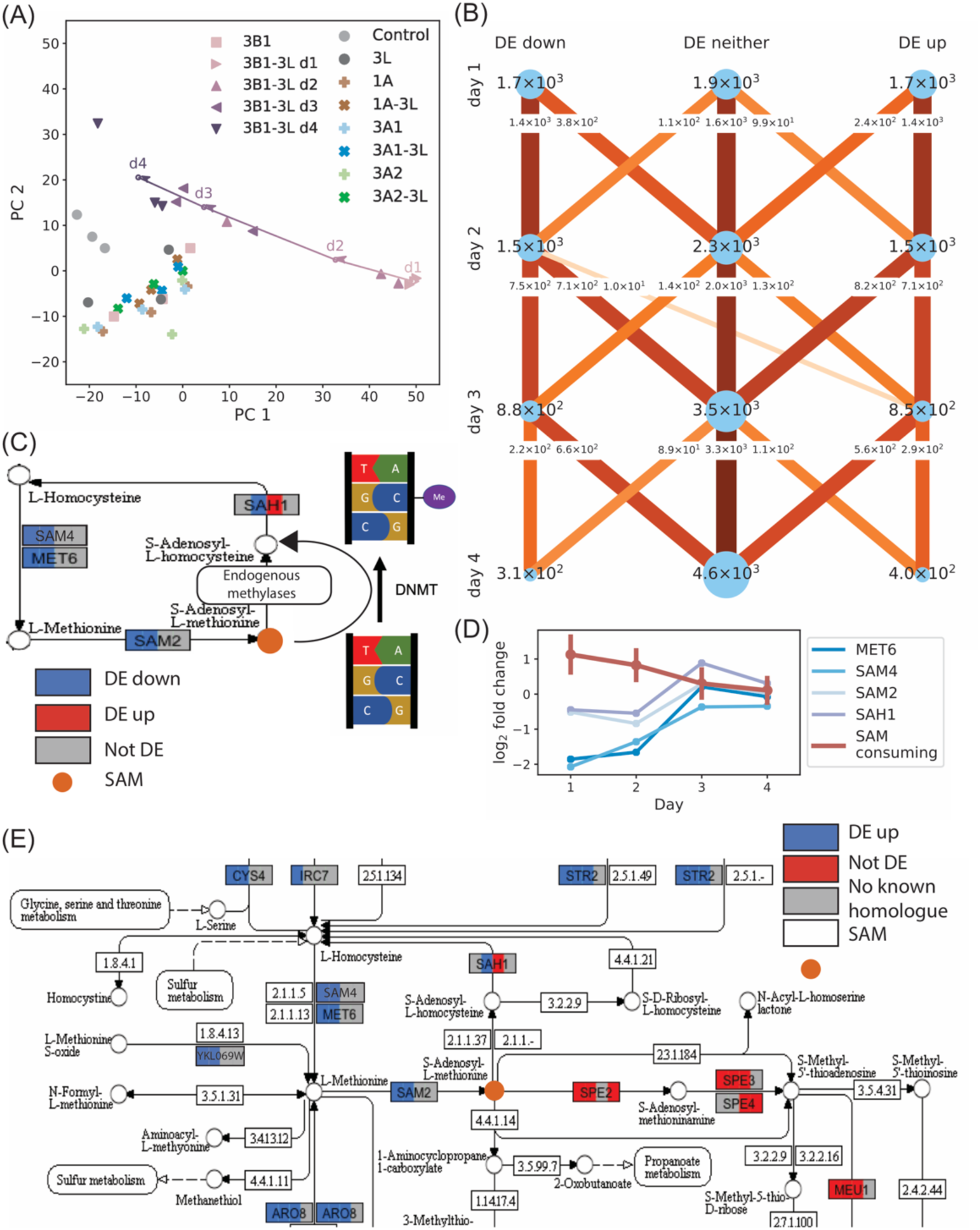
DNMT expression causes broad transcriptional changes including differential expression patterns that reduce SAM availability in the 3B1-3L condition. **(A)** Principal component analysis of transcriptome-wide mRNA levels. Open circles and arrows, labelled d1 through d4, indicate the mean of triplicates for each time-course day. **(B)** Numbers of genes differentially expressed in time course compared to control. Node labels and size indicate the number of differentially expressed genes on each day after DNMT induction. Edge labels and color indicate the number of genes transitioning between differential expression states. **(C)** Time-course differential expression status of methionine cycle genes relative to control. **(D)** Log_2_ fold-change of methionine cycle genes and SAM-consuming genes in the 3B1-3L condition relative to control. SAM-consuming genes were defined as the union of genes belonging to co-clustered GO terms related to SAM function and the set of spermine synthesis genes SPE3, SEP4, MEU1. Error bars show standard deviation over a total of 59 genes. **(E)** Time-course differential expression status of methionine cycle, spermine synthesis and related genes.

Analysis at the level of differentially expressed (DE) genes further supported the findings of PCA (Table S4). While all knock-in conditions, in general, had many DE genes relative to control (median # “DE up” genes = 442, median # “DE down” genes = 434), the number of DE genes for early days of the time course (more than 3,000 of the 5,273 annotated genes in *K. phaffii* on days 1 and 2) far exceeded the number of DE genes for later days and other knock-in conditions. Furthermore, the number of DE genes decreased monotonically with time (Figure 2B, Figure S7B). The preponderant flow of genes from DE to non-DE states during the time course and the progression of time-course expression along PC1 could be interpreted as resulting from a combination of two contributions: (1) cells may modify gene expression via endogenous regulatory mechanisms to produce a coordinated initial response to the stress of DNMT expression and DNA methylation; (2) DNA methylation itself may act on individual genes to alter their transcriptional output via changes in chromatin or the transcription process. We will computationally characterize these contributions in the subsequent sections.

Transcriptome-based hierarchical clustering of samples showed some mixing of conditions (Figure S1B), although the control samples were still an outgroup. To better visualize the clustering pattern, we performed hierarchical clustering of control, 3L and methylating double knock-in conditions only (Figure S1C) and found a structure separating 3B1-3L early days, 3B1-3L later days, 3A-3L, and control conditions into subclusters. We reasoned that part of the transcriptional similarity among knock-in conditions may be explained by a common, generic response to heterologous expression of human proteins. In support of this hypothesis, the set of 179 “DE up” genes common to all non-methylating conditions was enriched for gene ontology (GO) terms related to the ribosome and protein translation (Figure S8A, Table S5), consistent with the fact that the increased translational burden of expressing a heterologous protein can elevate the transcription of ribosome-related genes in *K. phaffii* (35). Moreover, this set of common “DE up” genes showed a high degree of overlap with “DE up” genes in the *de novo* methylating conditions, demonstrating the generality of the ribosome-related transcriptional response across conditions (Figure S8B,C). Similarly, the set of 129 “DE down” genes common to all non-methylating conditions showed a high degree of overlap with “DE down” genes in the *de novo* methylating conditions (Figure S8). This set was enriched for genes functioning in mitotic sister chromatid segregation and localizing to the spindle pole body (Table S5). For both common “DE up” and “DE down” sets, the overlap with 3B1-3L DE genes decreased with time-course progression (Figure S8C).

Overall, the agreement of enriched GO terms with previously reported response to heterologous expression and the consistent differential expression pattern across conditions indicated that modulation of these ∼300 common genes was likely a transcriptional response to the stress of heterologous protein expression. However, we did not observe consistent upregulation of signature genes associated with heat shock proteins and protein processing in the endoplasmic reticulum known to be triggered by protein misfolding and aggregation (36) (Figure S9).

### Cells adapt to DNMT3B1-3L-induced DNA methylation stress by modulating the availability of SAM

To disentangle expression changes attributable to active cellular response from those attributable to biochemical/biophysical changes in chromatin, we reasoned that molecular adaptation to exogenous methylation stress should involve coordinated gene sets participating in common pathways or biological processes. We therefore applied GO analysis to sets of “DE up” and “DE down” genes in each condition (Supplementary Table S4, and Supplementary Table S5) (27). Focusing on our 3B1-3L time-course data, we clustered the enriched GO terms based on their overlap of DE genes (Figure S10, Supplementary Methods: “Gene ontology analysis and GO term clustering”). Two of these clusters contained terms related to the production and consumption of S-adenosyl methionine (SAM), the essential methyl donor to DNMT for 5mC DNA methylation and to other methyltransferases for many endogenous methylation processes (37).

The first cluster contained terms related to metabolic pathways and cellular respiration. Among the enriched pathways was the cysteine and methionine metabolic pathway that included the methionine cycle. In this cycle, methionine is converted to SAM which acts as a methyl donor for many endogenous metabolic reactions and, in our engineered cells, for DNA methylation catalyzed by the exogenous DNMTs (37). Methyl group donation converts SAM to S-adenosyl-l-homocysteine (SAH) which is, in turn, converted to L-homocysteine and then back to methionine, completing the cycle. Examining the genes encoding enzymes for SAM synthesis and recharge of methionine from SAH, we found that almost all were significantly suppressed during the first two days of our time-course experiment (Figure 2C), implying that the cells were tuning their expression program to counter the DNMT methylation activities. Within this gene set, the significantly decreased expression of SAM4, MET6, and SAM2 on day 1 and 2 would likely slow down the synthesis of SAM and its precursor methionine, thus decreasing the availability of the methyl-donor SAM. Similarly, the decreased expression of SAH1 on day 2 should impede the conversion of SAH to L-homocysteine, thereby increasing the concentration of SAH, a known potent inhibitor of both DNMTs and histone methyltransferases (38). Interestingly, while differential expression of other methionine cycle genes ceased after two days post DNMT induction, *SAH1* was one of only 10 genes to transition from being “DE down” on day 2 to “DE up” on day 3, potentially to relieve the inhibition of histone methyltransferases imposed by the accumulation of SAH. The paucity of genes that transitioned between these DE states in our time-course (Figure 2B) further supported the idea that changes in SAH1 expression resulted from an adaptive gene regulatory network response to methylation stress, rather than as a direct consequence of methylating the *SAH1* gene itself. These findings together supported that the early suppression of genes catalyzing the methionine cycle was an active cellular response to reduce the stress of *de novo* DNA methylation by depriving DNMTs of the methyl-donor SAM.

The second cluster contained GO terms that grouped genes with endogenous SAM consuming functions. Genes belonging to this cluster included spermine synthesis genes SPE2, SPE3, SEP4 and MEU1, histone methyltransferases SET1, SET5, DOT1, and NOP1, ribosomal methyltransferases RKM3 and RKM4, and a variety of tRNA methyltransferases (Supplementary Table S5). These genes were upregulated in the time-course relative to control, with the most dramatic expression increases occurring on days 1 and 2, coincident with the aforementioned period of down-regulation of methionine cycle genes (Figure 2D). The simultaneous and opposing regulation of methionine cycle and SAM consuming genes likely functioned to alter the output of metabolic pathways to decrease SAM production while increasing SAM consumption through endogenous processes. The increased expression of endogenous histone and ribosomal methyltransferases might also suggest a way of compensating for the decreased availability of SAM and outcompeting the DNMTs. Figure 2E illustrates this dual modulation in a network view of cysteine and methionine metabolism (39) (see Figure S11 for differential expression status of all genes in KEGG pathway). The magnitude of this transcriptional response was greatest on the first two days post DNMT induction, which could explain why the minimum global methylation level occurred on day 2. Using 3L as reference in our differential expression analysis did not change our main conclusions (Figure S12).

Our time-course measurement of SAM levels showing the lowest concentration on day 2 was consistent with our transcriptomic observation (Figure S13, Supplementary Methods: “SAM measurements”). However, changes in the expression levels of metabolic enzymes may not necessarily produce simple flux changes in metabolic pathways. A more thorough analysis involving metabolomics approaches would be thus needed in the future to resolve how the expression changes described above modify the dynamic outputs of the metabolic network regulating SAM/SAH levels.

Finally, although the proposed cellular response involved only a subset of the approximately 3000 genes differentially expressed on days 1 and 2, other examples of coordinated transcriptional change could also be attributed to modulating SAM levels. For example, the gene SLC25A26 encodes the only known SAM mitochondrial transporter in humans, mutation of which could lead to a mitochondria defect arising from SAM deficiency (40); in our study, the *K. phaffii* homologue PET8 was upregulated on days 1-3 (and in conditions 1A and 1A-3L), and genes related to mitochondrial function were frequently elevated throughout the time-course (Supplemental Table S5) (41). These expression patterns might reflect efforts to raise mitochondrial SAM levels and protect the mitochondria from a potential damage triggered by the SAM reduction during early stress response.

### DNA methylation is a significant predictor of expression status that improves with time post DNMT3B1-3L induction

We next sought to determine whether specific patterns of CpG methylation were associated with differential transcription of individual genes. A recent study that exogenously expressed murine DNMT3B1 in *S. cerevisiae* observed no association between gene methylation and changes in expression and attributed this finding to the absence of proteins capable of recognizing and mediating 5mC effects in yeast (17). In contrast to this result, however, by aggregating mCpG rates across “DE up”, “DE down” and non-differentially expressed (“No DE”) genes relative to control, we observed distinct methylation patterns for each gene set that were consistent across the time course (Figure 3A) and across all active DNMT conditions (Figure S14, Supplementary Methods: “Calculation of feature-aligned mCpG rates”). In particular, “DE down” genes were more highly methylated than “No DE” genes, which were, in turn, more highly methylated than “DE up” genes. The only exception to this pattern was in the lowly methylated 3B1 condition, where the methylation levels of “DE up” and “No DE” genes were comparable. Moreover, our time-course data demonstrated that the distinction between methylation patterns of “DE down” genes and those of “No DE” or “DE up” genes increased with time.

**Figure 3:**
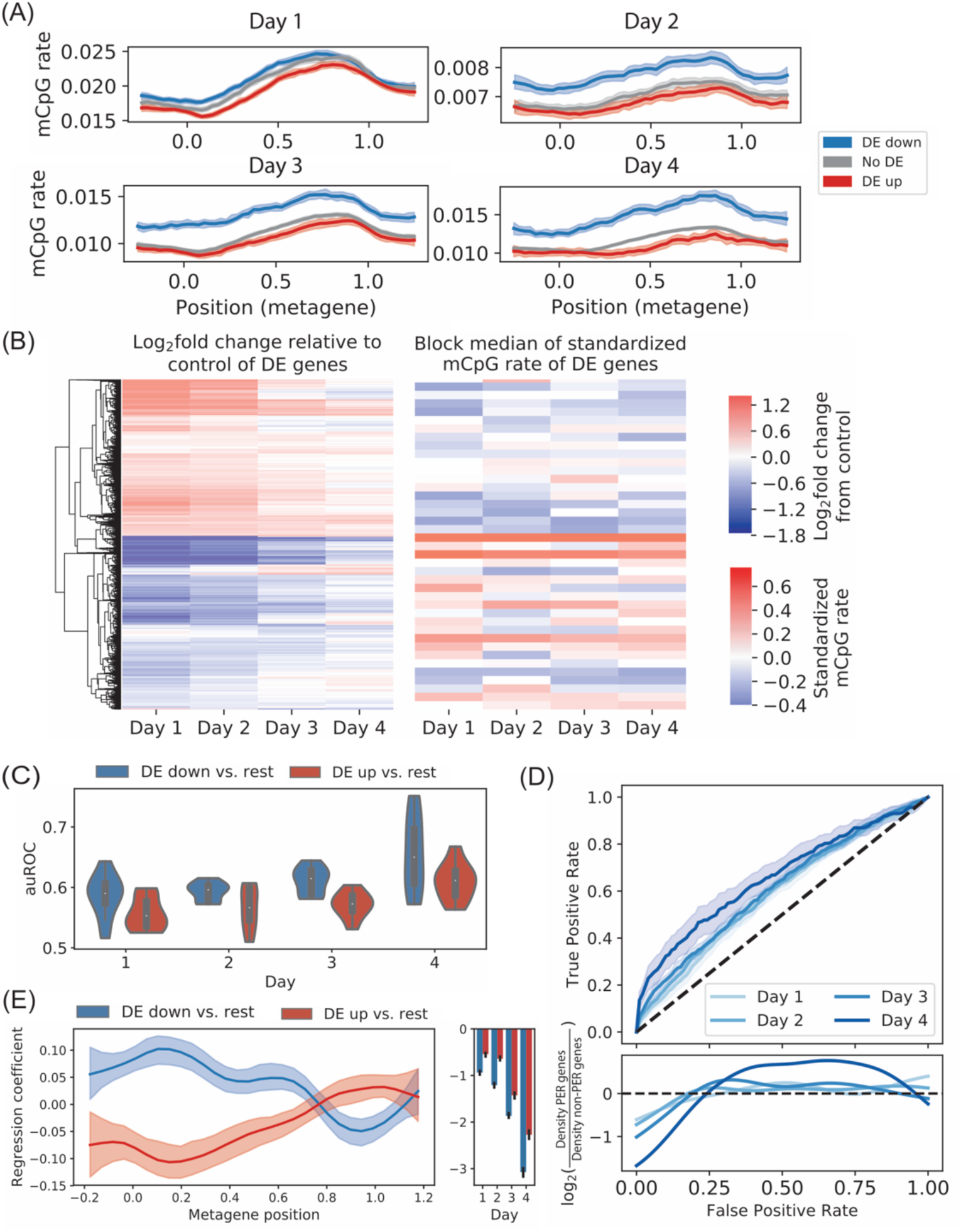
Changes in expression are inversely related to gene methylation by DNMT3B1-3L throughout time course. **(A)** Metagene plot of mCpG rates averaged across “DE down,” “no DE” and “DE up” genes on each day. (**B**) Clustering of differentially expressed genes by log_2_ fold-change in expression relative to control (left panel) and heat map of average gene mCpG rate in metagene interval [-0.2, 0.4] (right panel). The right panel heat map is organized by expression-based clustering, shows medians over blocks of 100 genes and is standardized on each day. **(C)** Distribution of auROC across 10 CV folds for classification of test data belonging to each time-course day. **(D)** ROC for classification of test set data in “DE down vs. rest” task (top) and log ratio of distributions of minimum false positive rates for correct classification of PER and non-PER “DE down” genes (bottom). Solid lines in ROC indicate mean true positive rate for given false positive rate taken across CV folds; shading indicates 95% confidence interval. **(E)** Regression coefficients for logistic regression classifiers. Solid lines and bar heights indicate parameters learned from all time-course data. Shading indicates 95% confidence interval estimated by bootstrap resampling of time-course data (see also Figure S16).

To gain more insight into the association between gene methylation and differential expression, we focused on our time-course data that could capture the dynamics of epigenetic and transcriptomic changes. We clustered DE genes by their log-fold changes relative to control across four days and used this clustering to organize heat maps of time-course expression and methylation (Figure 3B, Supplementary Methods: “Clustering gene expression changes and methylation patterns”). The paired heatmaps qualitatively confirmed the general inverse relationship between expression change and gene methylation. Figure S15 shows examples of this inverse relationship at select gene loci and a scatter plot of global gene expression against methylation rate demonstrating the inverse trend. To quantify the association, we built a pair of logistic regression classifiers and assessed their performance. Specifically, we considered two classification tasks that predicted the differential expression status of each gene from time-course day and the metagene methylation pattern. For the first task, termed “DE down vs. rest,” the classifier aimed to distinguish “DE down” genes from genes that were either “DE up” or “No DE” on a given day. For the second task, termed “DE up vs. rest,” the classifier aimed to distinguish “DE up” genes from genes that were either “DE down” or “No DE” on a given day (Methods).

By measuring performance statistics for 10-fold cross validation (CV) on each day separately, we made several observations that supported an association between DNA methylation and differential expression: first, for each CV fold on each day, area under the receiver operating characteristic (auROC) was greater than 0.5, the expected value for a null model, indicating that methylation was a statistically significant predictor of differential expression status throughout the time course (Figure 3C). Second, the performance on the “DE down vs. rest” task was uniformly better than that on the “DE up vs. rest” task, in terms of both auROC (Figure 3C) and area under the precision recall curve (auPRC) (Figure S16A), implying that methylation profiles were more informative for predicting decreases in expression than increases in expression. Third, classifier predictions generally improved with the number of days following DNMT induction as initial secondary effects were suppressed; this trend was apparent for both classifiers when performance was measured using auROC and for the “DE down vs. rest” classifier when performance was measured with auPRC. Finally, our logistic regression model continued to predict changes in gene expression measured relative to 3L condition (Figure S12C).

Overall, the pattern of gene methylation by DNMT3B1-3L was a moderate predictor of decreased expression. For example, on day 4 the “DE down vs. rest” classifier could identify 30.0% of the DE down genes at a false positive rate of approximately 9.0%, implying that the ratio of true positives to false positives, for this decision threshold, was 3.3 times that expected by chance (Figure 3D). By contrast, the pattern of gene methylation by DNMT3B1-3L was only a weak predictor of increased expression. For example, on day 4, identifying 30.0% of the up-regulated genes with the “DE up vs. rest” classifier required a false positive rate of approximately 23.0%, implying that the ratio of true positives to false positives, for this decision threshold, was only 1.3 times that expected by chance (Figure S16B). These findings supported the conventional understanding that DNA methylation is generally associated with transcriptional repression.

Finally, we examined the regression coefficients learned by our classifiers. The coefficients for day labels were negative and increased in magnitude as time progressed, reflecting the steady decrease in the number of differentially expressed genes as a function of time (Figure 3E). The regression coefficients for standardized mCpG rates from the two classifiers showed similar variations in magnitude with metagene position, but had opposite signs (Figure 3E), confirming the inverse relationship between gene methylation and transcription change. The magnitude of regression coefficients peaked just downstream of the gene TSS, indicating that changes in methylation in this region were most predictive of differential expression.

### Repression of pathway-enriched gene sets is poorly explained by DNMT3B1-3L-induced DNA methylation

Previous sections have separately considered endogenous gene regulation in dynamic stress response and changes in transcription associated with gene body methylation. To investigate the relationship between these two modes of gene regulation, we compared the performance of our “DE down vs. rest” classifier on putative endogenously regulated (PER) genes and non-PER genes defined as follows: at each time-course day, the PER set consisted of genes suppressed relative to control and annotated as participating in either a KEGG pathway or GO biological process that contained a significant number of “DE down” genes on that day. Conversely, the non-PER set on each time-course day consisted of transcriptionally supressed genes not in the PER set on that day.

Given that endogenous regulatory mechanisms ordinarily operate in the absence of DNA methylation, we hypothesized that PER genes were likely to be co-regulated by pathway networks and thus that gene methylation would be less predictive of transcriptional suppression of these genes than that of non-PER genes. To test this hypothesis, we calculated the minimum false positive rates (FPRs) at which each of the PER and non-PER genes was correctly identified as DE down by the “DE down vs. rest” classifier. For each gene, having lower minimum FPR implied that the gene’s differential expression status was more easily predicted by its methylation pattern. Comparing the empirical distributions of minimum FPRs for the two gene sets showed that PER genes were depleted at low FPR relative to non-PER genes (Figure 3D bottom, Supplementary Methods: “Density estimation for minimum FPRs”). This trend supported our hypothesis and was statistically significant on days 1 and 4 (*p* = 3.4 × 10^−2^, and 1.6 × 10^−2^, respectively; two-sided Mann-Whitney U test with Bonferroni correction). Overall, this result indicated that suppressed genes not part of any coordinately down-regulated pathway tended to have higher gene methylation which made their transcriptional suppression easier to predict, compared to PER genes.

In this analysis, we assumed that collective changes in the transcription of functionally related genes imply true endogenous regulatory mechanisms; however, an alternative explanation for such collective expression changes may be the hypermethylation of a common master regulator or changes in a common chromatin environment containing multiple functionally related genes.

We thus examined the potential role of common chromatin environment and gene methylation in suppressing the aforementioned methionine cycle-associated genes. Of the four methionine cycle genes (*SAM2, SAM4, MET6*, and *SAH1*), only *SAM2* and *SAH1* occur on the same chromosome — roughly 23kb apart — ruling out co-regulation via changes in a single common chromatin environment. To assess the likelihood that methylation at these individual genes might account for their collective suppression, we assumed the null hypothesis that the probability of each methionine cycle gene being DE down could be predicted by the “DE down vs. rest” classifier solely based on gene methylation patterns. The p-values for observing 3 of the 4 methionine cycle genes to be “DE down” on day 1 and all 4 of these genes to be “DE down” on day 2 were 0.09 and 0.006, respectively, indicating that the collective down regulation of these genes was not likely explained by their gene methylation patterns alone. These results supported that individual gene hyper-methylation or similarity of chromatin environments do not fully explain the collective suppression of the methionine cycle-associated genes. However, we cannot rule out the possibility that collective changes in pathway-enriched genes may be due to a common master regulator epigenetically modulated by DNMT activity.

Finally, we considered the relationship between gene methylation and endogenous regulation for “DE up” genes. Defining PER and non-PER gene sets as above for “DE up” genes, we found that our “DE up vs. rest” classifier could more easily predict the up-regulation of PER genes than that of non-PER genes using gene methylation patterns. This result is illustrated by the significant enrichment of PER genes at low FPR values for all time course days (Figure S16B, *p* = 1.5 × 10^−3^, 3.7 × 10^−2^, 3.5 × 10^−5^, and 4.2 × 10^−2^ for days 1-4, respectively; two-sided Mann Whitney U test with Bonferroni correction). Biologically, this finding implied that genes upregulated as part of broader changes in pathways were protected from methylation in the 3B1-3L condition.

### DNMT3L linearly increases genome-wide methylation level and DNMT3B1 methylates near centromeres and telomeres

To investigate the role of DNMT3L, we examined the scatter plots of average methylation rates in 10kb bins between single knock-in and corresponding double knock-in conditions (Figure S17A) and observed high linear correlation (r=0.83 for 3A1 vs. 3A1-3L and r=0.92 for 3A2 vs. 3A2-3L). For the case of 3B1 vs. 3B1-3L, the correlation was lower (r=0.69), likely because of higher statistical fluctuation in the genome-wide low methylation rate by DNMT3B1 alone. This analysis implied that DNMT3L amplifies the overall methylation level genome wide, without noticeable changes in genomic distribution.

To study potential differences in methylation activity between DNMT3A and DNMT3B, we computed the Spearman correlation of smoothed methylation patterns across the genome and found the correlation to be generally high among 3A/3A-3L family members (r∈[0.83, 1.00]), but slightly lower between 3A/3A-3L and 3B1-3L d4 (r∈[0.69, 0.79]). To find out what is causing the difference between 3B-3L-d4 and 3A family, we made scatter plots of ranked average methylation rate in 10kb genomic bins (Figure S17B); while most bins were scattered along the diagonal, there were eight outlier bins, corresponding to sites that were comparatively lowly methylated by 3A1-3L and 3A2-3L, but highly methylated by 3B1-3L. Strikingly, every outlier bin was located near a centromere or telomere, as confirmed by a local correlation analysis (Figure S17C). In terms of sequence characteristics, these regions had high CpG ratio. Our finding is consistent with the fact that DNMT3B interacts with the centromere-specific histone variant CENP-C in mammals (42).

### Convolutional neural networks learn distinct sequence preferences of DNMT3 family members

We next sought to study DNA sequence determinants of CpG methylation specific to each DNMT3 family member. To identify fundamental sequence features characterizing preferentially methylated CpGs, six convolutional neural networks (CNNs) of the same architecture were trained for the six experimental conditions: 3A1, 3A2, 3A1-3L, 3A2-3L, 3B1, 3B1-3L d1. Each network predicted the methylation rate of a cytosine in the CpG context within or proximal to a gene (Methods). Input to the CNNs consisted of a 201 bp-long DNA sequence, centered around the C of interest, together with the metagene position of each nucleotide. We included the information about metagene position in order to quantify and separate out the effect of CpG location, which might be confounded by inhomogeneous H3K36me3 along the gene body, from that of sequence content on methylation (Figure 1D). It was previously shown that the N-terminus of histone 3, in particular unmethylated H3K4, is required for *de novo* DNA methylation in *S. cerevisiae* (43) and that DNA methylation spatially correlates with H3K36me3 in several species including *S. cerevisiae* (17). Since H3K4me3 and H3K36me3 are biased towards the transcription start and termination sites, respectively, our model implicitly accounted for the association between these histone modifications and DNA methylation by using the metagene coordinate as a predictor variable. The sequence length of 201 bps was chosen based on the length scale of local chromatin encompassing one nucleosome and linker DNA.

We assessed the performance of the CNNs in two complementary ways: first, the CNNs accurately reproduced the observed test-set mCpG rates across metagene positions for all six knock-in conditions (Figure 4A), confirming that the models were indeed utilizing the information about metagene position. Second, to assess whether the CNNs also utilized the sequence information to predict methylation rates, we permuted the nucleotides flanking CpG while fixing the metagene position for each test-set input and then compared the CNN loss for each original input sequence to losses for the corresponding permuted sequences. Defining informative sequences as those with loss in the top 10% of the distribution of permuted loss, we found that the fraction of CpG sites with informative flanking sequence information was significant in all conditions (*p* < 4.1 × 10^−105^ for all conditions, Binomial test) (Figure 4B). The percentage of CpG sites with informative flanking sequence ranged from 13.0% to 17.5%. Furthermore, sequence content was progressively more informative of methylation for the 3B1, 3A1 and 3A2 conditions, and co-induction of DNMT3L uniformly increased the role of sequence in determining methylation.

**Figure 4:**
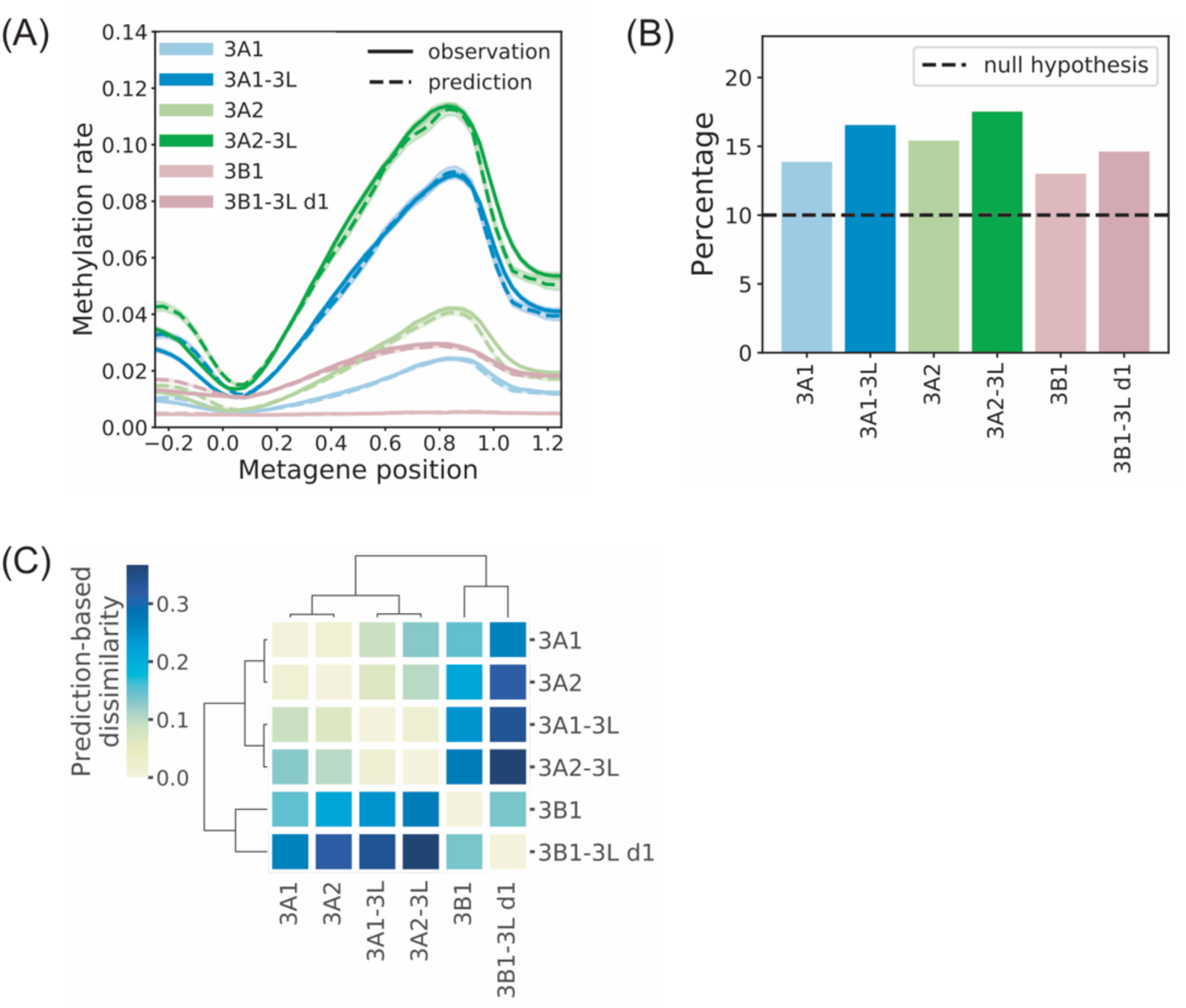
Performance of deep convolutional neural networks (CNN) for predicting DNA methylation. **(A)** Metagene plot of predicted and observed CpG methylation averaged across *K. phaffii* genes. Shading indicates 95% Bayesian credible interval for observed mCpG rates and 95% confidence interval for prediction. **(B)** Fraction of informative sequences in the test set. An informative sequence is defined as a sequence whose loss is in the top 10% of the distribution of losses for inputs obtained by sequence permutation. Dotted line shows expected percentage when sequence is not useful for prediction. **(C)** Clustering of conditions based on prediction-based dissimilarity (PBD) (Methods).

Given that sequence information contributed to methylation prediction, we next asked whether the patterns recognized for predicting high and low methylation were universal or specific to DNMT3 paralogs. For this purpose, we defined a prediction-based dissimilarity (PBD) measure between knock-in condition pairs, such that the value increased with the ability of CNNs to recapitulate observed differences in standardized mCpG rates between the conditions (Supplementary Methods: “Calculation of prediction-based dissimilarity”). On the one hand, high PBD implied that CNNs identified condition-specific sequence and metagene preferences that could be used to reliably predict differences in standardized observed methylation rates between the two conditions. On the other hand, low PBD implied that patterns of sequence and metagene preferences were similar between a pair of conditions so that CNNs could not reliably predict differences in standardized observed methylation rates between the two conditions. Hierarchical clustering of the six conditions based on PBD clearly separated DNMT3A isoforms from DNMT3B1 (Figure 4C), indicating that different sequence preferences direct *de novo* methylation by these two paralogs.

### Simulated annealing reveals distinct local nucleotide preferences of DNMT3A and DNMT3B

Given the evidence that our CNNs used paralog-specific sequence information to predict CpG methylation, we applied a network interpretation method to reveal the explicit sequence features preferred and avoided by each DNMT. Our interpretation method was based on the simulated annealing (SA) algorithm and built on previous uses of Markov Chain Monte Carlo sampling methods for extracting the learned features of CNNs (Methods; Figures S18,S19) (12). The method sampled new input sequences near maxima or minima of CNN-predicted methylation rates, retaining learned DNMT sequence preferences and aversions, respectively; and, it permitted representation of the predictive sequence features with motif logos (Figure 5A,B).

**Figure 5:**
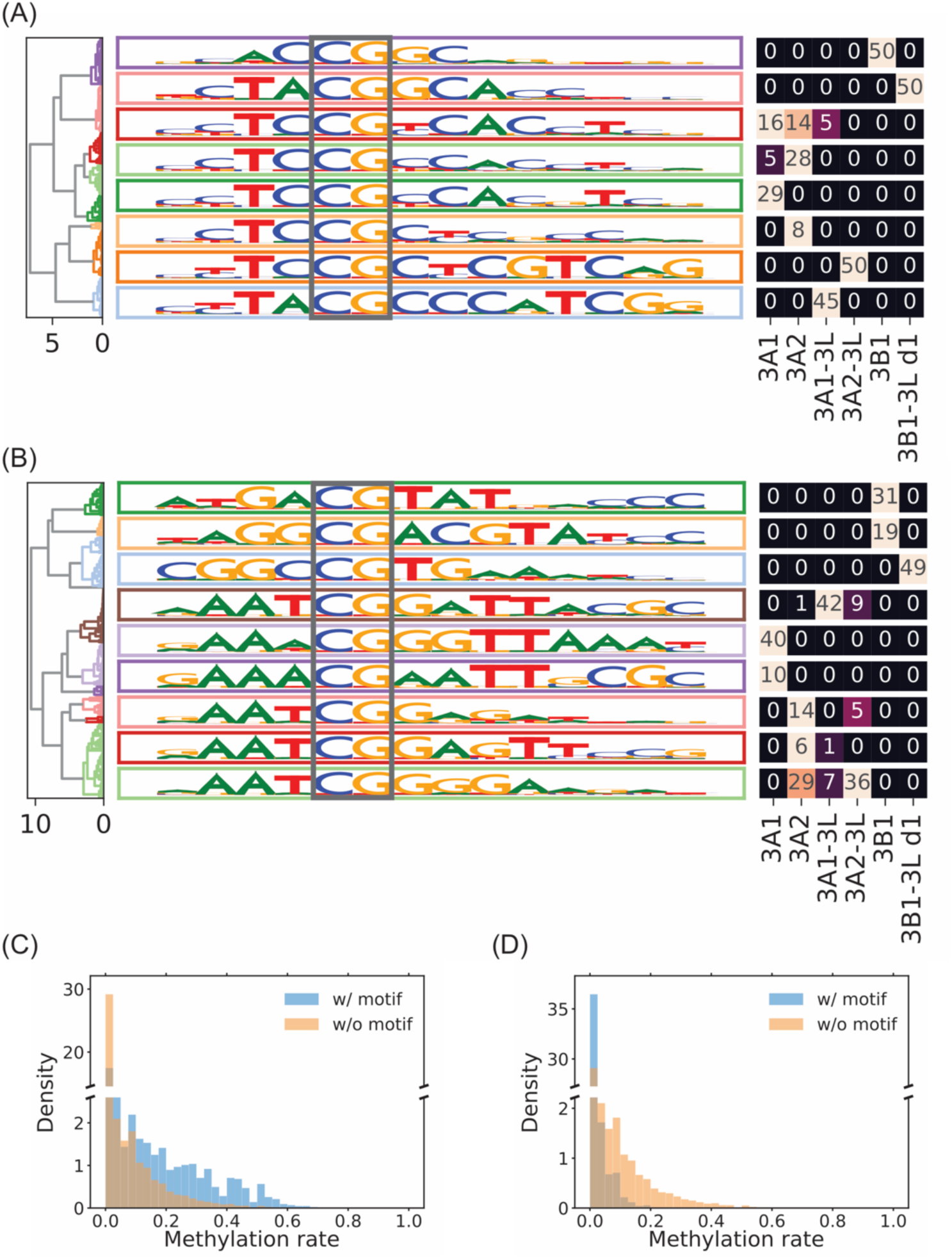
Local nucleotide patterns preferred or avoided by DNMTs. **(A)** Motif logos representing distinct patterns of nucleotide preferences identified in 300 iterations of SA interpretation. The table on the right shows composition of each motif cluster in terms of SA knock-in conditions. **(B)** Similar to (A), but for motif logos representing disfavored nucleotides. **(C)** Distributions of mCpG rates at CpG cytosines contained vs. not contained in motifs preferred by 3A1-3L. **(D)** Distributions of mCpG rates at CpG cytosines contained vs. not contained in motifs avoided by 3A1-3L.

Getting the true maximum of a neural network’s prediction function is a difficult task, and the true maximum might be the result of overfitting (44). Moreover, we could not ignore local maxima, since a DNMT might preferentially methylate multiple motifs. We therefore iterated SA using multiple initializations; for each of the six trained CNNs, SA interpretation was initialized at the top 50 test sequences with highest CNN-predicted methylation and produced 50 distinct sets of sequences sampled near maxima of the prediction function (Supplementary Methods: “Clustering of SA results”). Clustering the results of this interpretation procedure identified the motifs preferentially methylated by 3B1 and 3B1-3L, as well as those preferentially methylated by the 3A isoforms, and uncovered key nucleotides either representing shared preferences or distinguishing motif clusters (Figure 5A). The primary difference in nucleotide preference occurred at the +1 position where the 3A knock-in conditions preferred C or T, while the 3B1 knock-in conditions preferred G. Another notable difference occurred at the -1 position, where 3A1-3L and 3B1-3L preferred A, while all other conditions preferred C. Nucleotide preferences at the +/-2 positions were similar across conditions; all conditions preferred T at -2, except for the 3B1 which preferred A or T, and all conditions preferred C at +2, except for 3A2 which preferred C or T.

For motifs resisting *de novo* methylation, we initialized SA at the 50 test sequences with lowest CNN predicted methylation, sampling sequences near minima of the prediction functions, and then clustering (Figure 5B). The clustering separated the 3B1 and 3B1-3L conditions from the 3A conditions. The primary difference between motif clusters separating these conditions occurred at the -2 position, where the 3B1 conditions avoided the G nucleotide, while the remaining conditions containing 3A isoforms avoided A. Within the 3A cluster, the 3A1 isoform was also separated from the 3A2 isoform, with the 3A1 conditions largely avoiding T at the +3 position, while the 3A2 conditions largely avoided G at +3. In terms of shared features, we observed that all conditions showed aversion to A or G at the +2 position, with the exception of 3B1 which showed aversion to A or C.

To validate the identified sequence preferences and aversions, we constructed position-specific scoring matrices from the SA samples and compared the methylation rates at the CpGs in sequences matching our motifs to those at all other CpGs genome wide (Methods). This comparison validated our predictions in each case (Figure S20, Tables S6,S7). For example, for the 3A1-3L knock-in condition, methylation rates at CpGs matching a motif predicted to maximize methylation were significantly higher than other mCpG rates (Figure 5C; p = 4.57×10^-173^, Mann-Whitney U test), with mean mCpG rates 0.124 and 0.047, respectively. Similarly, methylation rates at CpGs matching a motif predicted to minimize methylation were significantly lower than other mCpG rates (p = 2.16×10^-99^, Mann-Whitney U test), with mean mCpG rates 0.019 and 0.047, respectively. Thus, the sequence features learned by each CNN and extracted by SA agreed with observed patterns of methylation *in vivo* and provided insights into distinct motifs preferentially methylated or avoided by the DNMT3A isoforms and DNMT3B1.

### Global features revealed by SA suggest a distinct local chromatin structure for regions highly methylated by the DNMT3A family

We extended the bases used to generate SA maximization motif logos from 14 bases to the full 201 bp sequence (Figure 6A for DNMT3A1-3L and Figure S21 for other conditions). As before, SA was initialized 50 times for each condition at the test set elements with greatest predicted methylation, and clustering largely separated the sequences by DNMT knock-in type (Figure S22). Repetition of poly(A) sequence appeared in all conditions, except for DNMT3B1, and was most apparent in the 3A1-3L condition (Figure 6A, Figure S21). By contrast, this periodic poly(A) pattern was absent in sequences with lowest predicted methylation (Figure S23,S24). This section therefore focuses on sequences highly methylated by 3A1-3L, and results for all six conditions are shown in Figures S25-S27.

**Figure 6:**
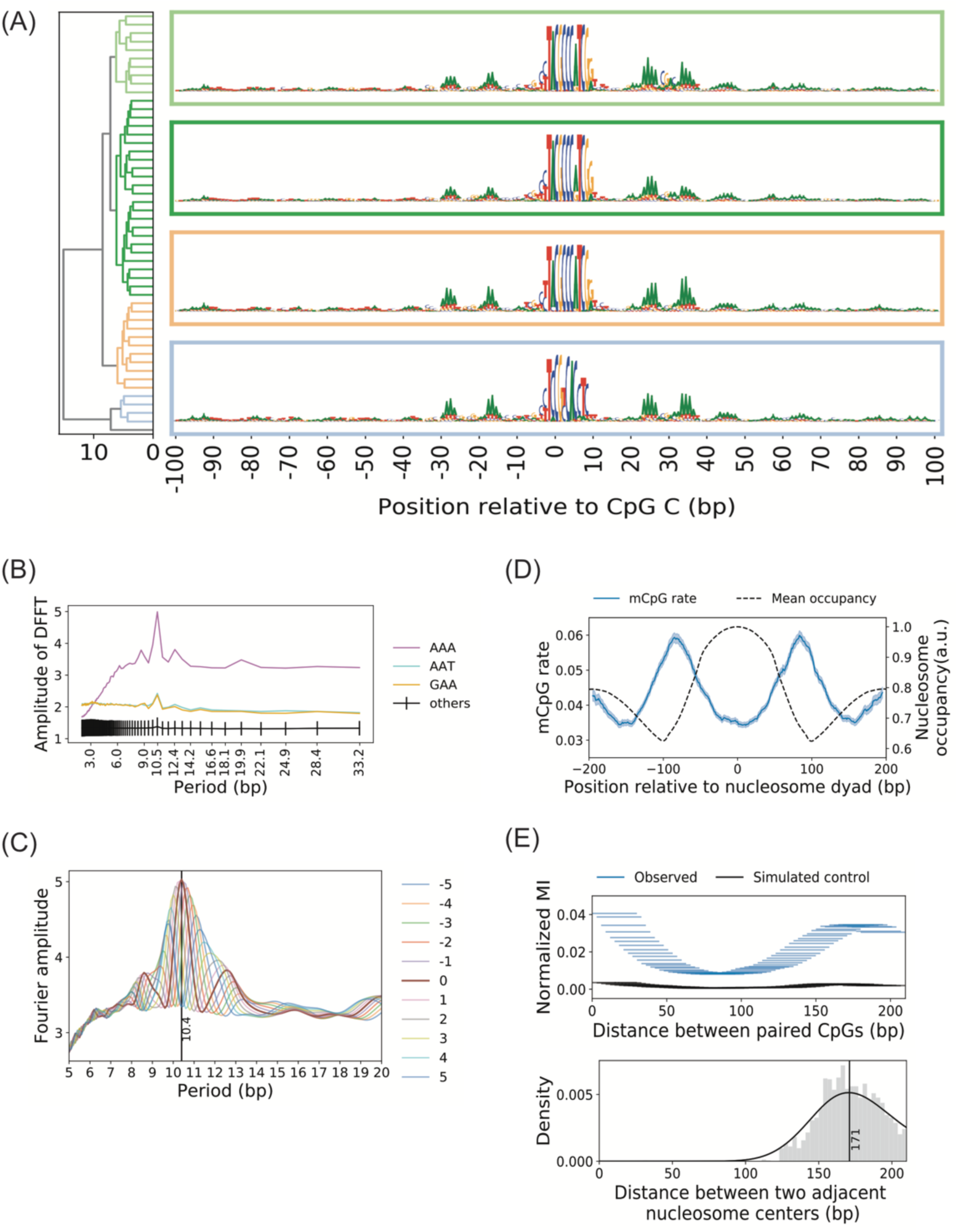
Global sequence features preferred by DNMT3A suggest a rotational positioning of CpG with respect to local chromatin structure. **(A)** Global sequence preferences of 3A1-3L learned by the CNN. **(B)** Average amplitude of discrete fast Fourier transform (DFFT) applied to 3-mer counts in individual sequences sampled by SA for the 3A1-3L condition. The top five 3-mers with highest amplitude at 10.5 bp are shown individually and amplitudes of the remaining 3-mers are summarized by their mean and standard deviation (Methods). **(C)** Periodic poly(A) pattern is in phase between the 5’ and 3’ region of CpG. Curves show the Fourier amplitude for count data obtained by shifting the AAA counts 3’ of central CpG by the indicated amount of bases. **(D)** Dyad-aligned nucleosome occupancy and corresponding mCpG rate of 3A1-3L. Shading indicates 95% confidence interval. **(E)** (Top) Mutual information (MI) of methylation status at two distinct CpG sites as a function of their separation distance, normalized by entropy. MI was estimated from the empirical joint distribution of methylation status at CpG pairs separated by a genomic distance within each indicated horizontal 30 bp window. Simulated negative control is based on independent sampling of binary methylation status from the site-specific mCpG rates. (Bottom) Distribution of distance between adjacent nucleosome dyads flanking a highly methylated CpG.

Periodic enrichment of the ploy(A) sequences could arise either from enrichment of poly(A) tracts in individual high methylation sequences or from averaging independent enrichment for mono(A) at neighboring base positions. To distinguish between these two cases and determine the period of A enrichment, we performed discrete fast Fourier transform (DFFT) on indicator variables for trinucleotide content in individual SA sequence samples as a function of genomic position relative to the CpG and then averaged the resulting amplitudes across samples for the same DNMT condition. Before calculating each transform, we replaced the central 19 bps with zeros to prevent the dominating motif from skewing the spectral analysis (Supplementary Methods: “Fourier analysis of periodic trinucleotides”). The strongest peaks in spectral amplitudes consistently occurred at period 10.5 bp and was largest for AAA in all DNMT3A conditions (Figure 6B, Figure S25A). Interestingly, this pattern was less pronounced for DNMT3B; for 3B1-3L d1 we observed only a small peak at 10.5 bp, while 3B1 did not show any notable 10.5 bp periodicity of trinucleotides (Figure S25A). Thus, our SA interpretation method revealed that the CNNs trained on DNMT3A conditions utilized the presence of AAA at 10.5 bp periodicity to predict high CpG methylation rate.

We next sought to determine the relative phase of poly(A) between the 5’ and 3’ sides of the central CpGs in our SA samples by progressively shifting the 3’ region by a single base. For each shift, we calculated the spectral amplitude of indicator variables for AAA in each sequence by taking the amplitude of the dot product with discrete complex exponential functions with period between 5 and 20 bp. Averaging the result across sampled sequences for 3A1-3L demonstrated that AAA counts attained maximum period at 10.4 bp with zero shift, implying that poly(A) patterns were in phase by full helical turns between the 5’ and 3’ sides of highly methylated CpGs (Figure 6C). Similar results held for other DNMT3A family knock-in conditions, but not for the DNMT3B family (Figure S25B). By contrast, lowly methylated sequences did not show any reproducible periodic pattern of trinucleotides (Figure S26).

It is well known that nucleosomal DNA is sometimes enriched for AA/TT dinucleotides with 10.5 bp periodicity and these dinucleotides tend to be positioned in the minor groove facing toward the histone core (45–50), thereby contributing to the rotational positioning of nucleosomes *in vivo* (46, 47). Given the known role of periodic AA/TT in nucleosome positioning, the presence of periodic AAA in phase on both sides of the CpG could imply that the CNN learned to associate high methylation on nucleosomal DNA with specific rotational positioning of CpGs with respect to the histone core. An alternative possibility, however, would be that highly methylated CpGs actually resided in the linker DNA between two phased nucleosomes, as supported by the observed preferential methylation of linker DNA by each DNMT (Figure 6D, Figure S27A). In this case, the presence of periodic AAA on both sides of a CpG and in phase by full helical turns might function to enforce a specific relative orientation of two adjacent nucleosomes flanking the CpG. We estimated the linker length between adjacent nucleosomes flanking a highly methylated CpG to be ∼25 bp (Figure 6E), larger than the ∼20 bp estimated from the genome-wide nucleosome repeat length of 166 bp observed in *K. phaffii* (19). Interestingly, a recent study shows that 10n linkers may facilitate compact two-start zigzag stacking of nucleosomes in heterochromatin, while 10n+5 linkers may support more flexible, reversibly repressed chromatin structure (51).

Since wrapping DNA around histones can bring distal CpG sites to proximity in three-dimensional space and facilitate their co-methylation as a unit, we next investigated whether the coupling of methylation across the genome reflects local chromatin structure. For this purpose, we computed the normalized mutual information (MI) of methylation status for CpG pairs found within individual bisulfite sequencing fragments, binning all such pairs according to their distance (using a bin width of 30 bp, Supplementary Methods: “MI calculation”). As control, we performed a simulation that replaced each observation of methylation status with a simulated observation independently sampled according to the mCpG rate at the corresponding site and recomputed the MI of methylation between paired observations. This analysis confirmed that the effect of periodicity of marginal methylation rate (Figure 6D) on MI was minimal (Figure 6E). The result clearly demonstrated that mutual information of CpG methylation status was highest for CpGs in close proximity (within 30 bp) and decayed as a function of separation distance, but then peaked again for CpG pairs roughly one nucleosome repeat length apart (Figure 6E). This second peak supported our hypothesis that CpGs on opposite linkers of the same nucleosome might be brought physically close to each other by the three-dimensional nucleosome structure and get frequently methylated as a unit. We found similar results for all other active DNMT conditions, except for the 3B1 single knock-in (Figure S27B).

## DISCUSSION

By introducing *de novo* DNA methyltransferases in a simple eukaryote that naturally lacks DNA methylation, we have characterized molecular adaptation of cells to ectopic DNA methylation stress, genetic information that guides the deposition of this epigenetic mark, and functional consequences at the level of individual gene transcription. Overall, it is surprising that each of the three *de novo* DNMT isoforms tested was by itself sufficient to reproduce many of the characteristics of DNA methylation in mammals, including synergistic interaction with DNMT3L, depletion of methylation at gene TSS, increasing methylation along gene bodies, and an inverse relationship between *de novo* methylation and transcription activity. Although we observed that global methylation rates were much lower than those found naturally in mammalian cells, it is intriguing that knockout of *de novo* DNMTs and DNMT1 from mouse ES cells followed by reintroduction of either DNMT3A2 or 3B1 also resulted in low CpG methylation rates of 0.07 and 0.028, respectively, similar to our findings in *K. phaffii* (16). Beyond these general similarities, our results provide several important insights.

First, *de novo* methylation in *K. phaffii* poses cellular stress that elicits a broad transcriptional response, demonstrating that DNMT expression disrupts normal cellular processes either through consumption of metabolites or direct effects of DNA methylation on transcriptional output. Our result contrasts with a previous study that found minimal differential expression after knocking in murine *DNMT3B1* in *S. cerevisiae* (17). This difference, potentially due to differences in species, is important, since our result shows that DNMT introduction and DNA methylation can significantly affect transcription in the absence of co-evolved cellular machinery interacting with DNA methylation.

Second, we report a coordinated transcriptional response that counters DNA methylation stress by limiting the abundance of the DNMT methyl donor SAM. The effect of SAM levels on DNA methylation and the role of the methionine cycle in modulating SAM availability have been observed previously in several cell types including Schwann cells, neurons of the dentate gyrus, and peripheral lymphocytes (14,52,53). Moreover, dysregulation of SAM has been implicated in diseases. For example, decreased SAM/SAH ratios are associated with DNA hypomethylation and poorer prognosis in hepatocellular carcinoma (54) and with hypomethylation and gene dysregulation in a mouse model of diabetic neuropathy (14). In line with these roles of SAM in regulating the mammalian methylome, our study shows that cells lacking intrinsic DNA methylation are still able to detect DNA methylation stress and mitigate this stress by modulating SAM cycle genes. This surprising result indicates that cellular mechanisms regulating DNA methylation via SAM are conserved across a diverse group of eukaryotes, including those lacking endogenous DNMTs.

Furthermore, our *K. phaffii* model recapitulates several details of the inverse relationship between gene methylation and transcription previously observed in species with endogenous DNMTs. For example, performance of our “DE down vs. rest” classifier was uniformly superior to the performance of our “DE up vs. rest” classifier. This finding is consistent with hypermethylation being a marker of transcription repression (55) and with the fact that absence of repression is not sufficient for hypertranscription of a gene. Specifically, the regression coefficients of our classifiers reveal that methylation of CpG sites located just downstream of the TSS plays the most important role in predicting differential gene expression status, while the second most important location corresponds to the TSS and promoter regions. This order of importance agrees with the recent finding that differential methylation in the region immediately 3’ of the TSS is most informative for predicting differential expression among tissues profiled in the Roadmap Epigenomics Project (56). Thus, our minimal *K. phaffii* system confirms the functional role of DNA methylation in higher-order eukaryotes.

Using the controlled epigenetic perturbation data, our CNN-based prediction of CpG methylation has quantified the effect of flanking sequence on methylation and discovered motifs preferred and avoided by each *de novo* DNMT. Specifically, we have shown that sequence is most important in determining methylation by the DNTM3A2 isoform, followed by DNTM3A1 then DNMT3B1, and that the combination of each of these isoforms with DNMT3L increases the effect of sequence on methylation. The reduced role of sequence in determining DNMT3B1 methylation is consistent with a previous report that H3K36me3 is important in recruiting DNMT3B1, but not DNMT3A2 (16), suggesting that DNMT3B1’s affinity for H3K36me3 may limit the effect of its intrinsic sequence preference.

In terms of specific motifs preferred and avoided by each isoform, our SA-based network interpretation resolves several discrepancies among previous investigations. Overall, our results are in best agreement with those of Wienholz *et al.* who studied *de novo* methylation of episomal DNA transfected in HEK293c18 cells and found sequence dependence of CpG methylation (13). Their findings that DNMT3A1 prefers T at -2 and C at +2, while avoiding A at -2, agree well with our results. They also report that DNMT3B1 prefers T at -1 and G at +1 and avoids C at +1. While our analysis confirms that DNMT3B1 prefers G at +1, our other findings related to DNTM3B1 differs from those of Wienholz *et al.*, perhaps due to the relatively small role of sequence in determining DNTM3B1 methylation in our data. In an earlier study of sequence preference, Handa *et al.* identified the consensus sequences CTTGCGCAAG and TGTTCGGTGG associated with high and low CpG methylation, respectively, in human cells (15). Because methylation at these consensus sequences likely resulted from combined effects of DNMT3A and DNMT3B, we may compare the consensus sequences with common patterns of preference and aversion shared across our results (Figure 5A,B). The high methylation consensus sequence agrees well with our results, except at positions -1 and +2. At +2, our results indicate a preference for C/T and suggest that the A predicted by Handa *et al.* may actually be disfavored; both of these findings are supported by data from Wienholz *et al.* Similarly, our results disagree with the low methylation consensus sequence of Handa *et al.* at position +2 and positions 5’ of -1, while agreeing with the findings of Wienholz *et al.* at the +2 and -2 positions. Thus, our results clarify paralog-specific sequence preferences that contribute to methylation landscapes.

Finally, our study highlights a connection between DNA methylation and nucleosome structure mediated by periodic poly(A) sequence features that extend beyond the core sequence motifs preferred by DNMTs. Further experimental investigations are needed to determine whether the poly(A) sequences play a role in rotational positioning of adjacent nucleosomes for preferential methylation of linker DNA or they restrict the rotational position of nucleosomal CpGs on the histone core. It also remains to be seen whether ordered arrays of centromeric nucleosomes, as those reported in *S. pombe* (57), can directly recruit DNMT3B1 (Figure S17B,C), indicating an evolutionarily conserved interaction between CENP-A/C histones with DNMT3B1. Nevertheless, the observed separation of peaks in mCpG mutual information by the nucleosome repeat length signals that the three-dimensional organization of DNA in chromatin is important for determining the pattern of *de novo* DNA methylation *in vivo*.

## FUNDING

This work was supported in part by funds from National Institutes of Health R01CA163336 and the Grainger Engineering Breakthroughs Initiative to JSS; the L.S Edelheit Family Biological Physics Fellowship to AF; an American Heart Association Scientist Development Grant (17SDG33650087) and National Institutes of Health R01GM127497 to PP; and by the National Science Foundation Graduate Research Fellowship Program under Grant No. DGE – 1746047 to M.G. Funding for open access charge: R01CA163336.

