## Supplementary Methods, Tables and Figures for "Epigenetic engineering of yeast reveals dynamic molecular adaptation to methylation stress and genetic modulators of specific DNMT3 family members"

**This PDF file includes:**

Supplementary Methods
Supplementary Tables S2, S3, S6 and S7
Supplementary Figures S1 to S27
References for Supplementary Methods and Figures

### Supplementary Methods:

#### RNA-seq analysis

Paired-end RNA-seq reads were mapped to the *K. phaffii* genome using TopHat2 (1) (version 2.1.1) with the options -g 1 --prefilter-multihits --library-type fr-unstranded. Mapping reads to an augmented reference genome containing all human DNMT sequences caused frequent multiple alignment to different DNMTs, because of the sequence similarity of isoforms (data not shown). To prevent these ambiguous alignments, we created separate versions of the genome that contained only one of the knocked-in DNMT1A, DNMT3A1, DNMT3A2, and DNMT3B1 sequences, as well as all of the DNMT3L, KanR, and ZeoR sequences and the *K. phaffii* reference genome. Reads from each condition expressing DNMT1A, DNMT3A1, DNMT3A2, or DNMT3B1 were aligned to the genome version containing the corresponding sequence. Reads from 3L and control conditions were aligned to the version of the genome containing all DNMT, KanR, and ZeoR sequences and the *K. phaffii* reference. Mapped reads were filtered with samtools view (2) (version 1.7) using the options -h -f 3 -F 3596 -q 13, and read pairs mapping to different chromosomes were removed. Gene expression values, in units of FPKM (fragments per kilobase of transcript per million mapped reads), were calculated from filtered bam/sam files using cuffnorm (3) (version 2.2.1) with the options --library-norm geometric --library-type fr-unstranded.

#### DNMT Western blot

Cell pellets were resuspended in 1X NuPAGE LDS Sample Buffer (Life Technologies) with 2.5 % β-mercaptoethanol, incubated for 5 minutes at 95°C, sonicated, and incubated for 5 minutes at 95°C again. Lysates were loaded onto a 4-12% NuPAGE Bis-Tris Protein Gel alongside WesternSure prestained ladder (LiCor), electrophoresed, and transferred onto a 0.45 µm nitrocellulose membrane (BioRad). Total protein was stained using Revert Total Protein Stain (LiCor) and imaged at 700 nm on an Odyssey Fc Imager (LiCor). Membranes were then blocked for 1 hour with 5% nonfat dried milk in TBS (50 mM Tris pH 7.4 and 150 mM NaCl) containing 0.1% Tween-20 (TBST). Membranes were rinsed three times in TBST and incubated with anti-human DNMT rabbit antibodies (1:1000, Cell Signaling Technology, DNMT1 #5032, DNMT3a #3598, DNMT3a2 #32578, DNMT3B1 #57868, or Sigma, DNMT3L #SAB1401663) according to manufacturer's instructions. Following primary antibody incubation, membranes were rinsed three times in TBST and incubated for 1 hour with HRP-conjugated anti-rabbit IgG (1:2000, Cell Signaling Technology, #7074). Membranes were then rinsed with TBST three times and incubated for five minutes in Clarity Western ECL Substrate (BioRad). Membranes were imaged with a 30 second exposure time on an Odyssey Fc imager.

**WGBS analysis**

We used Bismark (4) (version 0.18.0) to process the whole-genome bisulfite sequencing (WGBS) data for two biological replicates from each of our 13 conditions. Paired-end reads from each experiment were trimmed using Trim Galore (https://github.com/FelixKrueger/TrimGalore, version 0.4.4) with the command trim_galore --paired --trim 1 --clip_r1 8 --clip_r2 8 --three_prime_clip_r1 8 --three_prime_clip_r2 8, followed by removal of read duplicates.

Reads were aligned to the bisulfite converted *K. phaffii* genome with Bismark as follows: we first sought to align reads as pairs using the command bismark --bowtie2 -N 1 -D 25 -R 5 --score_min L,0,-0.4. --X 2000 --unmapped. Then, to raise alignment rates, we attempted a second alignment of the first-strand reads from unaligned read pairs using the command bismark --bowtie2 -N 1 -D 25 -R 5 --score_min L,0,-0.4. We did not attempt to align the second read from initially unmapped read pairs, because alignment rates for this set of reads were much lower than those of the first reads (data not shown). Table S3 provides paired-end and first-strand single-end alignment rates along with coverage and bisulfite non-conversion rates estimated from spike-in unmethylated lambda phage DNA.

After alignment, we obtained summary files which described, for each read, the methylation status at CpG-context cytosines by running the command "bismark_methylation_extractor --no_overlap --comprehensive" on the bam files produced by paired-end alignment and the command "bismark_methylation_extractor --comprehensive" on the bam files produced by single-end alignment. The resulting files were concatenated into a single CpG-context file that served as input for mutual information calculations in Supplementary Methods: "MI calculation."

Finally, we generated CpG report files containing the counts of methylated and unmethylated observations aggregated across aligned reads for each C in the CpG context. These reports where generated by running the command bismark2begGraph on the concatenated CpG-context file followed by running the command coverage2cytosine. The CpG report files associated with each WGBS experiment were used to estimate methylation rates at each C in a CpG context.

**Clustering experiments by window average methylation**

To check the reproducibility of WGBS experiments, we clustered the WGBS samples represented by vectors of window averaged maximum likelihood estimated methylation probabilities for CpG context cytosines. Window averaging for each sample used window width of 4 kb and stride of 4 kb. Hierarchical clustering used average linkage and was based on Pearson dissimilarity of window average vectors (Figure S1A).

**Empirical Bayes approach for estimating mCpG rate**

We used an empirical Bayes approach to estimate the probability of methylation at C's in the CpG context, which we termed mCpG rates, from the observed counts of methylated and unmethylated status (Supplementary Methods: "WGBS analysis"). This approach combines the number $m_i$ of methylated observations at site $i$ and the number $n_i$ of unmethylated observations at site $i$ with a beta prior derived from genome-wide data. The genome-wide information moderates the maximum likelihood estimates (MLEs) $m_i/(m_i + n_i)$ in the case of low coverage.

Specifically, for C with index $i$, the probability of methylation $\Theta_i$ is a random variable described by a posterior distribution equal to the product of a beta prior on $\Theta_i$ and a binomial likelihood for observations, divided by a normalizing factor:

$$\mathbb{P}\,(\,\Theta_\mathrm{i} = \theta\;|m_i, m_i + n_i, \alpha, \beta) = \frac{f(m_i\;|m_i + n_i, \theta)B(\theta|\alpha,\beta)}{\int_0^1 f(m_i\;|m_i + n_i, \theta)B(\theta|\alpha,\beta)\,d\theta}.$$

Here $f$ denotes the binomial probability mass function, and $B$ denotes the beta probability density function with shape parameters $\alpha, \beta$. The Empirical Bayes approach chooses $\alpha, \beta$ to maximize the Bayesian evidence at all CpG-context C's, when these observations are assumed independent:

$$\begin{aligned}\hat{\alpha}, \hat{\beta} &= \underset{\alpha,\beta}{\operatorname{argmax}} \prod_i \int_0^1 f(m_i|m_i + n_i, \theta)B(\theta|\alpha,\beta)\,d\theta \\ &= \underset{\alpha,\beta}{\operatorname{argmax}} \sum_i \log\left(\int_0^1 f(m_i|m_i + n_i, \theta)B(\theta|\alpha,\beta)\,d\theta\right).\end{aligned}$$

The R package ebbr (https://github.com/dgrtwo/ebbr, version 0.1) was used to obtain an MLE pair $\hat{\alpha}, \hat{\beta}$ for each condition after first restricting to C's with at least one observation. After fitting these prior parameters for each condition, we estimated the mCpG rate $\hat{p}_i$ at the $i^{th}$ CpG C as the Bayes posterior mean of $\Theta_i$,

$$\begin{aligned}\hat{p}_i &= \mathbb{E}_{\mathbb{P}\,(\,\Theta_\mathrm{i}\;|m_i, m_i + n_i, \hat{\alpha}, \hat{\beta})}[\,\Theta_\mathrm{i}\,] \\ &= \frac{m_i + \hat{\alpha}}{m_i + \hat{\alpha} + n_i + \hat{\beta}}.\end{aligned}$$

**Calculation of feature-aligned mCpG rates**

We summarized trends in mCpG rate relative to genomic features, such as transcription start sites (TSSs) or nucleosome dyads, by applying a sliding window average to relative genomic coordinates. To define our window average statistic, let $\{h_j(i)\}_{j \in F}$ denote the set of functions mapping the genomic coordinate and strand information of a cytosine at location $i$ to a signed genomic distance relative to feature $j$ in the set $F$ of all genomic features of interest. For example, if $F$ denotes the set of all gene TSSs, then $h_j(i)$ is the signed number of base pairs from the TSS of gene $j$ to the C labeled by $i$, where the 3' direction of the gene's coding strand is taken as positive. Denoting the set of all CpG-context C's within a fixed sliding window starting at position $w$ and with width $\Delta w$ as

$$I(w) = \{i : w \leq h_j(i) < w + \Delta w \text{ for some } j \in F\},$$

the mean methylation probability in the window is the random variable

$$\overline{\Theta}(w) = \frac{1}{|I(w)|} \sum_{i \in I(w)} \Theta_i.$$

For each window, we summarized this random variable by its expected value

$$\mathbb{E}[\overline{\Theta}(w)] = \frac{1}{|I(w)|} \sum_{i \in I(w_1)} \hat{p}_i ,$$

where $\hat{p}_i$ is the posterior mean of $\Theta_i$, as previously described. In the main text, we refer to both $\mathbb{E}[\overline{\Theta}(w)]$ and $\hat{p}_i$ as mCpG rates, since it is clear from the context whether window averaging has occurred.

We used 95% credible intervals to quantify the distribution of $\overline{\Theta}(w)$ about its expected value. These intervals were estimated using a bootstrap approach. For each window $w$, we repeatedly drew independent samples of size $|I(w)|$ from $I(w)$ with replacement, called this collection of samples $B(w)$, and sampled from the distribution

$$\bar{\theta} \sim \frac{1}{|B(w)|} \sum_{i \in B(w)} \Theta_i.$$

The lower and upper bounds of the 95% credible interval for $\overline{\Theta}(w)$ were the 2.5 and 97.5 percentiles of the distribution of sampled $\bar{\theta}$.

We applied this method to three cases which used the following choices for $h_j$, window widths, and window strides. First, to summarize methylation trends relative to gene TSSs, we defined $h_j(i)$ to be the number of base pairs (bp) from the TSS of gene $j$ to the C at site $i$, taking the 3' direction of the gene's coding strand as positive. The window width and stride were 50 bp and 10 bp, respectively. Second, to summarize methylation trends in metagene coordinates, defined as the distance downstream of TSS scaled by gene length, we modified the preceding definition of $h_j(i)$ by dividing the signed distance between gene TSS and C by gene length. The window width and stride were 0.15 and 0.03, respectively. Finally, to summarize methylation trends relative to nucleosome dyads, we defined $h_j(i)$ to be the signed number of bp from the nucleosome dyad to the C at site $i$, taking the 3' direction of the strand containing the C as positive. The window width and stride were 3 bp and 1 bp, respectively. Credible interval estimates for all windows used 100 bootstrap samples.

**Gene expression PCA**

Log-scale gene expression values were obtained from expression counts by applying variance stabilizing transformation to library-size-normalized count values using the vst function of DEseq2 (version 1.14.1) with option blind=FALSE (5). Library-size normalization used the DESeq median of count ratios method (6). Next, we used the removeBatchEffect function of the limma package (version 3.30.13) to remove variation in transformed expression that was attributable to the three batches in which RNA-seq experiments were performed (7). PCA was performed on the resulting batch-corrected log-scale expression values using scikit-learn (8).

**Gene ontology analysis and GO term clustering**

We used DAVID to perform gene ontology (GO) analysis, testing each knock-in condition for functional enrichment in genes sets with expression significantly increased relative to control ("DE up" genes) and gene sets with expression significantly decreased relative to control ("DE down" genes) (9). As a preliminary step, *K. phaffii* gene names annotated by Love *et al.* based on homology to *S. cerevisiae* genes (10) were converted to Saccharomyces Genome Database IDs (SGD IDs) (11). This conversion allowed DAVID to use a custom gene background composed of only those genes identified in *K. phaffii*. Restricting to genes with valid SGD IDs, we then tested the "DE up" and "DE down" gene sets of each knock-in condition for functional enrichment compared to the background of all *K. phaffii* genes. Tests were performed via programmatic access to DAVID using the RDAVIDWebService package (12), and GO terms with Benjamini-Hochberg adjusted p-values less than 0.05 were accepted as being enriched. Supplementary Table S5 provides enriched terms for both "DE up" and "DE down" genes in each condition.

To summarize the GO results for our time-course data, we performed hierarchical clustering on those GO terms enriched in any of the "DE up" and "DE down" gene sets on days

1-4 (Figure S10; Supplementary Table S5). Distance between a pair of GO terms $T_1$ , $T_2$ was calculated as

$$1 - J(T_1, T_2)$$

where $J(T_1, T_2)$ is the Jaccard similarity between the set of all DE genes ("DE up" or "DE down" on any day) associated with $T_1$ and the set of all DE genes associated with $T_2$. Hierarchical clustering based on these distances used average linkage. Clusters were determined using the fcluster of scipy (https://www.scipy.org/) function with the "inconsistent" criterion and parameters t=1.4, depth=4.

**SAM measurements**

Approximately 150 million cells were collected from each strain studied over a 5 day time course, pelleted by centrifugation and frozen at -80°C until analysis. For SAM quantification, cell pellets were lysed in 125 µL of Y-PER Yeast Protein Extraction Reagent (Thermo Scientific) at room temperature with agitation for 20 minutes. Cell debris was pelleted by centrifugation at 14,000 x g for 10 minutes. The supernatant was collected and diluted with PBS to the equivalent of 4E8 cells per mL.

SAM concentration of 50 µL (2E7 cells) diluted samples was analyzed in duplicate using a competitive S-Adenosylmethionine (SAM) ELISA Kit (Cell BioLabs, Inc.) according to manufacturer's instructions. Briefly, a 96-well protein binding plate was coated overnight with SAM conjugate. A SAM-BSA standard curve and Pichia samples were added to the plate and incubated for 10 minutes on a rocking platform followed by addition of 50 µL anti-SAM antibody and incubation on a rocking platform for 1 hour at room temperature. The plate was then washed 3 times with 250 µL wash buffer and 100 µL secondary antibody conjugated to horseradish peroxidase (HRP) was added and incubated at room temperature for 1 hour. After washing 3 more times, 100 µL HRP substrate solution was added and incubated at room temperature on a rocking platform for 15 minutes. The reaction was stopped by addition of 100 µL of stop solution and absorbance at 450 nm and 620 nm was measured using a Synergy HTX Multi-Mode Reader (BioTek).

To analyze the results, the 620 nm absorbance values (reference) were subtracted from the 450 nm absorbance values and the resulting values were subtracted from the average of the standard curve blank wells. Standard curve absorbances were plotted against their known concentrations and a logarithmic curve was fit to the data ($R^2 = 0.988$). This curve was used to determine SAM concentrations in each sample.

**Clustering gene expression changes and methylation patterns**

To identify methylation patterns associated with genes following common time-course differential expression trajectories, we clustered differentially expressed genes by their log-fold change relative to control. Clustering was restricted to genes that were differentially expressed for at least one day. Each gene was represented by a 4-dimensional vector of its $\log_2$ fold-change relative to control on day 1-4. We computed the pair-wise Euclidean distance between these vectors and applied hierarchical clustering with average linkage (left panel in Figure 3B). This hierarchical clustering was also used to organize the rows of mCpG heatmap in the right panel of Figure 3B. To construct the gene-level mCpG rate heatmap, we represented each gene by a 4-dimensional array of its mean mCpG rate in the metagene interval [-0.2,0.4] on time-course days 1-4. Array elements corresponding to the same day were standardized by median centering and scaling by the median absolute deviation. After standardization, we took the block median of groups of 100 genes on each day and applied $\tanh(x)$ transformation to suppress extreme values for visualization purpose (right panel in Figure 3B).

**Description of logistic regression tasks and input**

We used two-class logistic regression models to predict time-course differential expression status from metagene methylation patterns and time-course day. We considered two classification tasks. The first task, termed “DE down vs. rest”, aimed to distinguish “DE down” genes from genes that were not “DE down” (“DE up” or “not DE”). The second task, termed “DE up vs. rest”, aimed to distinguish “DE up” genes from genes that were not “DE up” (“DE down” or “not DE”).

To construct the dataset for each task, we partitioned the metagene interval [-0.2,1.2] into 30 equally sized bins. Next, for each time-course day, we performed bin-wise standardization of methylation inputs as follows: letting $\hat{p}_{i,g,d}$ denote the mCpG rate at the $i^{th}$ CpG-context C in the metagene interval [-0.2,1.2] of gene $g$ on day $d$, the corresponding standardized regression input was

$$x_{i,g,d} = \tanh\left(\frac{1}{50}\,\frac{\hat{p}_{i,g,d} - c_{B(i,g)}}{s_{B(i,g)}}\right).$$

In this equation,

- the function $B(i, g)$ maps the index $i$ of C and the associated gene $g$ to the metagene bin $b$ containing the C;
- $c_b$ and $s_b$, used to standardize within each bin, are respectively the median mCpG rate and the difference between the 75$^{th}$ and 50$^{th}$ percentiles of mCpG rates with metagene coordinates within 0.05 of the bin center; and

- the $\tanh$ operation severs to limit the magnitude of outlier mCpG rates.

For each gene and day, we supplemented the standardize methylation inputs $x_{i,g,d}$ with a length-four input $\boldsymbol{w}$ containing one-hot encoding of the time-course day. For day $d$, the components of $\boldsymbol{w}$ were

$$w_{d\prime} = \delta_{d\prime,d}$$

where $\delta_{d\prime,d}$ is the Kronecker delta function.

For the "DE down vs. rest" task, the response variables were

$$y_{g,d} = \begin{cases} 1, \text{g is "DE down" on day d} \\ 0, \text{otherwise} \end{cases},$$

while for the "DE up vs. rest" task, the response variables were

$$y_{g,d} = \begin{cases} 1, \text{g is "DE up" on day d} \\ 0, \text{otherwise} \end{cases}.$$

In summary, the final dataset for each task consisted of the input/output pairs

$$\left( \left[ \left(x_{i,g,d}\right)_{i \in I(g)}, \left(w_{d'}\right)_{d' \in \{1,2,3,4\}} \right], y_{g,d} \right)_{g \in G, d \in \{1,2,3,4\}}$$

where $I(g)$ denotes the set of indices of CpG-context C's associated with gene $g$ and $G$ denoes the set of all genes.

**Logistic regression model**

Our logistic regression models were parameterized by 30 weights $\boldsymbol{\beta} = \left(\beta_j\right)_{j \in \{1..30\}}$, associated with the metagene methylation bins, and by four weights $\boldsymbol{\alpha} = \left(\alpha_d\right)_{d \in \{1..4\}}$, associated with the time-course day indicators. To prevent overfitting, we imposed a prior on the distribution of $\boldsymbol{\beta}$ to penalize weight configurations fluctuating rapidly as a function of metagene position:

$$\mathbb{P}(\boldsymbol{\beta}) \propto \exp\left( -\frac{\lambda}{2} \sum_{j=1}^{28} \left( \frac{\beta_j - 2\beta_{j+1} + \beta_{j+2}}{30^2} \right)^2 \right),$$

where $(\beta_j - 2\beta_{j+1} + \beta_{j+2})/30^2$ is proportional to the discrete approximation of the second order derivative of the weights $\beta_j$ with respect meta gene position. Combining this prior with the likelihood of response variables given model parameters, the log posterior probability of $\boldsymbol{\alpha}, \boldsymbol{\beta}$ was

$$\log \mathbb{P}\left(\boldsymbol{\alpha},\boldsymbol{\beta} \mid (y_{g,d})_{g\in G,\, d\in\{1,2,3,4\}}\right)$$
$$= \left(\sum_{g\in G}\sum_{d=1}^{4} y_{g,d}\log \mathbb{P}(y_{g,d}=1\mid \boldsymbol{\alpha},\boldsymbol{\beta}) + (1-y_{g,d})\log\left(1-\mathbb{P}(y_{g,d}=1\mid \boldsymbol{\alpha},\boldsymbol{\beta})\right)\right)$$
$$-\frac{\lambda}{2}\sum_{j=1}^{28}\left(\frac{\beta_j - 2\beta_{j+1}+\beta_{j+2}}{30^2}\right)^2 + \text{const.}$$

where

$$\mathbb{P}(y_{g,d}=1\mid \boldsymbol{\alpha},\boldsymbol{\beta}) = \frac{\exp\left(\sum_{d'=1}^{4}\alpha_{d'}w_{d'} + \sum_{i\in I(g)}\beta_{B(i,g)}\,x_{i,g,d}\right)}{1+\exp\left(\sum_{d'=1}^{4}\alpha_{d'}w_{d'} + \sum_{i\in I(g)}\beta_{B(i,g)}\,x_{i,g,d}\right)}.$$

For fixed $\lambda$, we determined the values of $\boldsymbol{\alpha},\boldsymbol{\beta}$ maximizing this log posterior probability using the root function from the optimize package of scipy (https://www.scipy.org/) with options: method = "hybr", maxfev = 34000 and factor = 10.

**Choice of logistic regression smoothness penalty**

We selected the value of $\lambda$ using 10-fold cross validation (CV). For each fixed value of $\lambda$ in the set $\{10^k : k \in \{-5,-4,\dots 1\}\}$, we measured the following performance statistics: the area under the receiver operating characteristic (AUROC), the area under the precision recall curve (AUPRC), and AUPRC minus baseline (Figure S2-S4). The baseline in the "AUPRC minus baseline" statistic was calculated using

$$\text{baseline} = \frac{\sum_{(g,d)\in T} y_{g,d}}{\sum_{(g,d)\in T} 1},$$

where $T$ is the set of (gene, day) pairs in the test set. We measured the three performance statistics on each CV fold's full test set as well its restriction to an individual time-course day. Figure S2-S4 summarizes these statistics. For both "DE down vs. rest" and the "DE up vs. rest" tasks, we selected a value of $\lambda$ that yielded best median AUROC averaged across days: $\lambda = 0.01$ for both tasks.

Finally, for this best $\lambda$ value, we retrained the models on all available data (no holdout data) and estimated 95% confidence intervals for regression parameters via bootstrap resampling of the training data with 100 iterations (Figure 3E). To assess how $\lambda$ affects determination of relative importance of methylation at different metagene positions, we performed the same procedure for several other values of $\lambda$: $\lambda \in \{10^k : k \in \{-4,-3,-1\}\}$ (Figure S16C).

**Density estimation for minimum FPRs**

Each “DE down” gene on a given time-course day was assigned a minimum false positive rate (FPR) for correct classification by considering the “DE down vs. rest” classifier performance on the cross-validation fold that contained the gene in its test set. Because each pairing of a “DE down” gene and a time-course day appeared in the test set of exactly one cross-validation fold, this procedure assigned exactly one minimum FPR to each pair. The set of minimum FPRs associated with all such pairs was partitioned into PER and non-PER gene sets. Treating PER and non-PER genes separately, we built histograms of minimum FPRs for “DE down” genes using 100 bins partitioning the [0,1] interval. Both histograms were smoothed with kernel density estimation using gaussian kernels with bandwidth 0.1. Kernels were renormalized to have unit area in the [0,1] interval. The log ratio of density estimates for the non-PER and PER gene sets is shown in Figure 3D. We repeated this procedure to compare the distribution of minimum FPRs for correct classification of PER and non-PER “DE up” genes by the “DE up vs. rest” classifier. The log ratio of empirical density estimates for these two distributions is shown in Figure S16B.

**Description of CNN inputs**

For each knock-in condition, CNNs predicted the probability of methylation for a single CpG-context C from local sequence and metagene position information. Each CNN input was a 201 $\times$ 5 array. The first 4 columns of this array contained a one-hot encoding of the 201 bp nucleotide sequence centered on the C at which methylation was predicted and described in terms of the nucleotide content of the strand containing this C. The row entries of the last column contained the metagene coordinate for the nucleotide corresponding to the row.

**Architecture and training of CNN**

We constructed and trained CNNs using the python package Keras (https://keras.io). Our CNN models had one convolutional layer which slid 80 filters of size 6 $\times$5 along the rows of the 201 $\times$ 5 input array. Convolutional layer output was passed through a max-pooling layer with size 7 and stride 7 and then through a dropout layer with dropout rate 0.2 to reduce overfitting (13). Dropout layer output was passed to a fully connected layer of 40 neurons, then to another dropout layer with dropout rate of 0.2. Finally, the output of the second dropout layer was passed to a single output neuron encoding the methylation rate predicted by the network. Rectified linear (ReLu) activation functions were used throughout our network except at the output layer where we used a sigmoid activation to restrict predictions to the interval (0,1).

The quality of a CNN prediction, $\hat{p}$, for a single network input with $m$ methylated status observations in the total of $N$ observations was measured with the loss function

$$\ell(\hat{p}) = -\log\left[\binom{N}{m}\hat{p}^{m}(1-\hat{p})^{N-m}\right] = -m\log(\hat{p}) - (N-m)\log(1-\hat{p}) + \text{constant}$$

which is the negative log-likelihood of the $N$ independent observations given the predicted methylation probability $\hat{p}$. The loss for the training, validation, and test set was the sum of these individual losses over the elements of each set. We trained CNNs using the stochastic gradient descent optimizer with batch of size 1000, Nesterov momentum with parameter 0.9, and an initial learning rate $10^{-3}$ with decay factor $10^{-6}$. Training stopped when the validation loss did not decrease for more than 100 epochs.

**Calculation of prediction-based dissimilarity**

Our prediction-based dissimilarity (PBD) measured the extent to which observed differences in standardized methylation rates between conditions could be recapitulated by differences in standardized predictions of CNNs trained on the two conditions. The PBD between conditions A and B was calculated using the Pearson correlation

$$\mathrm{PBD}_{A,B} = \frac{\sum_i \left(\frac{p_{A,i} - \overline{p_A}}{\sigma_A} - \frac{p_{B,i} - \overline{p_B}}{\sigma_B}\right)\left(\frac{\hat{p}_{A,i} - \overline{\hat{p}_A}}{\hat{\sigma}_A} - \frac{\hat{p}_{B,i} - \overline{\hat{p}_B}}{\hat{\sigma}_B}\right)}{\sqrt{\sum_i \left(\frac{p_{A,i} - \overline{p_A}}{\sigma_A} - \frac{p_{B,i} - \overline{p_B}}{\sigma_B}\right)^2}\sqrt{\sum_i \left(\frac{\hat{p}_{A,i} - \overline{\hat{p}_A}}{\hat{\sigma}_A} - \frac{\hat{p}_{B,i} - \overline{\hat{p}_B}}{\hat{\sigma}_B}\right)^2}}.$$

In this equation, $i$ indexes the set of CpG-context Cs covered with at least one read in the six conditions considered, $p_{A,i}$ and $\hat{p}_{A,i}$ are respectively the mCpG rate and predicted methylation rate of the $i^{th}$ C in condition A, $\overline{p_A}$ and $\sigma_A$ are the mean and standard deviation of the mCpG rates for condition A, and $\overline{\hat{p}_A}$ and $\hat{\sigma}_A$ are the mean and standard deviation of the predicted methylation rates for condition A.

**Simulated annealing**

We used the simulated annealing (SA) technique to perform probabilistic optimization of CNN-predicted methylation rates over allowed input sequences. The SA algorithm samples sequences via a discrete-time inhomogeneous Markov chain with transition probabilities that depend on a cost function $J$ and a temperature parameter $T$. Sequences corresponding to lower $J$ values are sampled with greater probability than those corresponding to higher $J$ values, and the ratio of these probabilities increases as temperature decreases.

Several properties of SA make it useful for revealing sequence patterns important for network prediction. First, at bases where changes in nucleotide content produce notable changes in $J$, the distribution of A,C,T,G across samples will diverge from the uniform towards a distribution that favors nucleotides minimizing $J$. Second, this distribution does not need to favor a single nucleotide. For example, when two nucleotides produce low values of $J$ at a certain base, both will be preferentially sampled and the frequency of nucleotides will reflect the extent to which they lower $J$. Third, the divergence of nucleotide content in samples from a uniform distribution can be used to measure the relative importance of base positions to determining $J$. Sequence logos are frequently used to summarize nucleotide preference and relative importance of bases, so we used SA samples to make base-wise MLE estimates of nucleotide probabilities and constructed sequence logos using the Kullbeck-Leibler divergence from a uniform distribution (Figure 5A,B and 6A).

We here describe our implementation of SA using the notation of (14). To sample a sequence $\boldsymbol{x}$ maximizing methylation (SA maximization), we used the cost function $J(\boldsymbol{x}) = -y(\boldsymbol{x})$ where $y$ denotes the pre-activation of the output neutron of the CNN trained on that condition. The pre-activation of the output neuron is the neuron's value before applying the sigmoid function. For each CNN/condition pair, we initialized 50 instances of SA at the 50 elements of the test set predicted to have highest methylation. For each initialization, we executed the following pseudocode where $\boldsymbol{x}_n$ is a CNN input described by a length 201 array with entries A,C,G,T (metagene information in network input is not changed in SA and is therefore not described by this notation).

---

**Algorithm: SA**

---

**Inputs**: $\boldsymbol{x_0}, J(\boldsymbol{x}), d, N_{iter}, N_{sample}, N_{interval}$

**for** $n$ in $(1,2, \ldots N_{iter})$

$T = d/\log(n+1)$

$i, j \sim \text{Unif\_without\_replacement}(\{0, \ldots, 99\} \cup \{102, \ldots, 200\})$ ## Center CG is fixed

$\boldsymbol{x}_{proposed} = \boldsymbol{x}_{n-1}$

$(\boldsymbol{x}_{proposed})_i \sim \text{Unif}(\{A, C, G, T\} - \{(\boldsymbol{x}_{n-1})_i\})$

$(\boldsymbol{x}_{proposed})_j \sim \text{Unif}(\{A, C, G, T\} - \{(\boldsymbol{x}_{n-1})_j\})$

$u \sim \text{Unif}([0,1])$

**if** $\exp\left(-\frac{J(\boldsymbol{x}_{proposed}) - J(x_{n-1})}{T}\right) > u$ ## Accept (1)

$x_n = x_{proposed}$

**else** ## Reject

$x_n = x_{n-1}$

**Return** $\{ x_n : (n > N_{iter} - (N_{sample})(N_{interval})) \textbf{ and } n \bmod N_{interval} = 0 \}$

The choice of proposing two mutations at each iteration is specific to our implementation. We also did not propose mutations at the center CG because the sequence content of this dinucleotide was fixed across all elements of training, test and validation sets. All conditions used parameters $N_{iter} = 8 \times 10^6$, $N_{sample} = 2 \times 10^4$ and $N_{interval} = 2$. The parameter $d$, related to the initial temperature, was chosen in a condition-specific manner described below. We used the same algorithm and parameters to sample sequences predicted to minimize methylation in each condition (SA minimization), except that we initialized at the 50 elements of the test set predicted to have lowest methylation and set $J(\boldsymbol{x}) = y(\boldsymbol{x})$. Figure S18A,B plots the minimum of $J(\boldsymbol{x})$ across previous iterations versus number of SA iterations. Data points represent averages of minima for 50 initializations and demonstrate convergence in the sense of negligible improvement in the quality of cost function minima after the first several million iterations.

**Choice of the temperature parameter $d$ in simulated annealing**

Our choice of $d$ parameter values specific to each CNN/condition pair was motivated by the empirical observation that the widths of SA objective function distributions varied between CNNs (Figure S18C,D). Differences in widths cause differences in probability of accepting proposed high cost states. For instance, for fix $d$, $\boldsymbol{x}_{n-1}$ and some $0 < \alpha < 1$, the transformation $J(\boldsymbol{x}) \to \alpha J(\boldsymbol{x})$ in equation (1), which corresponds to narrowing the distribution of SA objective function values, increases the acceptance probability of $\boldsymbol{x}_{\text{proposed}}$ when $J(\boldsymbol{x}_{proposed}) > J(x_{n-1})$.

To choose a value of $d$ suitable for the distribution of each objective function, we considered the heights of communication of local minima. A state $\boldsymbol{x}$ of the Markov chain is said to communicate with the set $X^*$ of global minima at height $h$ if there exists a path (a sequence of proposed mutations) from $\boldsymbol{x}$ to an element of $X^*$ such that the maximum value of $J$ along the path is at most $J(\boldsymbol{x}) + h$ (14). Communication heights are important in theoretical analysis of SA convergence which shows that, provided all states communicate with $X^*$ at height $d$, our cooling schedule yields convergence to a global minimum of $J$ in the limit of infinite simulation steps (14). Although we were not interested in obtaining true global minima (as fluctuation in sequence content at unimportant bases is essential to biological interpretation), we still needed to ensure that Markov chains could transition from shallow to deeper basins of the cost function. We therefore selected condition-specific values of $d$ by estimating a characteristic minimum communication height for sequences near local minima of $J$.

In the case of SA maximization, estimation for each condition used the following procedure: first, we supplemented each of the 50 test set inputs predicted to have highest methylation with 1,000 inputs sampled uniformly and without replacement from the union of training, validation, and test sets. Metagene information was fixed during SA, so we removed it as a source of variation in $J(\boldsymbol{x})$ by setting the metagene values of all 1000 sampled inputs to those of the corresponding high methylation input. Next, we estimated the fraction of inputs near strong local minima by plotting the inverse cumulative distribution function of $J(\boldsymbol{x})$ for the pooled 50,0000 samples, and observed a transition from rapid increase in $J(\boldsymbol{x})$ for inputs in the lowest percentiles of $J(\boldsymbol{x})$ to moderate increase in $J(\boldsymbol{x})$ for the bulk of the remaining inputs (Figure S19A). Using the 1$^{st}$ percentile of $J(\boldsymbol{x})$ as a threshold we assigned inputs below the threshold to a set termed "basin points" and the remainder of inputs to a set termed "bulk points". Finally, we made the following assumptions:

(1) if a basin point is not in $X^*$, then the Markov chain must pass through the set of bulk points to reach $X^*$ and
(2) for paths from basin points $\boldsymbol{x}$ that attain $\boldsymbol{x}$'s minimum communication height, the change in $J$ among bulk points on the path is negligible compared to the increase in $J$ required to enter the set of bulk points.

With these assumptions we replaced the problem of estimating minimum communication heights with that of estimating differences in $J$ between basin and bulk points. We estimated these differences with a Monte Carlo approach

$$d = \text{percentile}\left(1, \bigcup_{i=1}^{50} \left\{ J_{i_{(k)}} - J_{i_{(1)}} : 2 \le k \le 1001 \right\}\right)$$

where $i$ indexes the 50 test set inputs with highest predicted methylation, $J_{i_{(k)}}$denotes the kth smallest value of $J$ among the union of the 1000 samples associated with test set input $i$ and the input $i$ itself, and where $\text{percentile}(1, S)$ denotes the 1$^{st}$ percentile of the set $S$. Figure S19B illustrates this distribution and the value of $d$ selected for SA maximization in each condition.

For SA minimization, we used the same procedure, except that we set $J(\boldsymbol{x}) = y(\boldsymbol{x})$ and considered the 50 test set elements with lowest predicted methylation (Figure S19C,D).

**Clustering of SA results**

Among the 300 sets of SA samples, sequence preferences were consistently most pronounced at the nucleotides flanking the CpG site. We therefore used hierarchical clustering to organize SA results for each condition/initialization based on local sequence motifs and summarized the sequence preferences of each cluster containing more than one set of samples with a sequence logo. We applied this procedure twice: once for SA sample subsequences starting 4 bp 5' and ending 8 bp 3' of the central CpG C (fixed across SA samples) and once for the full 201 bp of all SA samples.

Specifically, to compare sequence preferences near C's with maximum predicted methylation, we first calculated base-wise single nucleotide frequencies across samples collected for each initialization of each condition. Next, we measured distances between all initialization pairs as the sum of base-wise Jenson-Shannon divergence

$$d(P,Q) = \sum_{i=96}^{109} \mathrm{JSD}(P_i || Q_i) \,.$$

Here, $P$ and $Q$ are $14 \times 4$ vectors of base-wise single nucleotide frequencies for sets of samples from two initializations and $\mathrm{JSD}(P_i || Q_i)$ is the Jenson-Shannon divergence between single nucleotide frequencies at base $i$. Finally, we used these pairwise distances to perform hierarchical clustering with complete linkage, assigning SA results for each initialization to a flat cluster by applying a distance threshold to the clustering diagram. To summarize the sequence preferences represented by each cluster, we aggregated SA samples for all initializations in the cluster, recalculated base-wise single nucleotide frequencies, and, for all clustering containing the results of more than one initialization, generated sequence logos using the code available at https://github.com/kundajelab/deeplift/blob/master/deeplift/visualization/viz_sequence.py (15).

We used the same procedure to analyze SA minimization results and compare sequence biases near C's with minimum predicted methylation. To cluster SA maximization and minimization results spanning the full 201 bp of network input sequences, we made two modifications to the above procedure. First, we allowed the base-wise sum on Jenson-Shannon divergences to run from 0 to 200. Second, we assigned SA results to flat clustering using the inconsistent statistic criterion of scipy (https://www.scipy.org/) rather than a distance threshold.

**Validation of SA motifs**

To verify that the motifs extracted from CNNs by SA maximization and SA minimization actually corresponded to sites of high and low methylation *in vivo*, respectively, we identified CpG sites with flanking sequence matching one of our SA motifs and compared the observed methylation at these sites with the observed methylation at CpGs not matching our motifs. For this purpose, we first constructed position-specific scoring matrices (PSSMs) summarizing the total sequence content of each of our 8 clusters of SA maximization sequences (Figure 5A) and each of our 10 clusters of SA minimization sequences (Figure 5B). Next, we used FIMO (16) (version 5.0.1) with default parameters to scan for each of these 18 motifs genome wide. We identified hits using the FIMO default p-value threshold of $10^{-4}$. For each knock-in condition, we filtered out hits with zero WGBS coverage and only considered mCpG rates at PSSM hits containing the central CpG. Finally, for each condition, we used the Mann-Whitney U test to compare the mCpG rates at retained motif hits with those at genome-wide CpG cytosines in the complement of the PSSM hits and with non-zero WGBS coverage.

#### Fourier analysis of periodic trinucleotides

We quantified the periodicity of trinucleotides in individual sequences sampled by SA by applying discrete fast Fourier transform (DFFT) to arrays of indicator variables for the presence of each trinucleotide at successive sequence positions. Sequences were analyzed individually by constructing a length-$199$ binary array indicating trinucleotide presence or absence, with the center 19 entries corresponding to the core motif region set to zero. We averaged the resulting DFFT amplitudes across the $10^6$ individual sequences sampled by SA ($2 \times 10^4$ samples per SA iteration and 50 iterations per condition) and plotted the results for the trinucleotides with greatest 10.5 bp amplitude in each condition (Figure 6B; Figure S25A,S26A).

We next identified the phase of trinucleotide periodicity between the 5' and 3' sides of the central CpG as follows: we tested different phase relationship by progressively inserting or deleting zeros into the zeroed-out central regions of the constructed arrays; the number of zeros inserted or deleted corresponded the phase shift of 3' subsequences in bp. For each shifted sequence, we calculated the Fourier amplitude of the binary vector by taking the amplitude of the dot production of the binary vector with a discrete complex exponential function with period between 5 and 20 bp. The resulting amplitudes were averaged over the $10^6$ individual sequences sampled for each DNMT knock-in condition (Figure 6C ; Figure S26B,S26B).

#### MI calculation

Mutual information (MI) between observations of binary methylation status was calculated as a function of distance between paired CpGs. Distances were binned with a width of 30 bp and with bin lower bounds ranging from 0 to 180 bp with a stride of 3 bp. In each bin, we paired any two methylation status observations that were obtained from the same pair-end read and that corresponded to CpG sites separated by a distance within the bin; we let $J$ be an index set labeling all distinct individual CpG methylation status observations appearing in such pairs. Denoting the $k^{th}$ pair as $(j_{k,x}, j_{k,y}) \in J \times J$, with $j_{k,x}$ 5' to $j_{k,y}$, we obtained $J_x = \{j_{k,x}\}_k$ and $J_y = \{j_{k,y}\}_k$, where redundant elements were removed. Next, we defined a function $m$ binarizing an indexed observation as

$$m(j) = \begin{cases} 1, \text{if the } j^{\text{th}} \text{ observation was methylated} \\ 0, \text{otherwise} \end{cases}.$$

Using this definition, we estimated the joint probability of methylation status random variables $X$ and $Y$ satisfying the binned distance criterion by

$$\hat{P}(X = a, Y = b) = \frac{\sum_k I[m(j_{k,x}) = a] I[m(j_{k,y}) = b]}{\sum_k 1},$$

where $a \in \{0,1\}$, $b \in \{0,1\}$, and $I[\cdot]$ is the indictor for an event. Similarly, the marginal probabilities of $X$ and $Y$ were estimated by

$$\hat{P}(X = a) = \frac{\sum_{j \in J_x} I[m(j) = a]}{\sum_{j \in J_x} 1}$$

$$\hat{P}(Y = a) = \frac{\sum_{j \in J_y} I[m(j) = a]}{\sum_{j \in J_y} 1}.$$

Using these estimates, we calculated the MI of $X$ and $Y$ as

$$MI(X;Y) = \sum_{a \in \{0,1\}} \sum_{b \in \{0,1\}} \hat{P}(X = a, Y = b) \log\left(\frac{\hat{P}(X = a, Y = b)}{\hat{P}(X = a)\hat{P}(Y = b)}\right),$$

and the Shannon entropy of X and Y as

$$H(X) = -\sum_{a \in \{0,1\}} \hat{P}(X = a)\, log\big(\hat{P}(X = a)\big)$$

$$H(Y) = -\sum_{a \in \{0,1\}} \hat{P}(Y = a)\, log\big(\hat{P}(Y = a)\big).$$

Finally, MI was normalized by the sum of entropy as following:

$$Normalized\, MI = 2\frac{MI(X;Y)}{H(X) + H(Y)}.$$

We used a simulated control to check that the oscillation in normalized MI (Figure 6E; Figure S27B) was not due to periodicity in marginal methylation rates at individual CpG sites. Joint methylation status was simulated by independently sampling from the mCpG rate of CpG sites composing a pair (Supplementary Methods: "Empirical Bayes approach for estimating mCpG rate").

**MNase-seq data processing**

Reads were mapped using Bowtie2 (17) with the options --end-to-end --sensitive --score-min L,-1.5,-0.3. Reads with mapping quality lower than 13 were filtered out. Technical replicates from Liachko *et al*. (18) were combined. Nucleosome occupancy was defined as the read coverage after extending the single-end reads from 5’ to 3’ end by 147 bp, and calculated using bedtools genomecov (19) with the options -bg -split -fs 147.

**Calculation of nucleosome occupancy and dyad calling**

We used NSeq with default parameters to call positioned nucleosome dyad positions (20). NSeq input was the combination of aligned reads from replicate MNase-seq experiments (18). Nucleosome centers identified by NSeq at 1% FDR were used to calculated dyad aligned mCpG rates.

### Supplementary Tables:

| Sample | Total number of reads | Alignment rate |
|---|---|---|
| 1A-3L s1 | 24495800 | 0.73 |
| 1A-3L s2 | 24639440 | 0.79 |
| 1A-3L s3 | 24628310 | 0.77 |
| 1A s1 | 20040150 | 0.77 |
| 1A s2 | 24497428 | 0.77 |
| 1A s3 | 24460734 | 0.72 |
| 3A2-3L s1 | 24645136 | 0.76 |
| 3A2-3L s2 | 24373744 | 0.74 |
| 3A2-3L s3 | 24465856 | 0.76 |
| 3A2 s1 | 20040600 | 0.76 |
| 3A2 s2 | 25655498 | 0.74 |
| 3A2 s3 | 24451724 | 0.71 |
| 3A1-3L s1 | 24587878 | 0.77 |
| 3A1-3L s2 | 24518532 | 0.77 |
| 3A1-3L s3 | 24624966 | 0.76 |
| 3A1 s1 | 20015888 | 0.74 |
| 3A1 s2 | 24444154 | 0.76 |
| 3A1 s3 | 24480888 | 0.75 |
| 3B1 s1 | 20128490 | 0.77 |
| 3B1 s2 | 24457414 | 0.70 |
| 3B1 s3 | 24579522 | 0.76 |
| 3L s1 | 20114228 | 0.76 |
| 3L s2 | 24627598 | 0.77 |
| 3L s3 | 24625042 | 0.78 |
| Control s1 | 24468368 | 0.75 |
| Control s2 | 24314466 | 0.77 |
| Control s3 | 24440600 | 0.76 |
| 3B1-3L d1 s1 | 24498878 | 0.77 |
| 3B1-3L d2 s1 | 24589936 | 0.76 |
| 3B1-3L d3 s1 | 25996380 | 0.75 |
| 3B1-3L d4 s1 | 25926784 | 0.72 |
| 3B1-3L d1 s2 | 24472072 | 0.72 |
| 3B1-3L d2 s2 | 24501640 | 0.78 |
| 3B1-3L d3 s2 | 25947748 | 0.76 |
| 3B1-3L d4 s2 | 25921332 | 0.71 |
| 3B1-3L d1 s3 | 24425484 | 0.75 |
| 3B1-3L d2 s3 | 25958024 | 0.75 |
| 3B1-3L d3 s3 | 25897668 | 0.70 |
| 3B1-3L d4 s3 | 25836792 | 0.73 |

**Table S2**: Summary of RNA-seq data. Alignment rate is the fraction of read pairs with accepted alignments. The suffixes s1,s2, and s3 indicate biological replicate sample index.

| Sample | | Alignment rate | | | Bisulfite conversion rate | CpG non-conversion rate | CpG methylation rate | Coverage |
|---|---|---|---|---|---|---|---|---|
| | | Overall | PE | R1 | | | | |
| 1A-3L | s1 | 0.45 | 0.41 | 0.14 | 0.9953 | 0.0043 | 0.0039 | 16.4 |
| | s2 | 0.43 | 0.35 | 0.27 | 0.9954 | 0.0042 | 0.0042 | 15.9 |
| 1A | s1 | 0.53 | 0.49 | 0.17 | 0.9952 | 0.0046 | 0.0042 | 25.0 |
| | s2 | 0.33 | 0.28 | 0.15 | 0.9953 | 0.0044 | 0.0040 | 7.7 |
| 3A2-3L | s1 | 0.50 | 0.42 | 0.26 | 0.9957 | 0.0042 | 0.0489 | 15.9 |
| | s2 | 0.41 | 0.33 | 0.22 | 0.9955 | 0.0043 | 0.0770 | 14.8 |
| 3A2 | s1 | 0.59 | 0.56 | 0.14 | 0.9953 | 0.0043 | 0.0199 | 27.8 |
| | s2 | 0.33 | 0.30 | 0.09 | 0.9959 | 0.0038 | 0.0243 | 11.9 |
| 3A1-3L | s1 | 0.35 | 0.33 | 0.06 | 0.9951 | 0.0049 | 0.0487 | 10.9 |
| | s2 | 0.41 | 0.31 | 0.30 | 0.9961 | 0.0038 | 0.0460 | 14.9 |
| 3A1 | s1 | 0.45 | 0.40 | 0.18 | 0.9948 | 0.0051 | 0.0125 | 21.0 |
| | s2 | 0.29 | 0.24 | 0.13 | 0.9953 | 0.0042 | 0.0148 | 8.1 |
| 3B1 | s1 | 0.59 | 0.55 | 0.15 | 0.9955 | 0.0042 | 0.0049 | 27.6 |
| | s2 | 0.44 | 0.34 | 0.32 | 0.9956 | 0.0042 | 0.0051 | 16.3 |
| 3L | s1 | 0.50 | 0.42 | 0.29 | 0.9957 | 0.0040 | 0.0042 | 23.9 |
| | s2 | 0.46 | 0.37 | 0.29 | 0.9955 | 0.0044 | 0.0038 | 17.0 |
| Control | s1 | 0.53 | 0.45 | 0.31 | 0.9955 | 0.0045 | 0.0040 | 19.3 |
| | s2 | 0.46 | 0.36 | 0.31 | 0.9957 | 0.0042 | 0.0040 | 16.7 |
| 3B1-3L d1 | s1 | 0.44 | 0.38 | 0.21 | 0.9954 | 0.0042 | 0.0209 | 15.9 |
| | s2 | 0.42 | 0.39 | 0.10 | 0.9953 | 0.0049 | 0.0205 | 12.8 |
| 3B1-3L d2 | s1 | 0.63 | 0.60 | 0.19 | 0.9951 | 0.0048 | 0.0070 | 23.3 |
| | s2 | 0.59 | 0.55 | 0.18 | 0.9951 | 0.0045 | 0.0076 | 21.9 |
| 3B1-3L d3 | s1 | 0.56 | 0.38 | 0.58 | 0.9955 | 0.0042 | 0.0107 | 20.5 |
| | s2 | 0.51 | 0.35 | 0.51 | 0.9956 | 0.0043 | 0.0125 | 19.2 |
| 3B1-3L d4 | s1 | 0.54 | 0.40 | 0.45 | 0.9944 | 0.0053 | 0.0115 | 20.0 |
| | s2 | 0.57 | 0.45 | 0.45 | 0.9946 | 0.0052 | 0.0126 | 20.8 |

**Table S3**: Summary of whole-genome bisulfite sequencing (WGBS) data. “Alignment rate: Overall” is the number of single-end reads aligned either as a read pair or as a single read, divided by the total number of single reads for which alignment was attempted. “Alignment rate: PE” is the number of read pairs aligned in initial paired-end alignment divided by number of read pairs for which paired-end alignment was attempted. “Alignment rate: R1” is the number of first reads that did not align as pairs but which aligned as single reads, divided by the number of first reads that did not align as pairs. “Bisulfite conversion rate” is the fraction of cytosines converted by bisulfite treatment in the unmethylated lambda phage spike-in DNA. “CpG non-conversion rate” is the fraction of CpG context cytosines that were not converted by bisulfite treatment of the lambda phage spike-in DNA. This statistic provides an estimate of CpG methylation levels attributable to incomplete conversion of unmethylated cytosines. “CpG methylation rate” is the CpG-context methylation rate of the *K. phaffii* genome. “Coverage” is the total number of bases mapped (in paired or single-end alignments) divided by the genome size.

| Condition | Mean mCpG rate | | p-value |
|---|---|---|---|
| | w/ motif | w/o motif | Mann-Whitney U test |
| 3A1 | 0.0316 | 0.0132 | 1.95E-144 |
| 3A2 | 0.0613 | 0.0213 | 7.04E-191 |
| 3B1 | 0.0059 | 0.0050 | 2.66E-14 |
| 3A1-3L | 0.1240 | 0.0470 | 4.57E-173 |
| 3A2-3L | 0.1268 | 0.0626 | 9.68E-22 |
| 3B1-3L d1 | 0.0410 | 0.0205 | 2.19E-65 |

**Table S6:** Validation of motifs identified by SA to be preferred by DNMTs.

| Condition | Mean mCpG rate | | p-value |
|---|---|---|---|
| | w/ motif | w/o motif | Mann-Whitney U test |
| 3A1 | 0.0094 | 0.0133 | 1.11E-21 |
| 3A2 | 0.0102 | 0.0215 | 2.03E-78 |
| 3B1 | 0.0046 | 0.0050 | 6.50E-05 |
| 3A1-3L | 0.0189 | 0.0474 | 2.16E-99 |
| 3A2-3L | 0.0204 | 0.0629 | 7.69E-168 |
| 3B1-3L d1 | 0.0148 | 0.0205 | 3.94E-12 |

**Table S7:** Validation of motifs identified by SA to be avoided by DNMTs.

### Supplementary Figures:

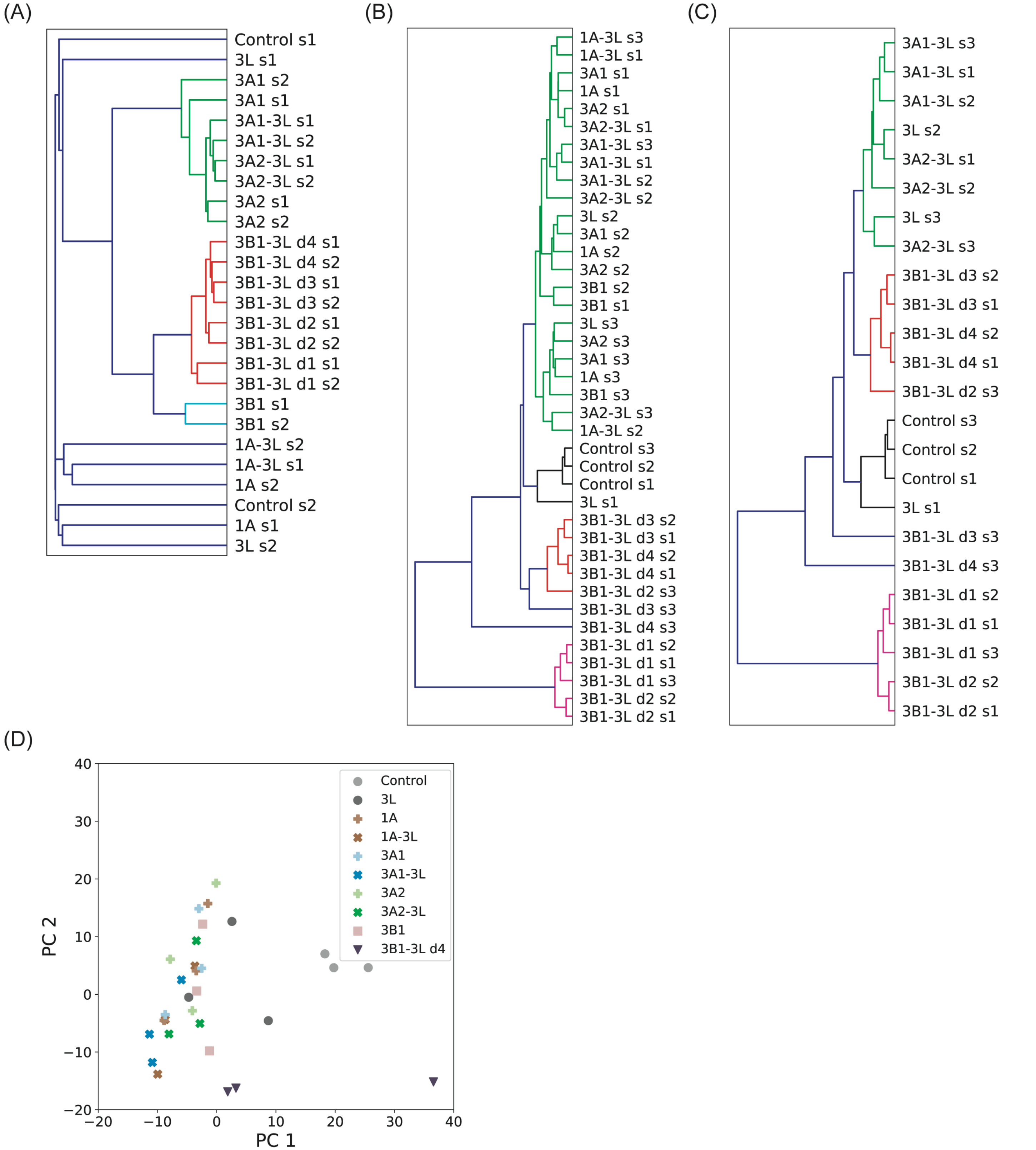


**Figure S1: Clustering of WGBS samples and RNA-seq samples. (A)** Hierarchical clustering of WGBS samples based on Pearson correlation between window average methylation rates (window size of 4 kb). Clustering used average linkage. **(B)** Hierarchical clustering of RNA-seq samples based on Pearson correlation between batch-corrected log-scale expression (Supplementary Methods: “Gene expression PCA”). Clustering used average linkage. **(C)** Similar to (B), including only double knock-in, 3L, and Control samples. **(D)** Principal component analysis of transcriptome-wide mRNA levels, excluding 3B1-3L d1-d3.

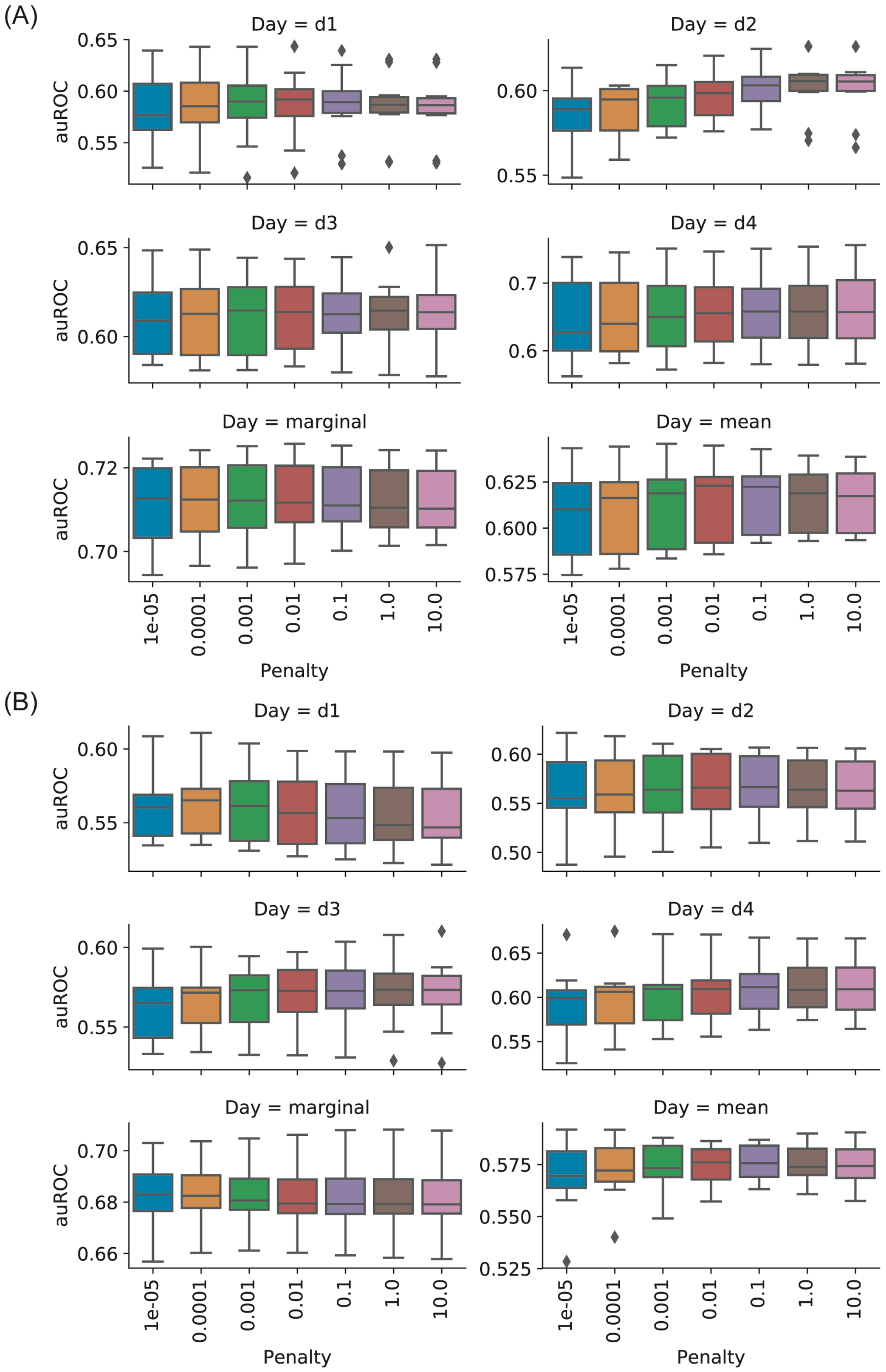


**Figure S2: Distributions of auROC** for **(A)** the “DE down vs. rest” classifier and **(B)** the “DE up vs. rest” classifier across test sets in 10-fold cross validation. Panels labeled by “Day = d*n*” shows the test set performance restricted to prediction of gene differential expression status on time-course day *n*. Panel labeled “Day = marginal” shows the test set performance using predictions of gene differential expression status on all days. Panel labeled “Day = mean” shows the distributions, across cross-validation folds, of the mean of performance statistics calculated by restricting to prediction on individual time-course days (Supplementary Methods: “Choice of logistic regression smoothness penalty”).

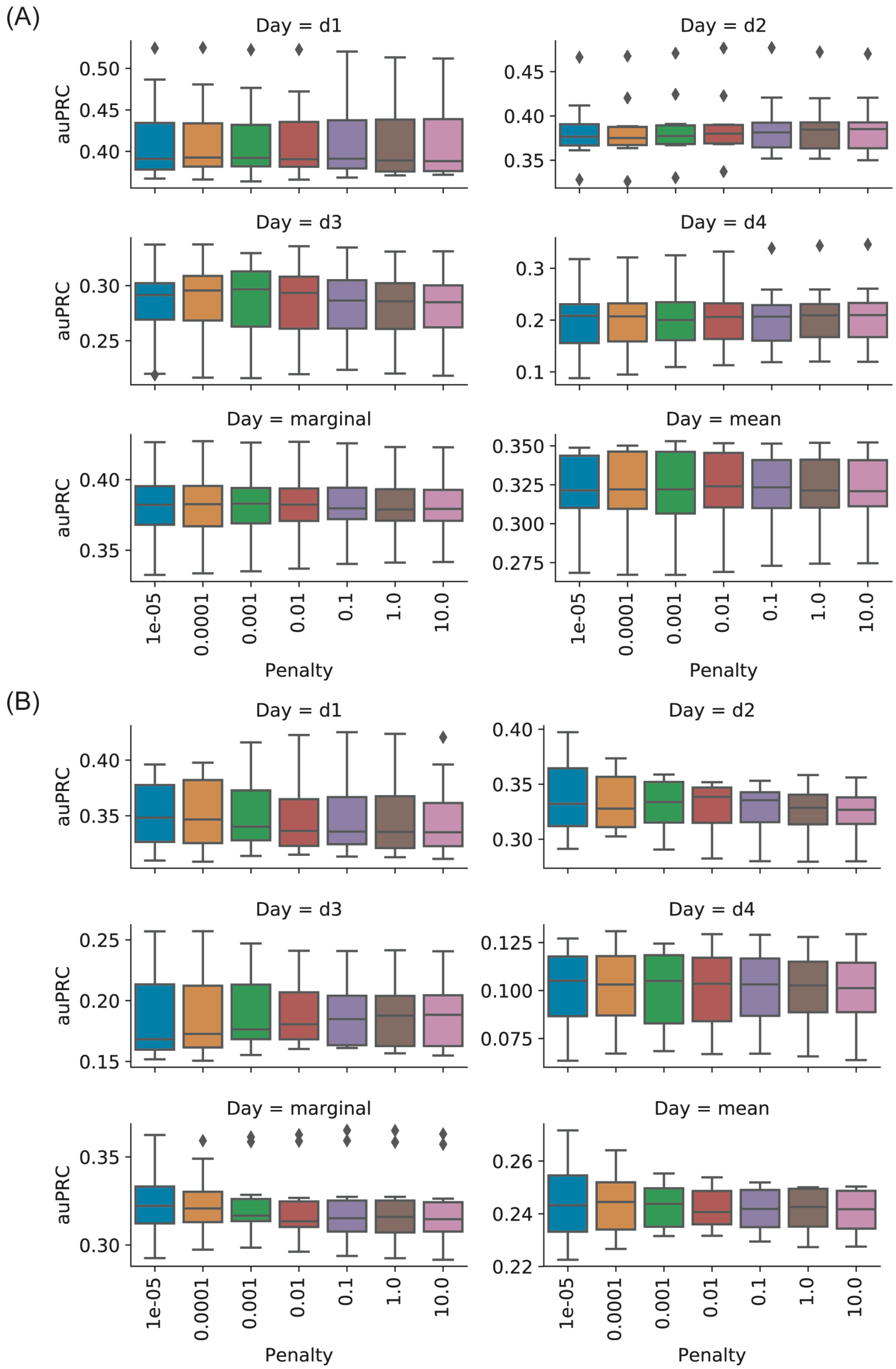


**Figure S3: Distributions of auPRC** for **(A)** the “DE down vs. rest” classifier and **(B)** the “DE up vs. rest” classifier across test sets in 10-fold cross validation. Panels labeled by “Day = d*n*” shows the test set performance restricted to prediction of gene differential expression status on time-course day *n*. Panel labeled “Day = marginal” shows the test set performance using predictions of gene differential expression status on all days. Panel labeled “Day = mean” shows the distributions, across cross-validation folds, of the mean of performance statistics calculated by restricting to prediction on individual time-course days (Supplementary Methods: “Choice of logistic regression smoothness penalty”).

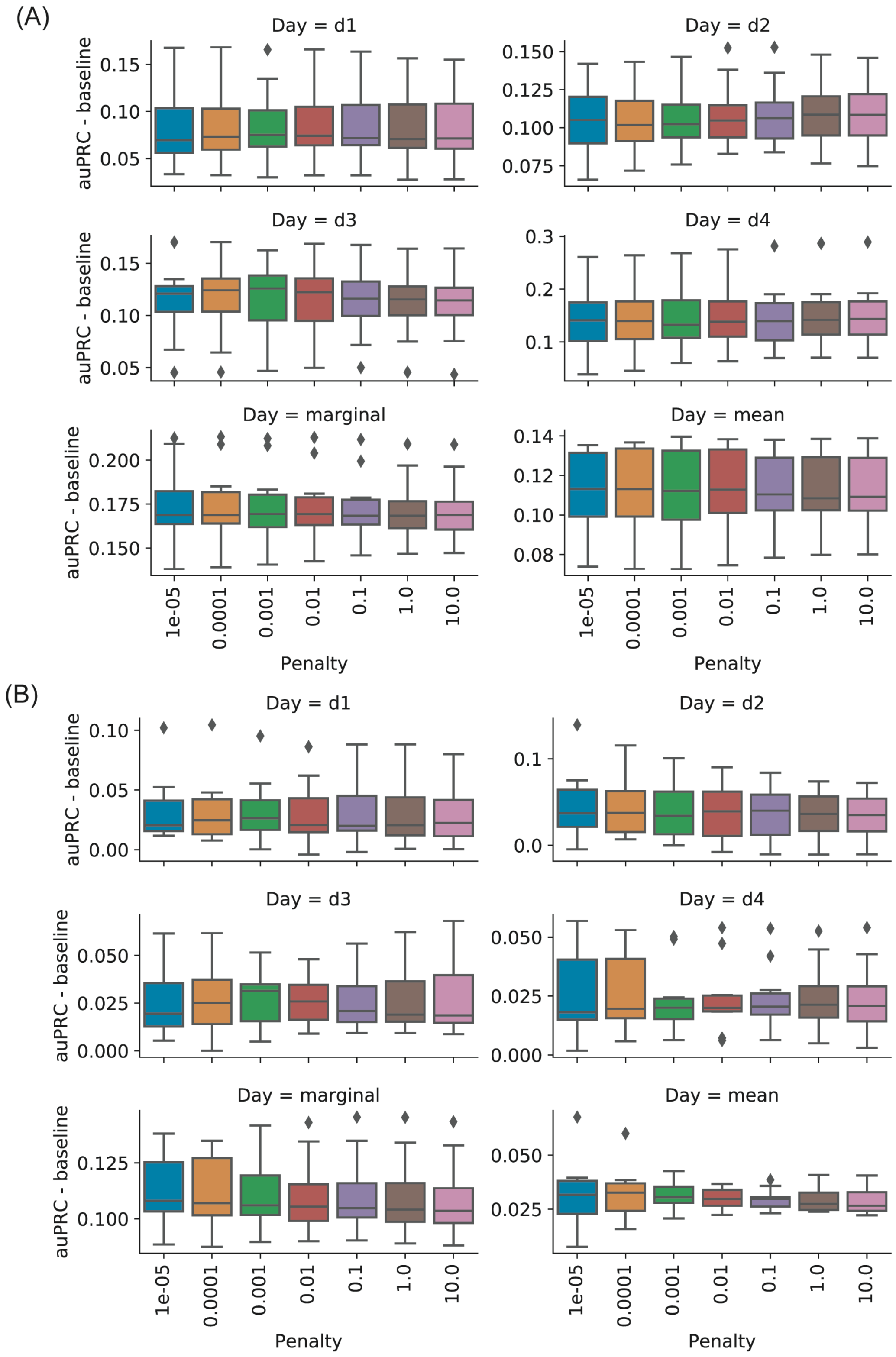


**Figure S4: Distributions of auPRC minus expected auPRC of a baseline classifier**, for **(A)** the "DE down vs. rest" classifier and **(B)** the "DE up vs. rest" classifier, across test sets in 10-fold cross validation. Panels labeled by "Day = d*n*" shows the test set performance restricted to prediction of gene differential expression status on time-course day *n*. Panel labeled "Day = marginal" shows the test set performance using predictions of gene differential expression status on all days. Panel labeled "Day = mean" shows the distributions, across cross-validation folds, of the mean of performance statistics calculated by restricting to prediction on individual time-course days (Supplementary Methods: "Choice of logistic regression smoothness penalty").

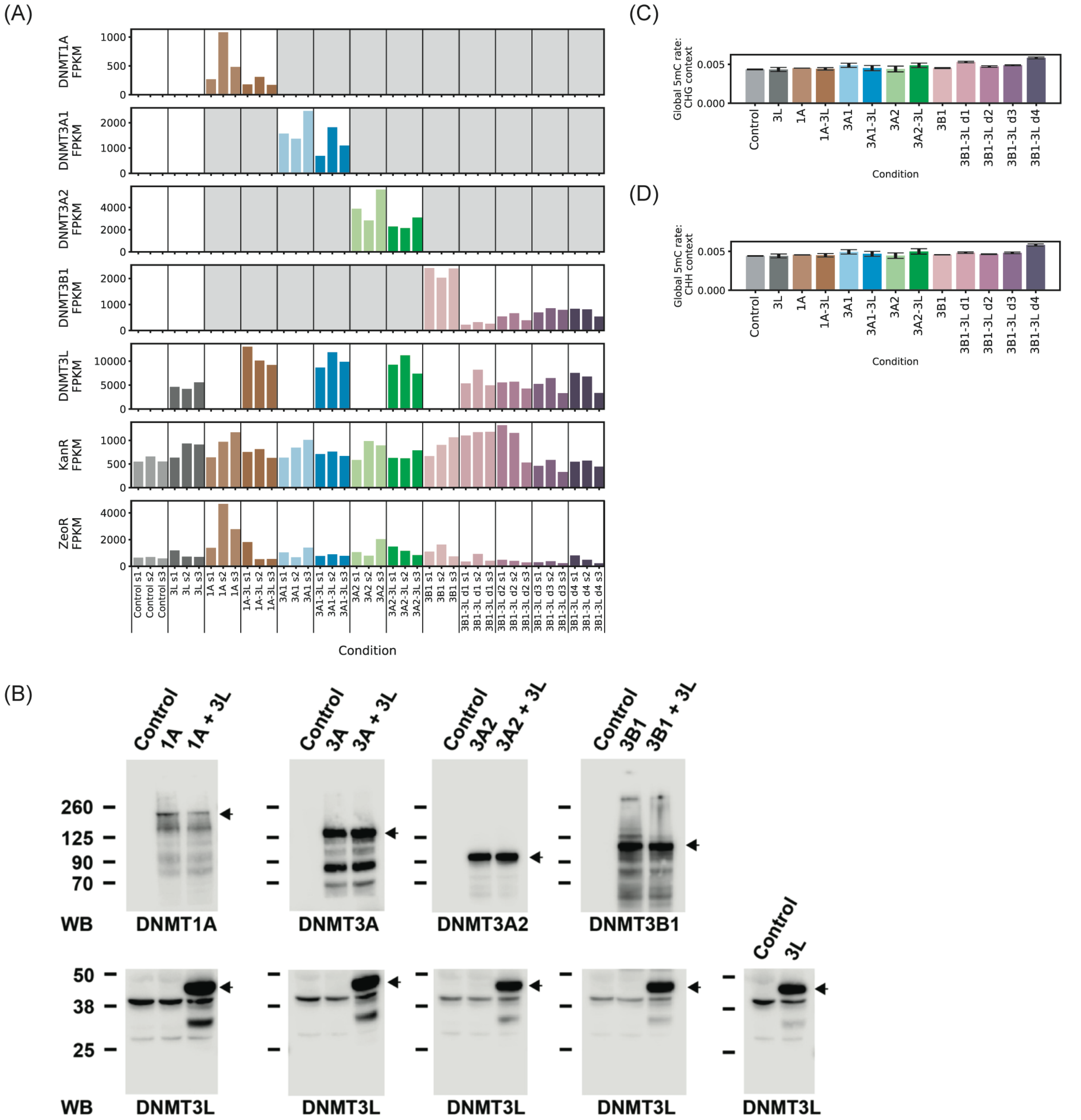


**Figure S5: Knocked-in DNMT expression and CHG- and CHH-context methylation. (A)** Expression of knocked-in DNMT genes by sample (in units of fragments per kilobase transcript per million (FPKM)). For each knock-in gene, grey regions indicate samples for which the knock-in sequence was excluded from the reference genome used for alignment. Knock-in genes were excluded from reference genomes to prevent multiple alignment of reads to similar DNMT isoforms. **(B)** Protein expression of DNMT in the different strains created was analyzed by Western blot: DNMT1A (1st column, top row), DNMT3A1 (2nd column, top row), DNMT3A2 (3rd column, top row), DNMT3B1 (4th column, top row) and 3L (bottom row). The black arrows indicate full-length products. **(C)** Fraction of 5mC in CHG context for each experimental condition. Bar height is the average of two replicates, and error bars show the maximum and minimum across replicates. **(D)** Same as (C), but for 5mC in CHH context.

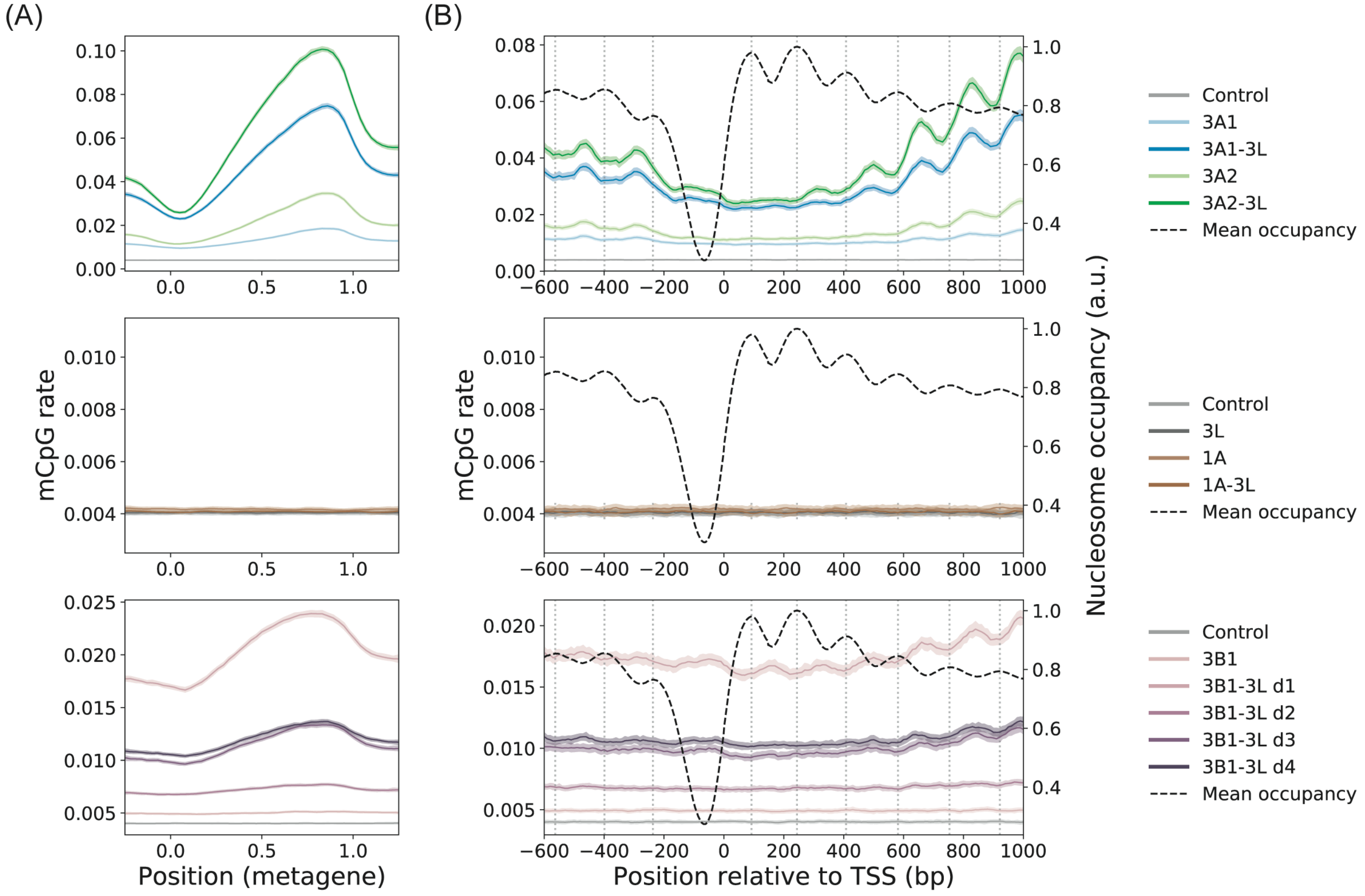


**Figure S6**: **(A)** Metagene plot of mCpG rates averaged across *K. phaffii* genes. CpG-context cytosines were assigned to a metagene coordinate, the distance downstream of the gene transcription start site (TSS) scaled by gene length. Solid lines indicate average mCpG rate of cytosines in a collection of sliding metagene coordinate windows (widow width of 0.15). Shading indicates 95% Bayesian credible intervals using empirical Bayes beta priors. **(B)** TSS-aligned mCpG rates and nucleosome occupancy averaged across *K. phaffii* genes. Lines and shaded credible intervals were calculated using a sliding window method similar to (A) (widow width of 50 bp).

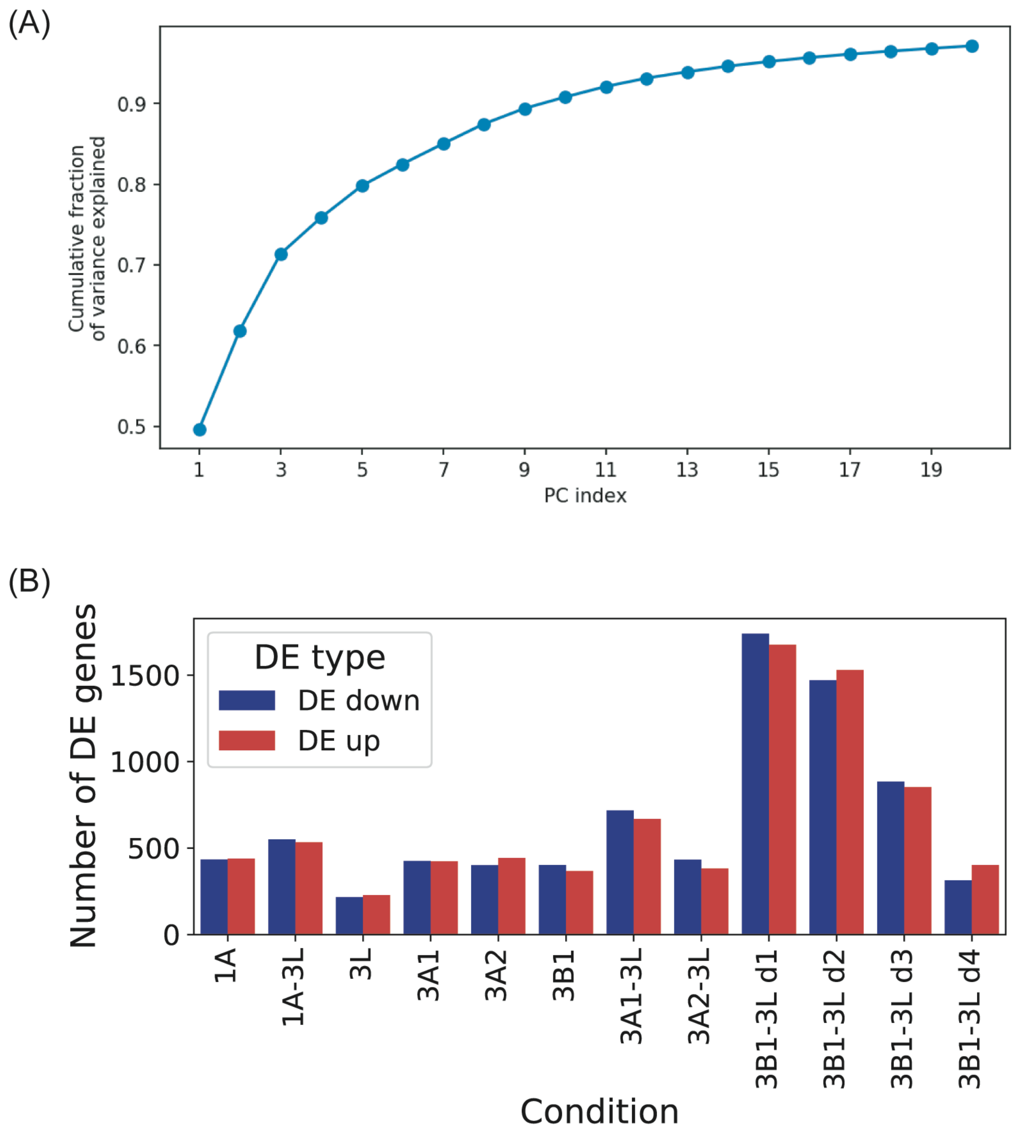


**Figure S7: Fraction of variance explained in PCA and number of differentially expressed (DE) genes for RNA-seq data**. **(A)** Fraction of variance explained by principal components in principal components analysis of gene expression for all samples. **(B)** Number of DE genes (5% FDR) in each condition.

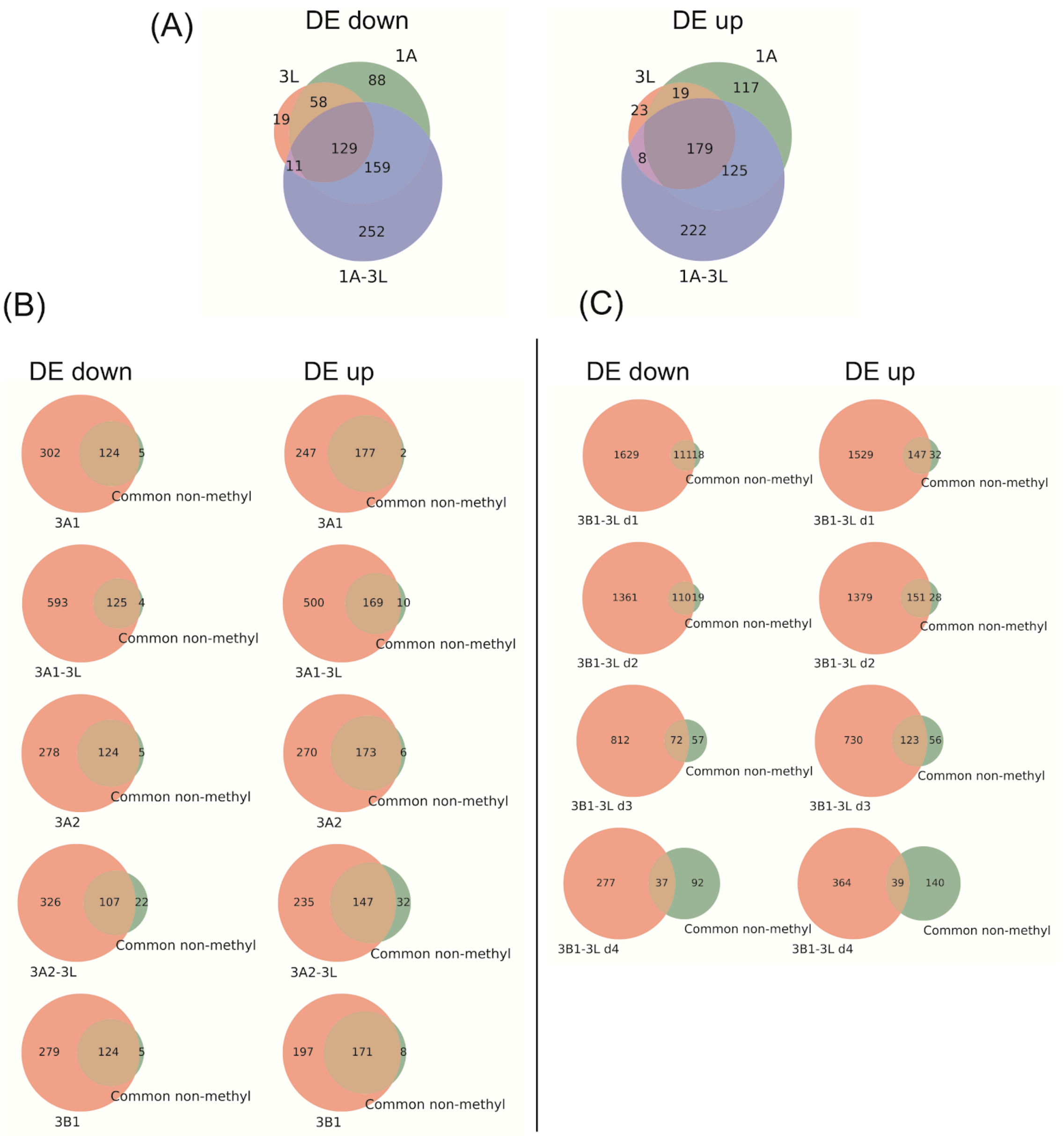


**Figure S8: Venn Diagrams of differentially expressed (DE) genes between conditions.** (A) DE genes in non-methylating 3L, 1A, and 1A-3L conditions overlap significantly. (B) DE genes common to 3L, 1A and 1A-3L conditions are also differentially expressed in *de novo* DNMT knock-in conditions. (C) Same as (B), but for 3B1-3L time-course data.

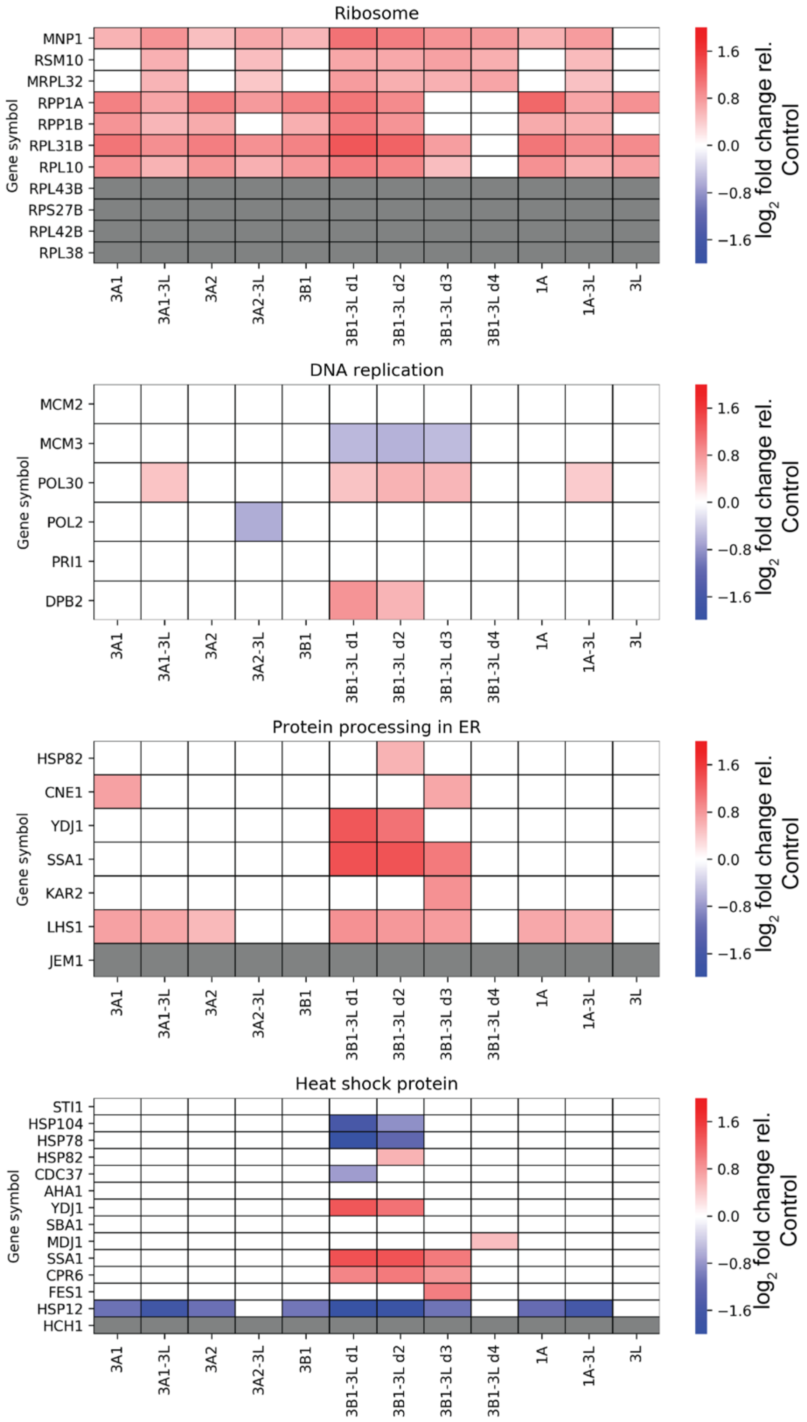


**Figure S9: Differential expression status of genes related to Unfolded Protein Response.** High expression of recombinant proteins may lead to increased expression of ER-related proteins and heat shock proteins, while suppressing ribosome and DNA replication genes potentially due to reduced cell growth. Our data do not show this trend. Grey cells in the table indicate genes that were missing in our data. To aid in visualization, minimum and maximum values of color maps are set at -2 and +2, respectively.

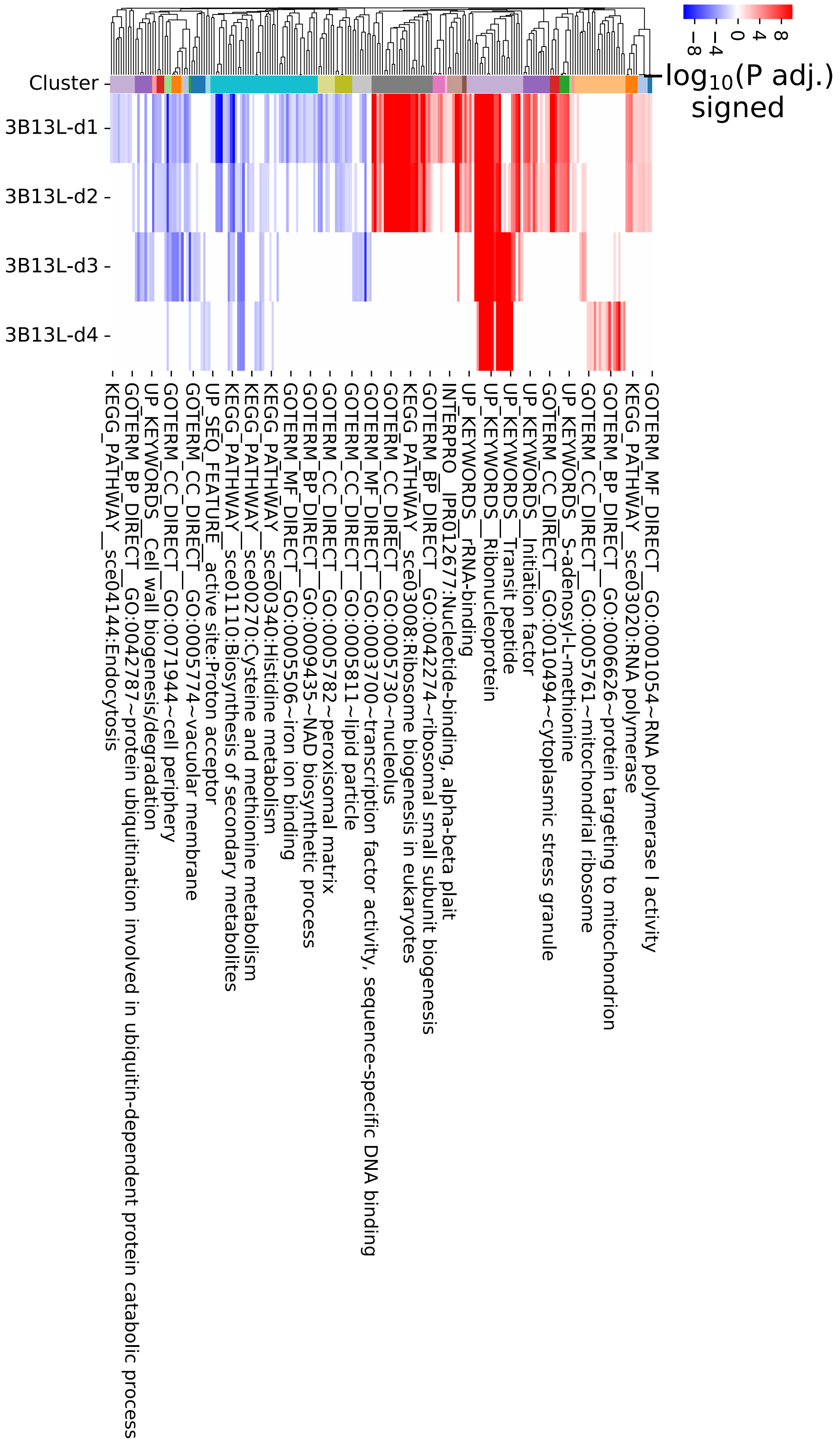


**Figure S10: Clustering of GO terms enriched in "DE up" or in "DE down" genes on any time-course day.** GO terms with Benjamini-Hochberg adjusted p-value less than 0.05 were clustered using the Jaccard distance calculated for sets of DE genes associated with each term. The heat map shows signed minus $\log_{10}$ adjusted p-values where positive sign indicates terms enriched in "DE up" genes, while negative sign indicates terms enriched in "DE down" genes. White cells indicate that enrichment is not significant on the given time-course day.

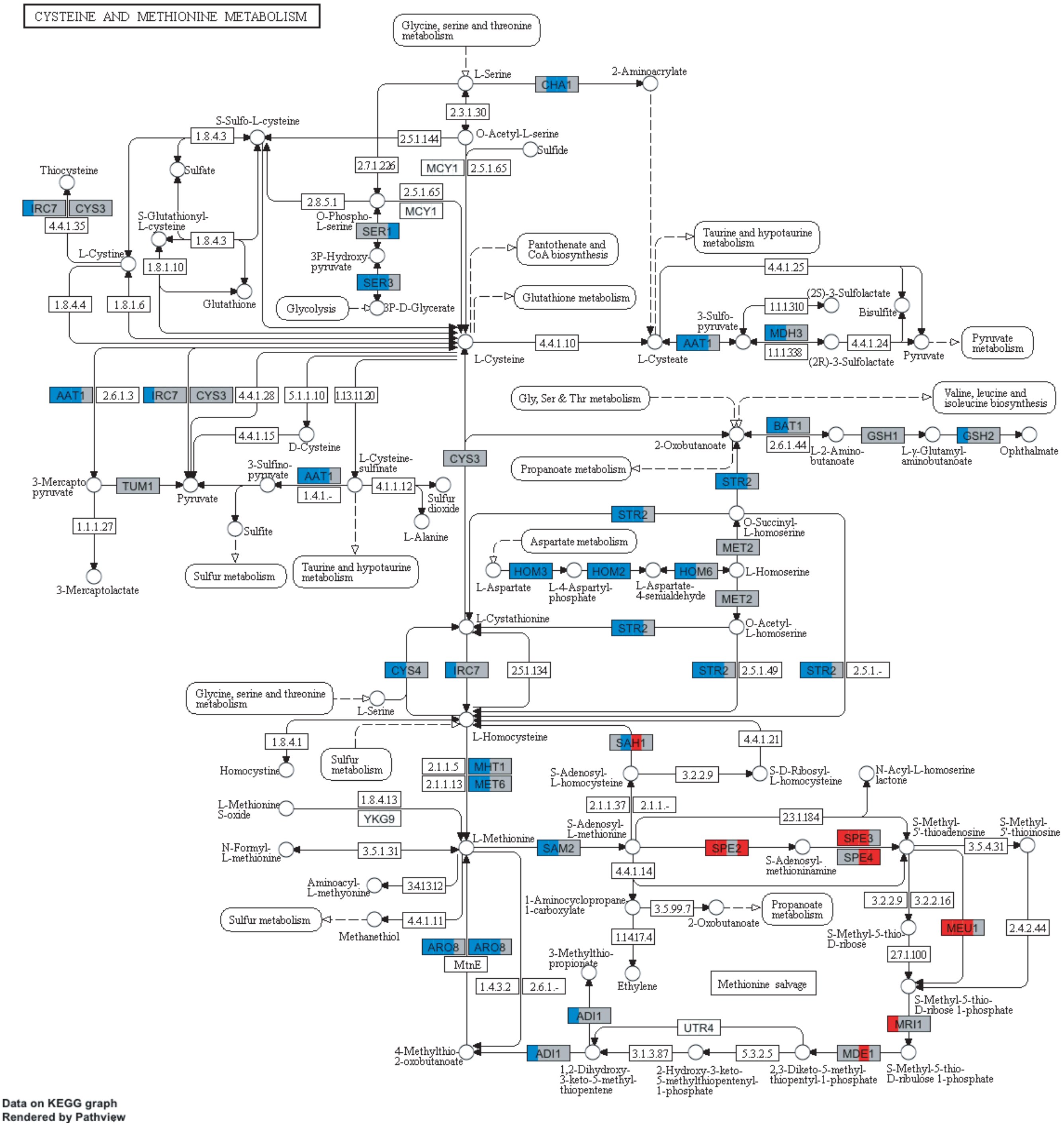


**Figure S11: Time-course differential expression of cysteine and methionine metabolism pathway genes (KEGG pathway: sce00270).** Each colored rectangle corresponds to the product of a gene that is expressed in *K. phaffii* and is homologous to a gene in *S. cerevisiae* (Homology was determined by Love *et al.,* who named *K. phaffii* genes according to their *S. cerevisiae* homologues (10)). A rectangle is partitioned into four quarters corresponding to the four time-course days, and the shading of quarters from left to right indicates differential expression status on progressive time-course days. Blue indicates “DE down” genes and red indicates “DE up” genes.

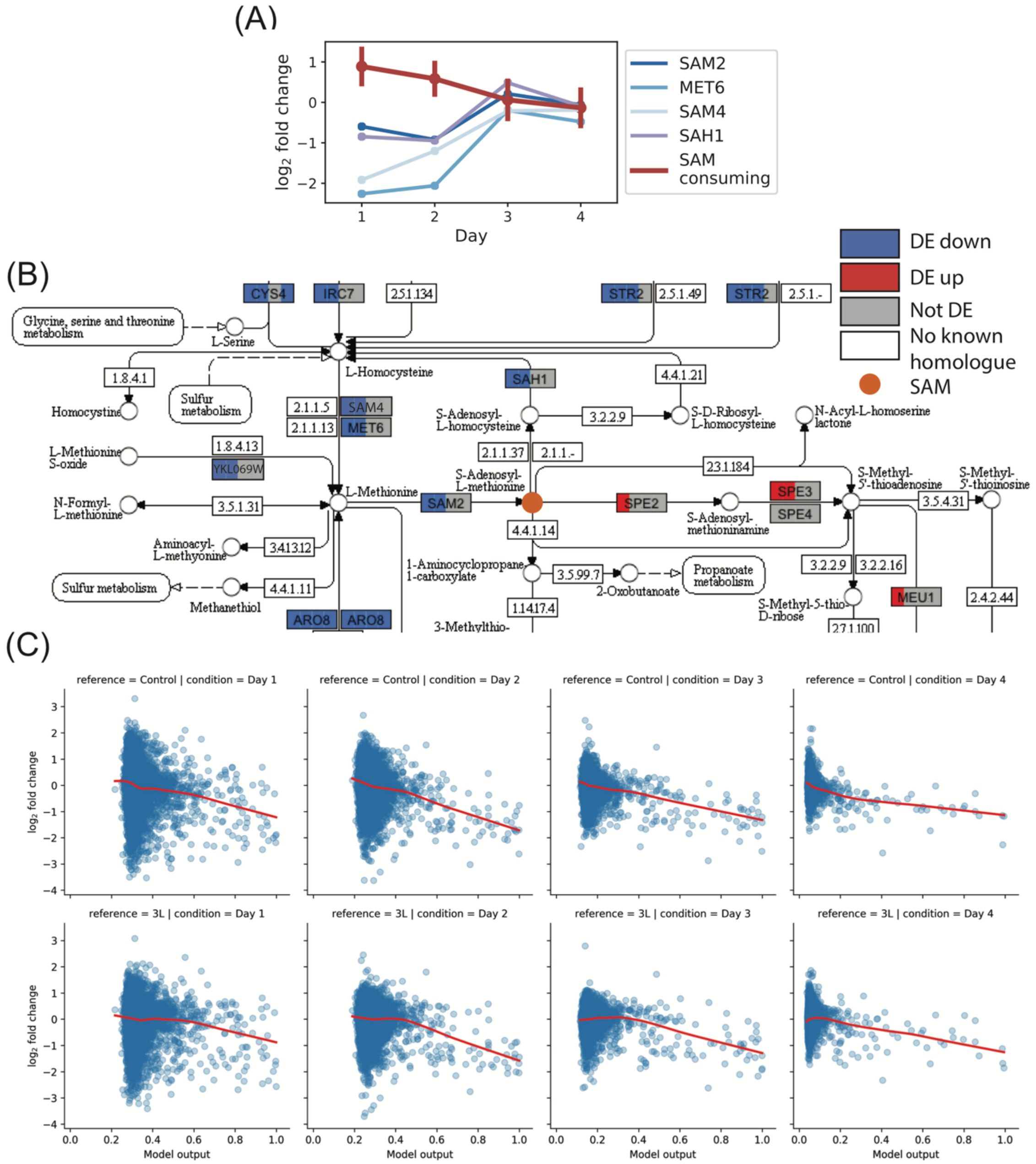


**Figure S12: Differential expression analysis using DNMT3L single knock-in as reference.** (A,B) correspond to Figure 2D,E, where the Control plasmid data were used as reference. (C) Anti-correlation of "DE down vs. rest" model output with expression $\log_2$ fold-change relative to Control (top row) and DNMT3L (bottom row) conditions. The *x*-axis of each panel is the output of the logistic regression model, where the model parameters are from Figure 3E.

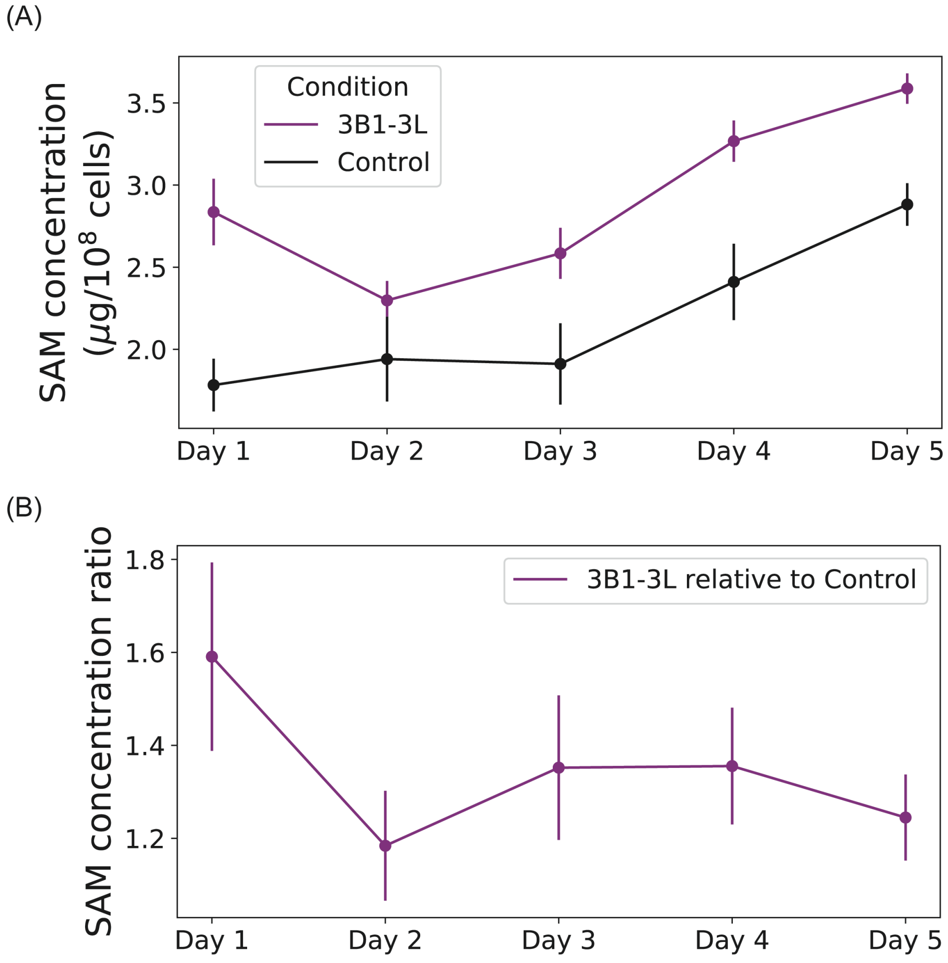


**Figure S13: SAM level in 3B1-3L knock-in cells as a function of time.** (A) Concentration of SAM in 3B1-3L knock-in cells and Control cells for time-course days 1-5, measured by the ELISA. (B) The ratio of SAM concentration in 3B1-3L to that in Control. Error bars indicate standard error calculated over 3 biological replicates.

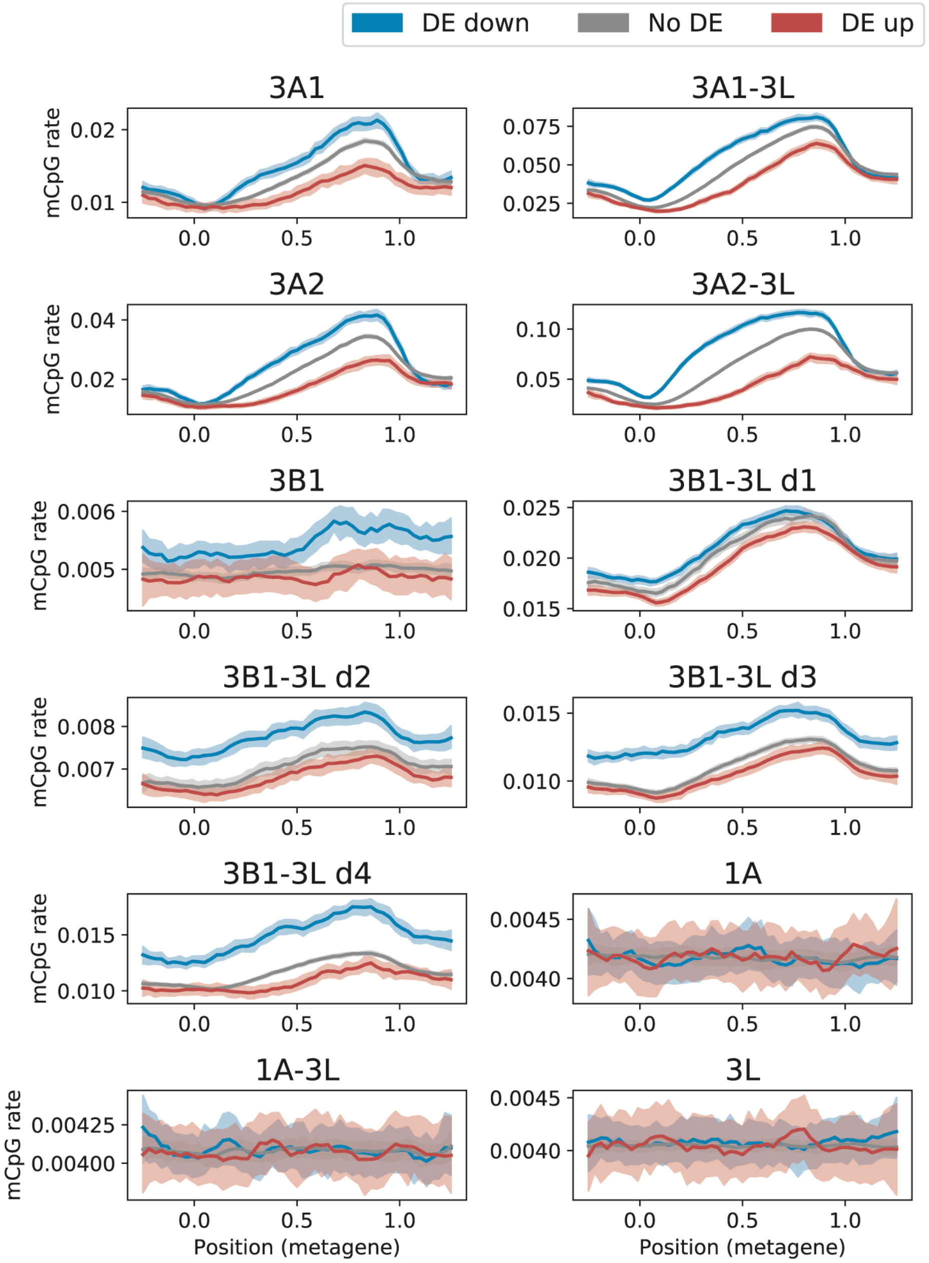


**Figure S14: Metagene plot of mCpG rates for genes grouped by differential expression state (5% FDR).** Solid lines indicate mean of Bayesian mCpG rates, and shading indicates 95% credible interval.

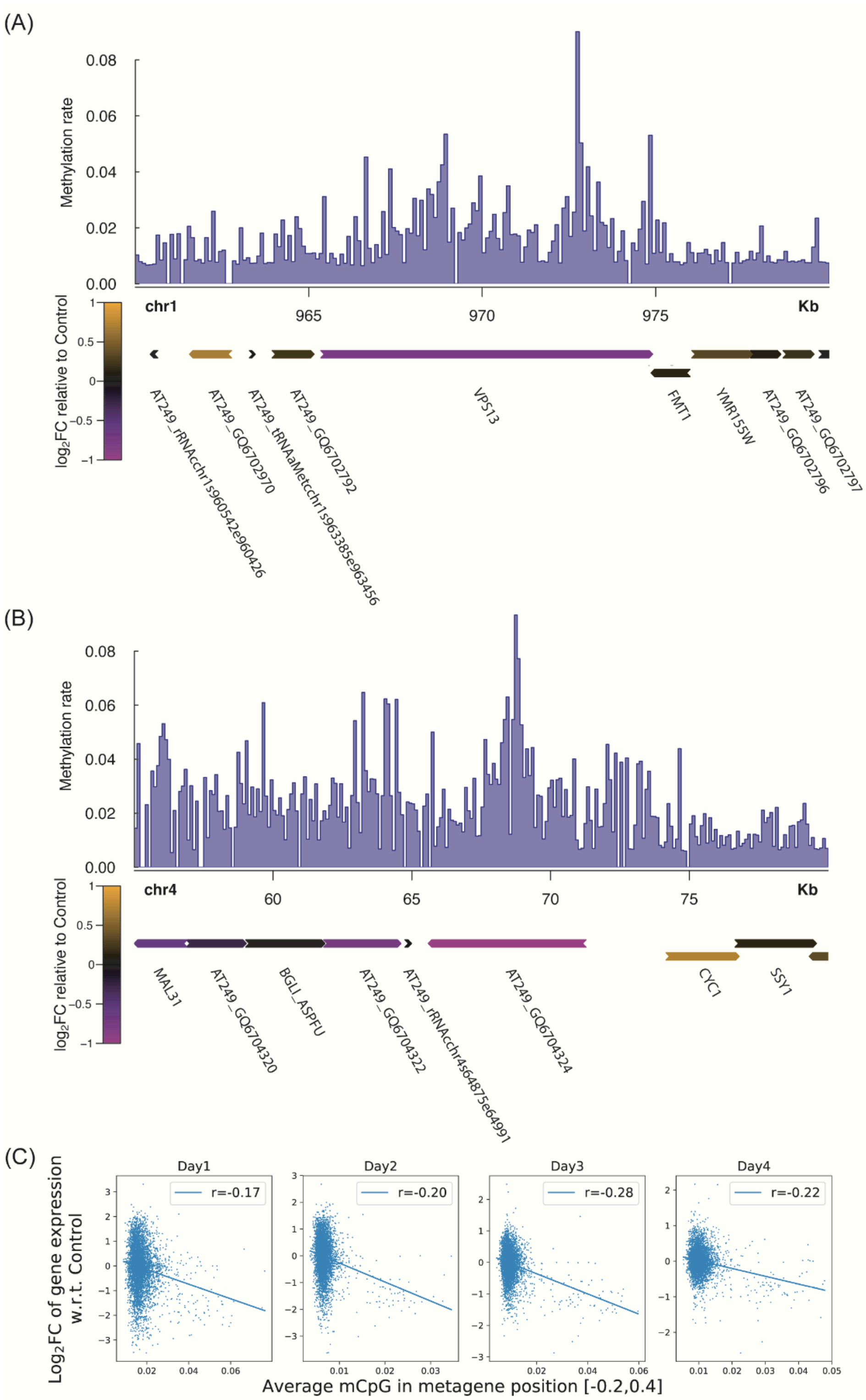


**Figure S15:** (A) Gene track in the region chr1:960,000-980,000 with corresponding methylation rate in 3B-3L d4 and expression $\log_2$ fold-change relative to Control. Arrows indicate the strand of genes. (B) Same as in (A), but in the region chr4:55,000-80,000. (C) Scatter plot of gene expression against methylation rate in 3B-3L time-course experiments. Correlation coefficient is shown in legend.

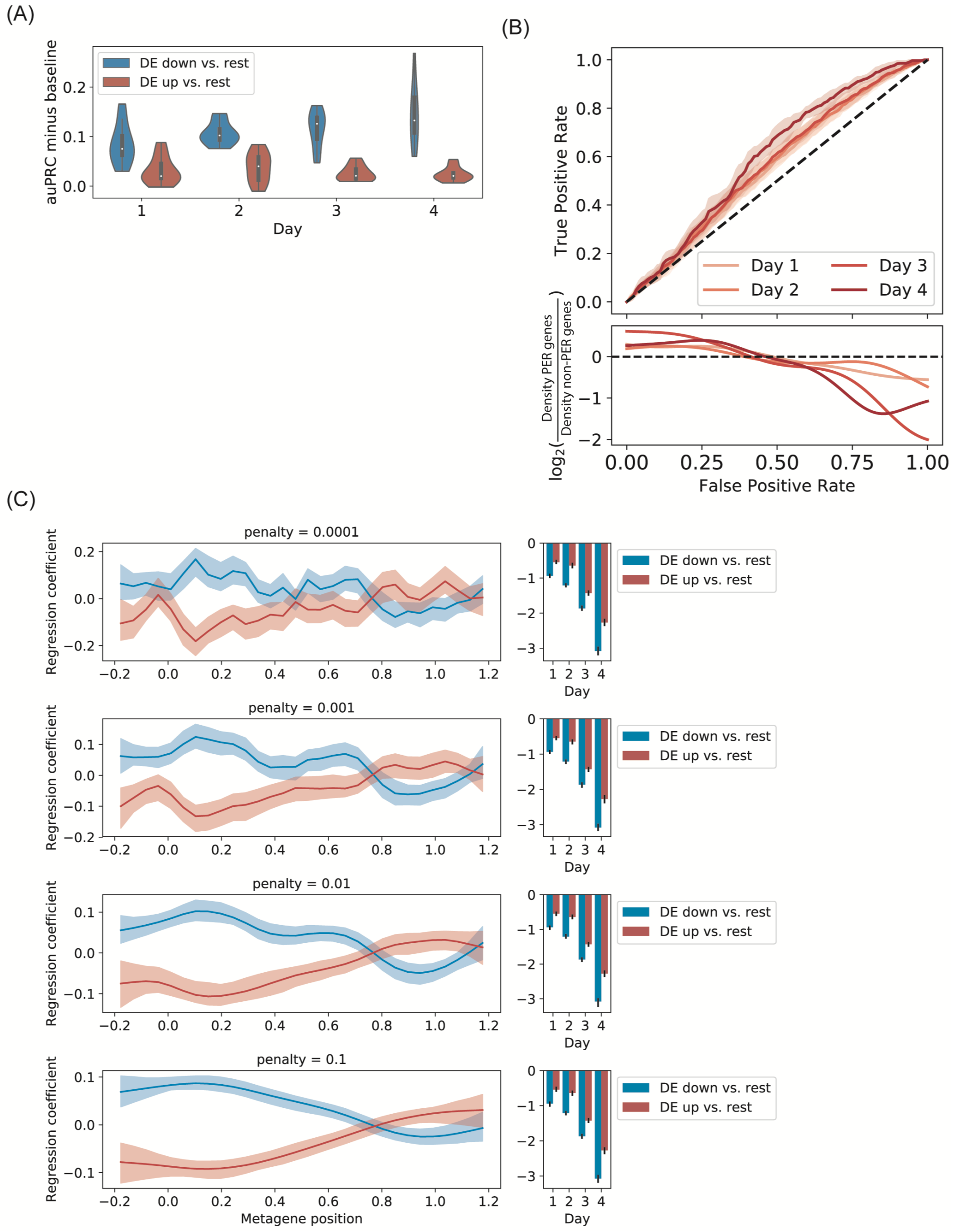


**Figure S16: Additional performance statistics for multivariate logistic regression classifier and effect of regularization parameter. (A)** Area under the precision recall curve (auPRC) minus the baseline auPRC defined in Supplementary Methods: “Choice of logistic regression smoothness penalty”. auPRC minus base line is given for “DE down vs. rest” and “DE up vs. rest” classifiers on test set data restricted to each time-course day. **(B)** Receiver operating characteristics (ROC) for classification of test set data in “DE up vs. rest” task (top) and log ratio of distributions of minimum false positive rates for correct classification of PER and non-PER “DE up” genes (bottom). Solid lines in ROC indicate mean of true positive rate for given false positive rate taken across CV folds; shading indicates 95% confidence interval**. (C)** Regression coefficients for logistic regression classifiers at different values of regularization parameter. Solid lines and bar heights indicate parameters learned from all time-course data. Shading indicates 95% confidence interval estimated by bootstrap resampling of time-course data.

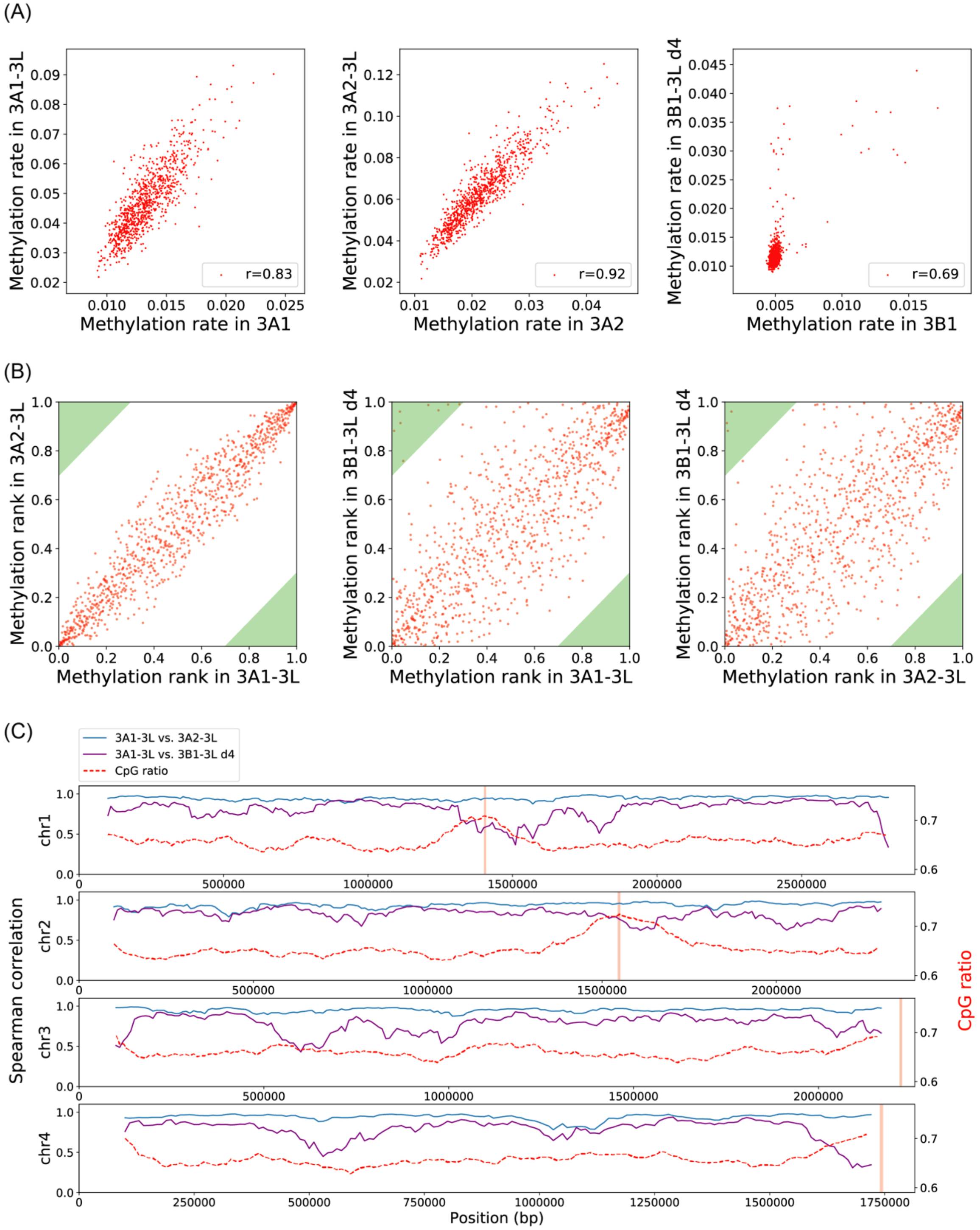


**Figure S17: Effect of DNMT3L and differential methylation patterns between DNTM3A and DNMT3B.** (A) Adding 3L linearly scales the methylation level of single DNMT knock-in. (B) Scatter plot of ranked CpG methylation rates in 10kb genomic bins in 3A1-3L vs. 3A2-3L, 3B1-3L vs. 3A1-3L and 3B1-3L vs. 3A2-3L conditions. The points in the green region represent outlier CpG sites that are highly methylated by 3B1-3L, but not by 3A1-3L or 3A2-3L. (C) Genome-wide track of Spearman correlation coefficient of methylation rates in the indicated conditions. We binned the genome in 10kb bins and calculated the Spearman correlation of average methylation rate in 20 adjacent bins between conditions. The red vertical lines indicate the centromeres. The methylation levels in 3A1-3L and 3A2-3L conditions are highly correlated (blue line), while the levels in 3A1-3L and 3B1-3L conditions (purple line) tend to deviate from each other near centromeres and telomeres, where the CpG ratio peaks. The CpG ratio is defined as the number of observed CpGs divided by the number of expected CpGs in each bin.

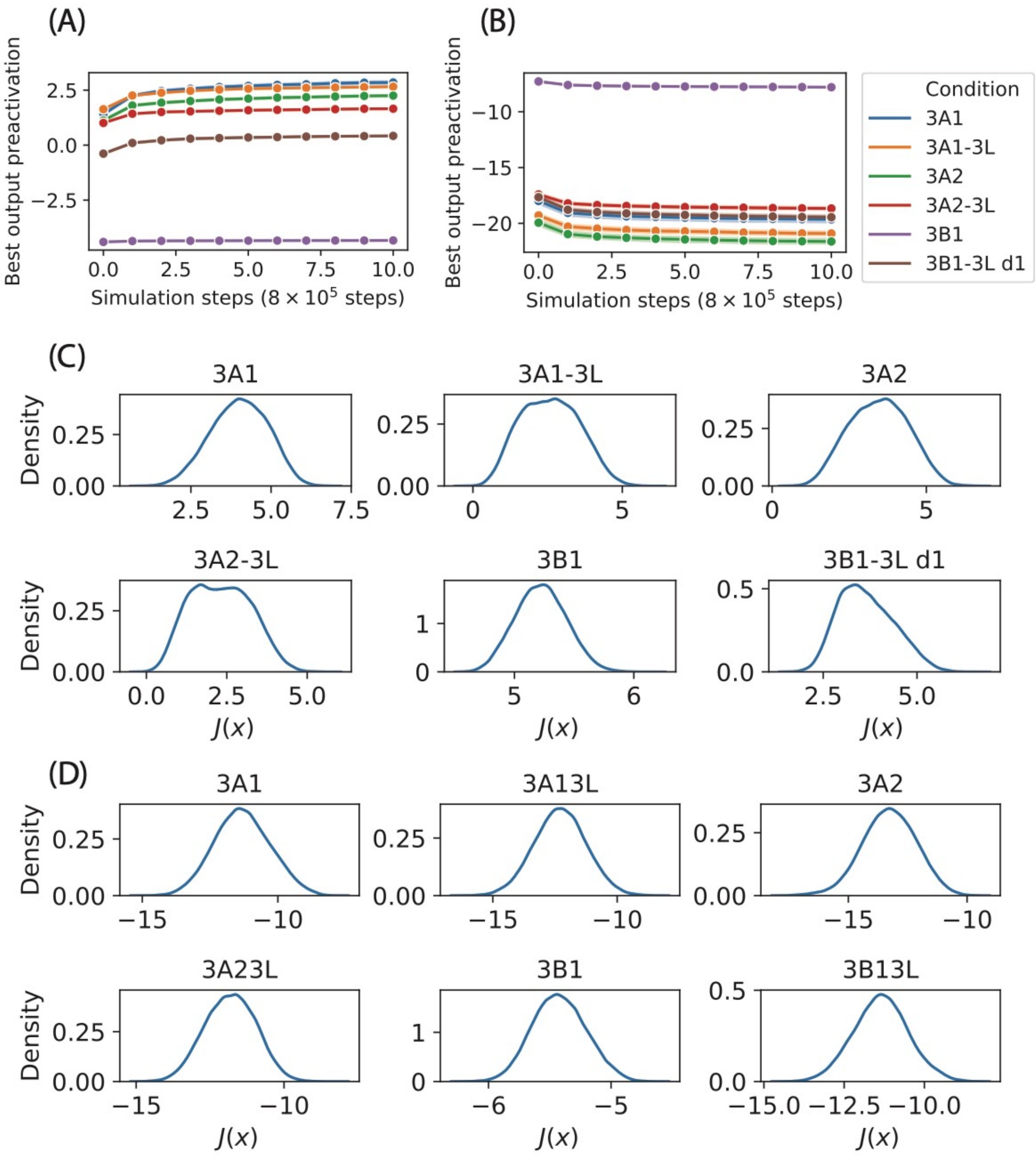


**Figure S18: Convergence of simulated annealing (SA) and variation in the width of objective function distributions with condition. (A)** Best output neuron pre-activations $y(\boldsymbol{x})$ vs. number of simulation steps for SA designed to maximize predicted methylation rates. **(B)** Best output neuron pre-activation $y(\boldsymbol{x})$ vs. number of simulation steps for SA designed to minimize predicted methylation rates. **(C)** Distributions of SA maximization objective function ($J(\boldsymbol{x}) = -y(\boldsymbol{x})$) by condition. **(D)** Distribution of SA minimization ($J(\boldsymbol{x}) = y(\boldsymbol{x})$) objective functions by condition.

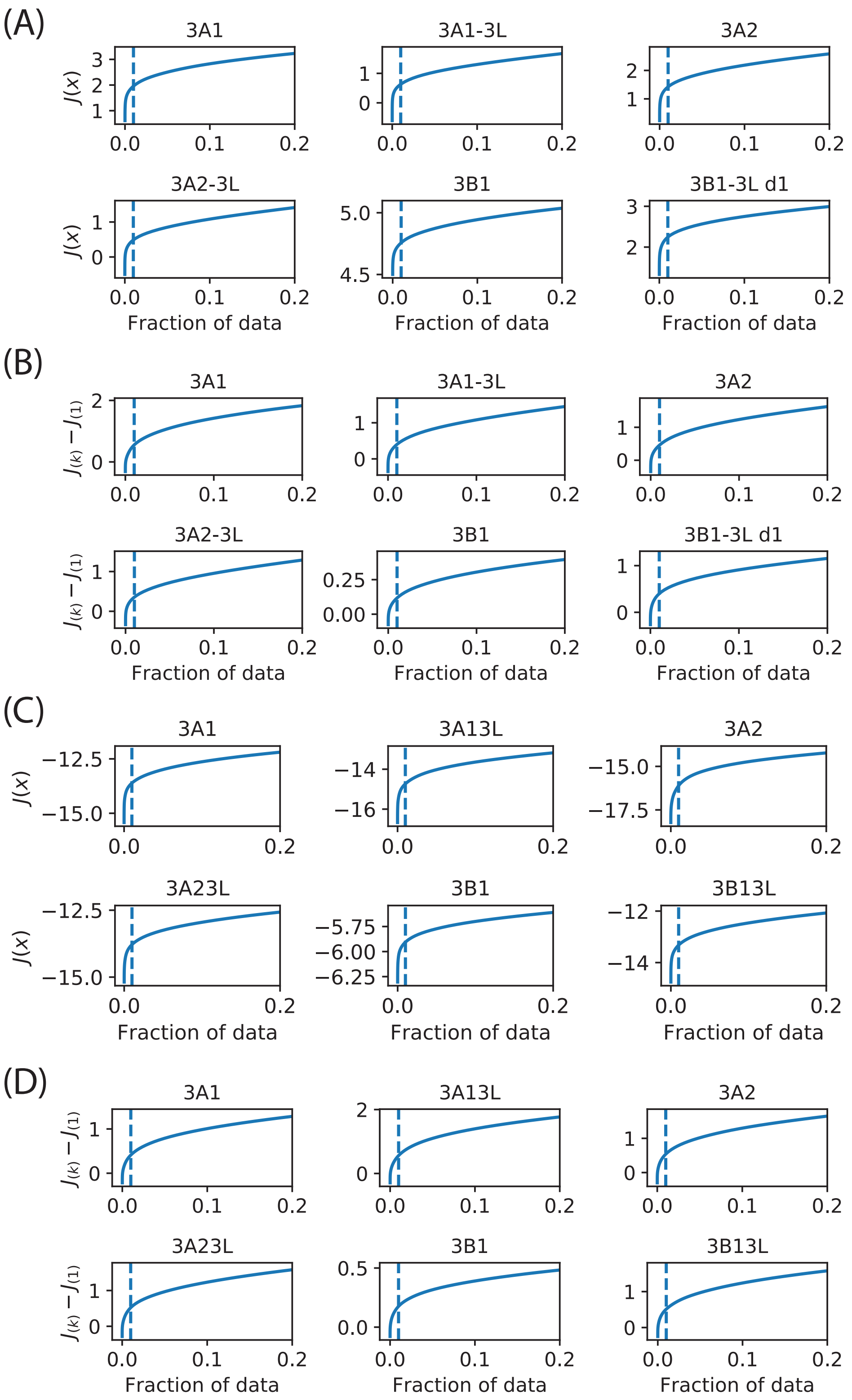


**Figure S19: Condition-specific choice of the $d$ parameter in SA. (A)** Inverse cumulative distribution function for SA maximization objective function. Dashed line illustrates objective function percentile separating "basin points" from "bulk points." **(B)** Inverse cumulative distribution function of difference between sampled SA maximization objective function outputs and minimum of objective function outputs across samples. Intersection of dashed line (at $x = 0.01$) and solid line indicates value of $d$ parameter used for SA maximization in each condition. **(C,D)** Similar to (A) and (B), but for SA minimization.

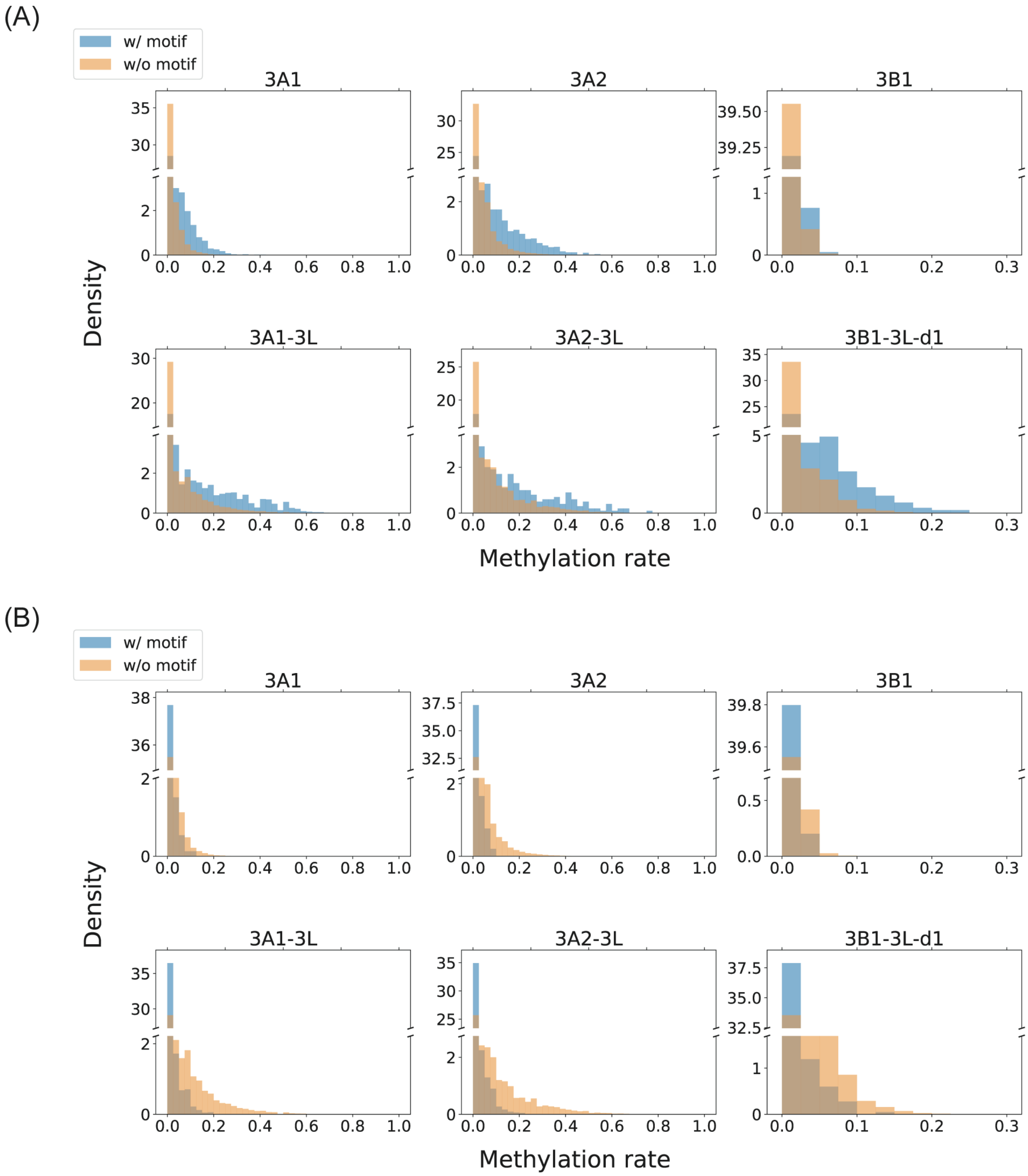


**Figure S20: Validation of the motifs identified by SA**. **(A)** Distribution of mCpG rates at CpG cytosines contained vs. not contained in motifs preferred by each DNMT (Figure 5A). **(B)** Distribution of mCpG rates at CpG cytosines contained vs. not contained in motifs avoided by each DNMT (Figure 5B).

(A)

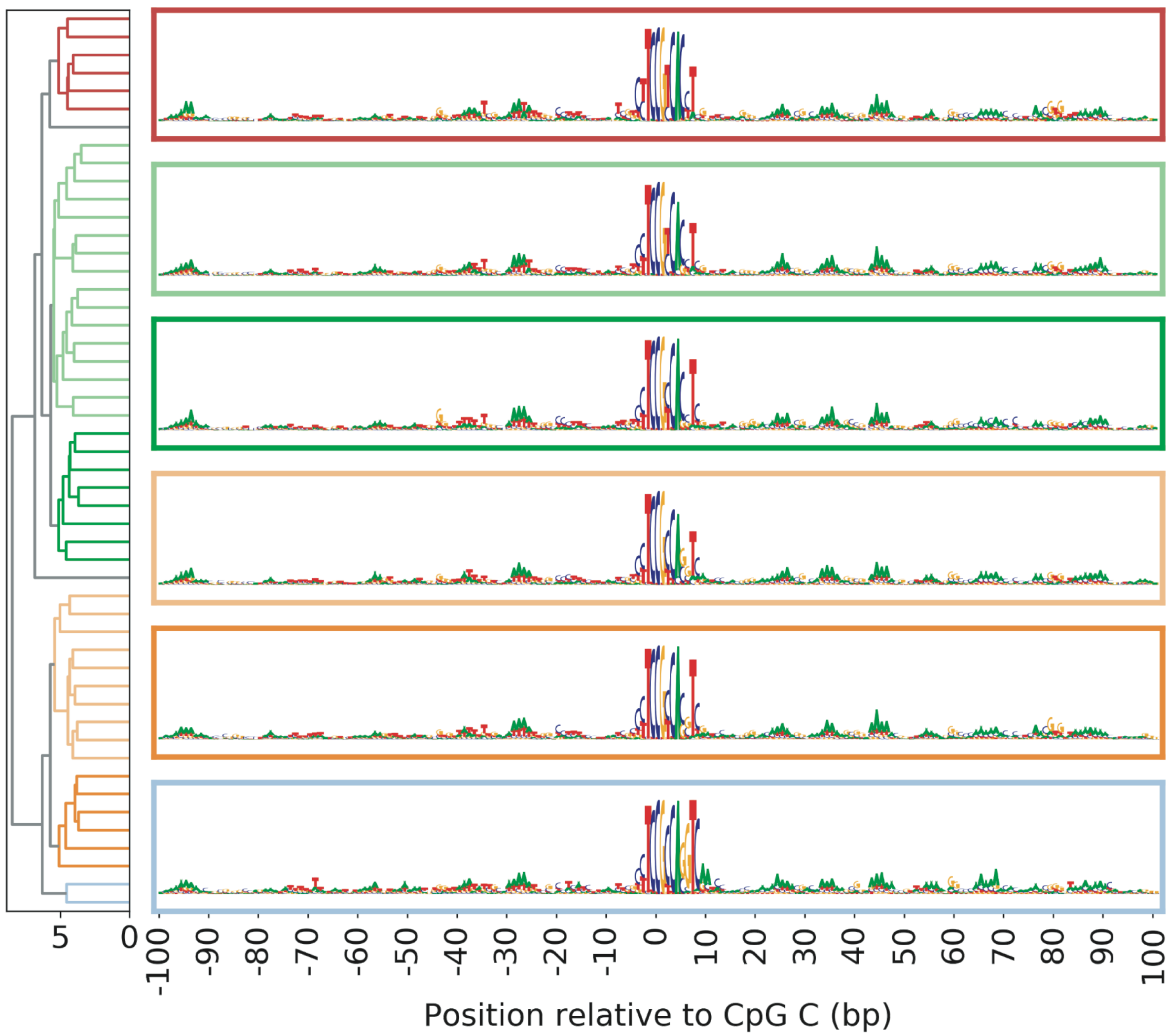

5
0
-100
-90
-80
-70
-60
-50
-40
-30
-20
-10
0
10
20
30
40
50
60
70
80
90
100
Position relative to CpG C (bp)


(B)

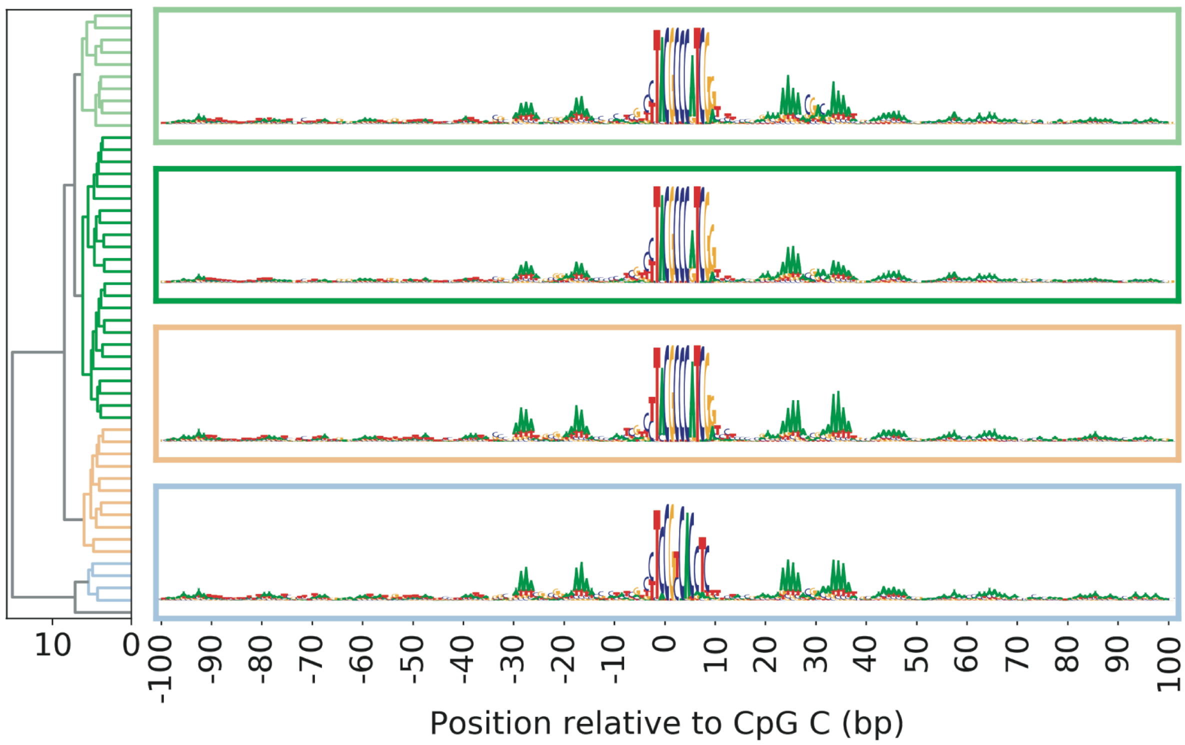

10
0
-100
-90
-80
-70
-60
-50
-40
-30
-20
-10
0
10
20
30
40
50
60
70
80
90
100
Position relative to CpG C (bp)

(C)

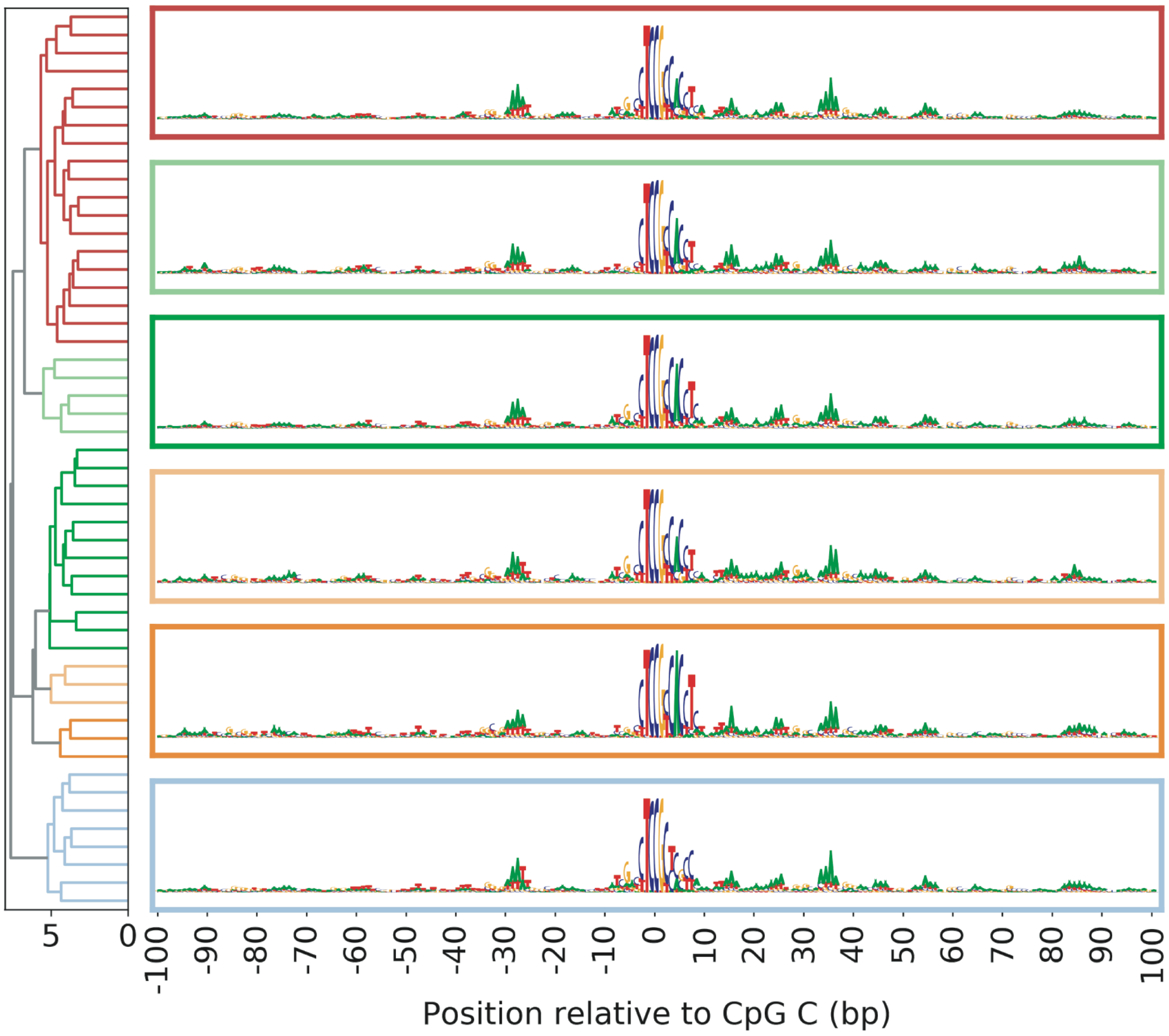

5
0
-100 -90 -80 -70 -60 -50 -40 -30 -20 -10 0 10 20 30 40 50 60 70 80 90 100
Position relative to CpG C (bp)


(D)

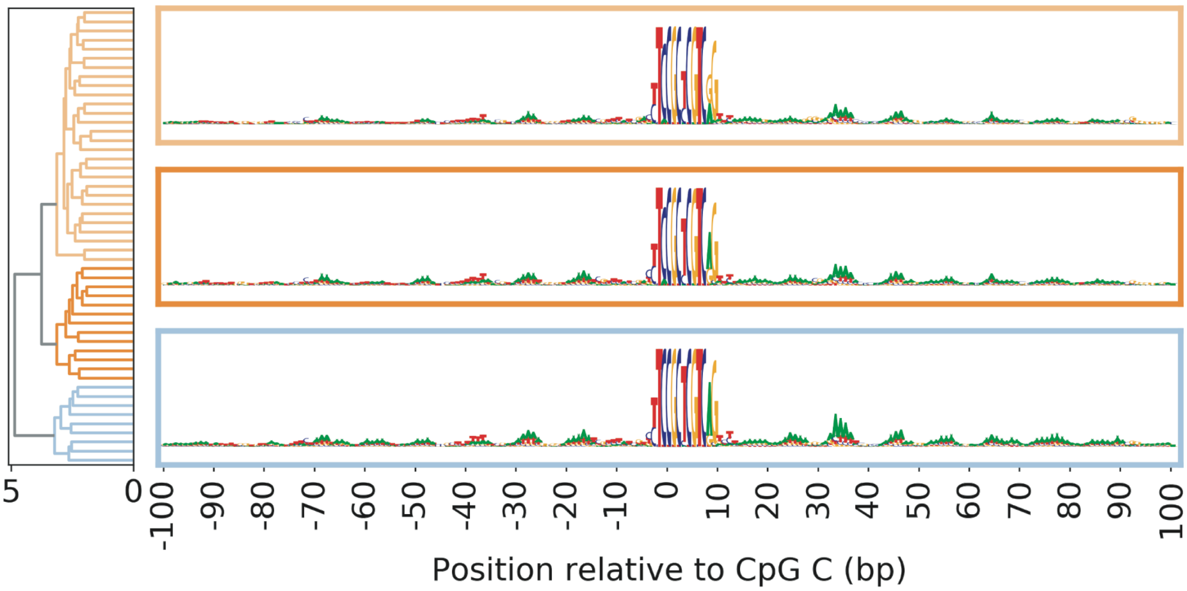

5
0
-100 -90 -80 -70 -60 -50 -40 -30 -20 -10 0 10 20 30 40 50 60 70 80 90 100
Position relative to CpG C (bp)

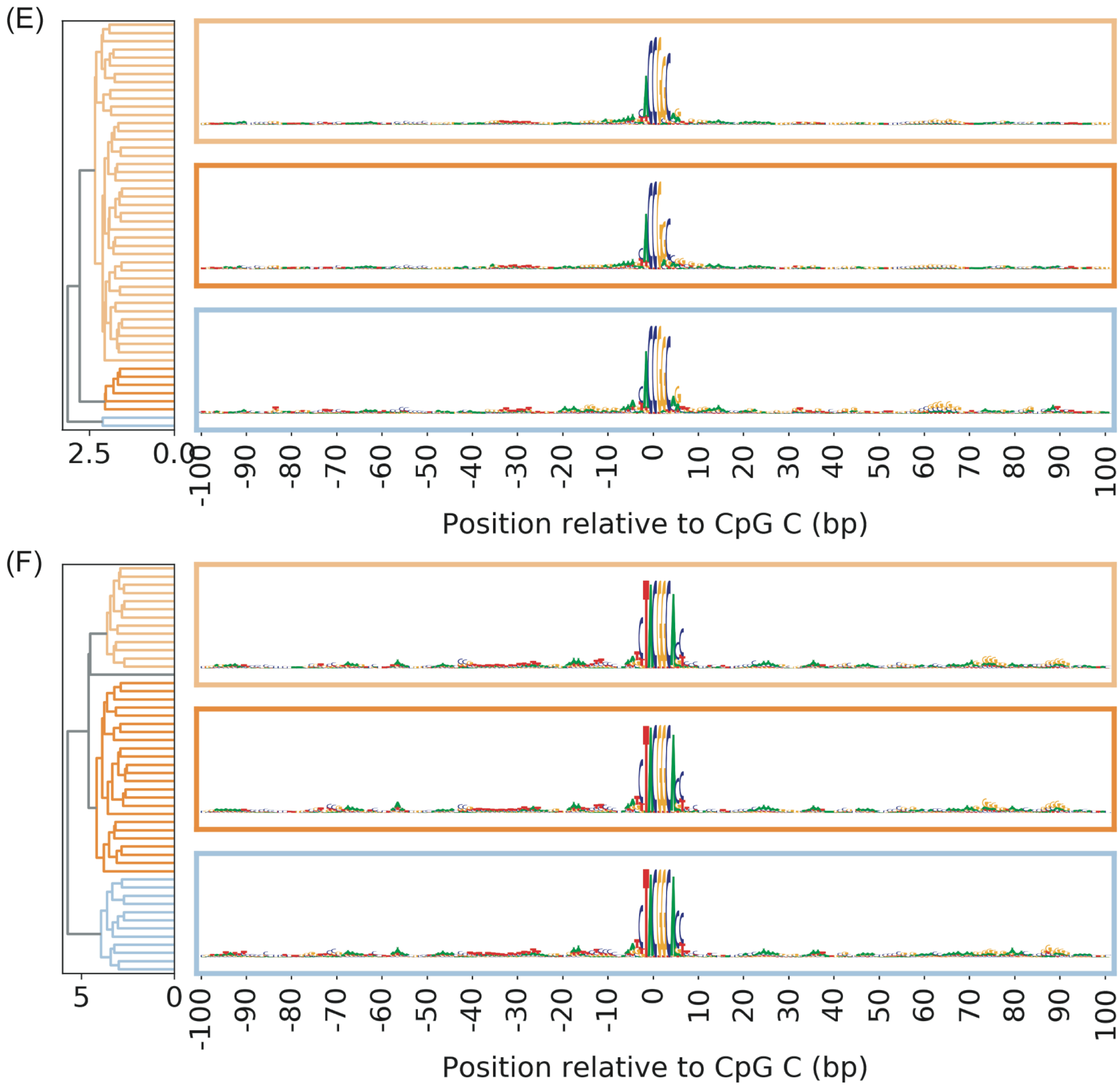


**Figure S21: Clustering of sequences predicted to maximize methylation** in **(A)** 3A1, **(B)** 3A1-3L, **(C)** 3A2, **(D)** 3A2-3L, **(E)** 3B1 and **(F)** 3B1-3L. The sequences were sampled in 50 iteration of SA interpretation for each of six conditions.

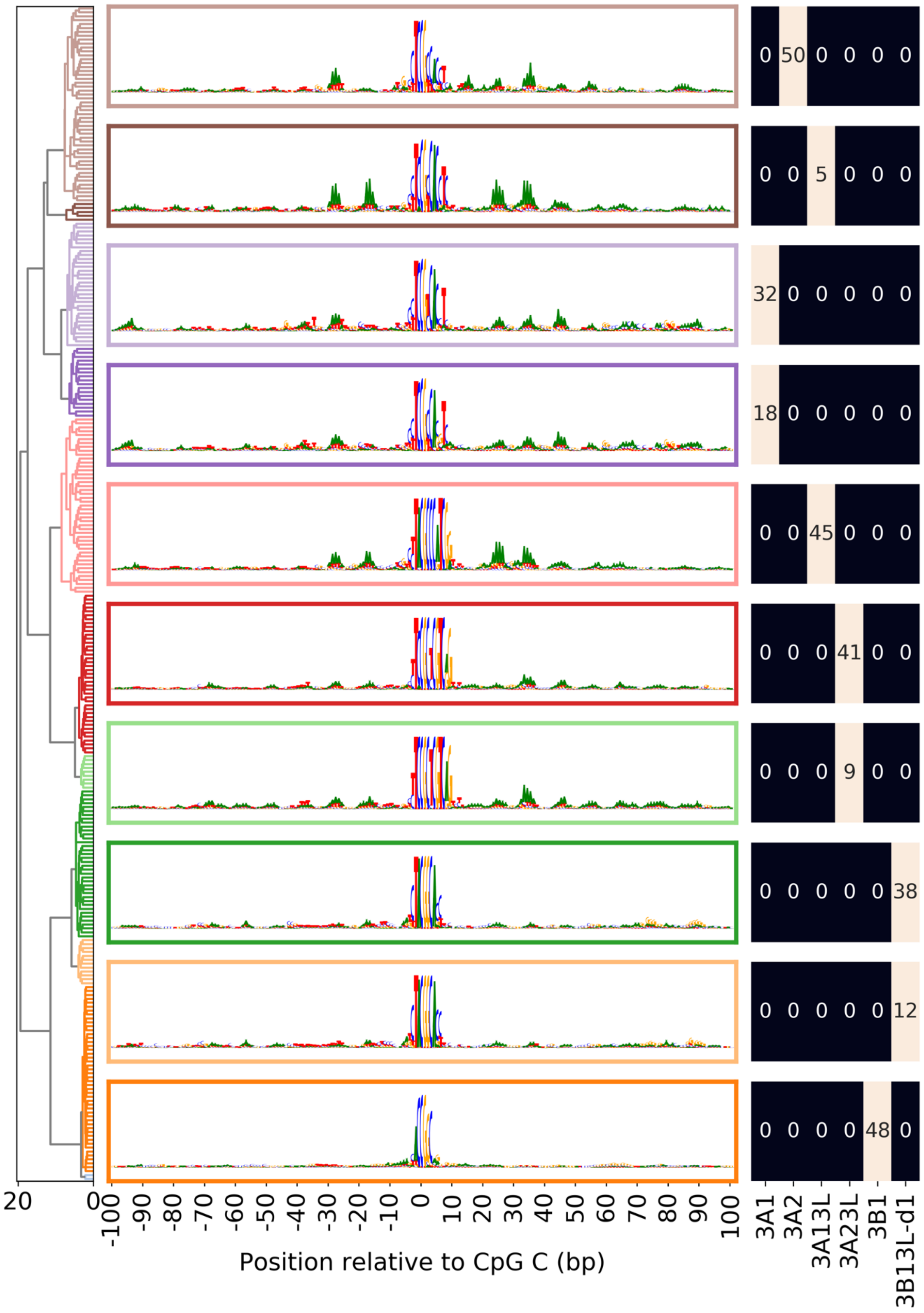


**Figure S22: Motif logos representing distinct patterns of sequence preferences identified in 300 iterations of SA interpretation.** The table on the right shows composition of each cluster in terms of SA results from each knock-in condition.

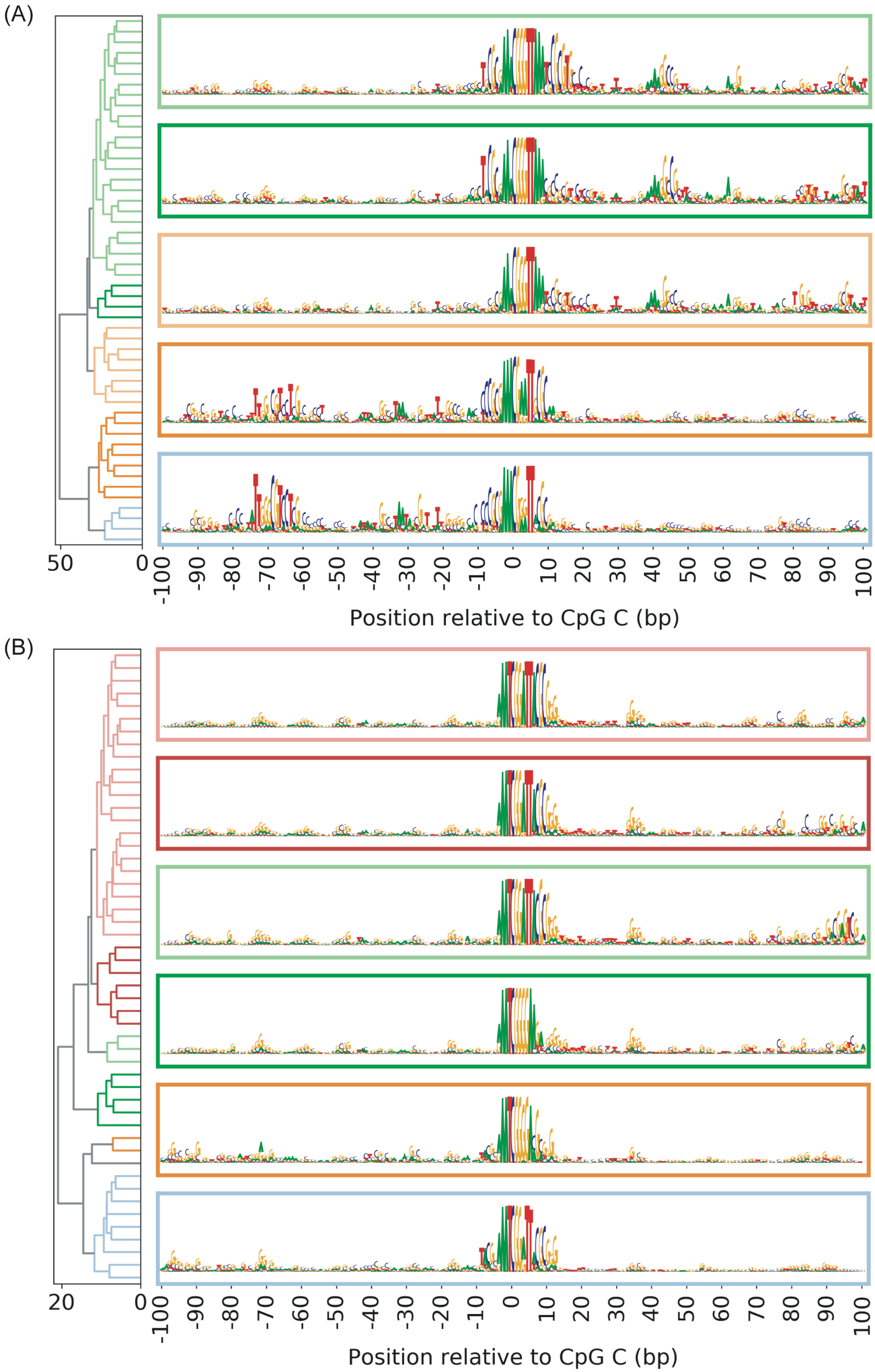
(A)
50
0
-100
-90
-80
-70
-60
-50
-40
-30
-20
-10
0
10
20
30
40
50
60
70
80
90
100
Position relative to CpG C (bp)
(B)
20
0
-100
-90
-80
-70
-60
-50
-40
-30
-20
-10
0
10
20
30
40
50
60
70
80
90
100
Position relative to CpG C (bp)

(C)

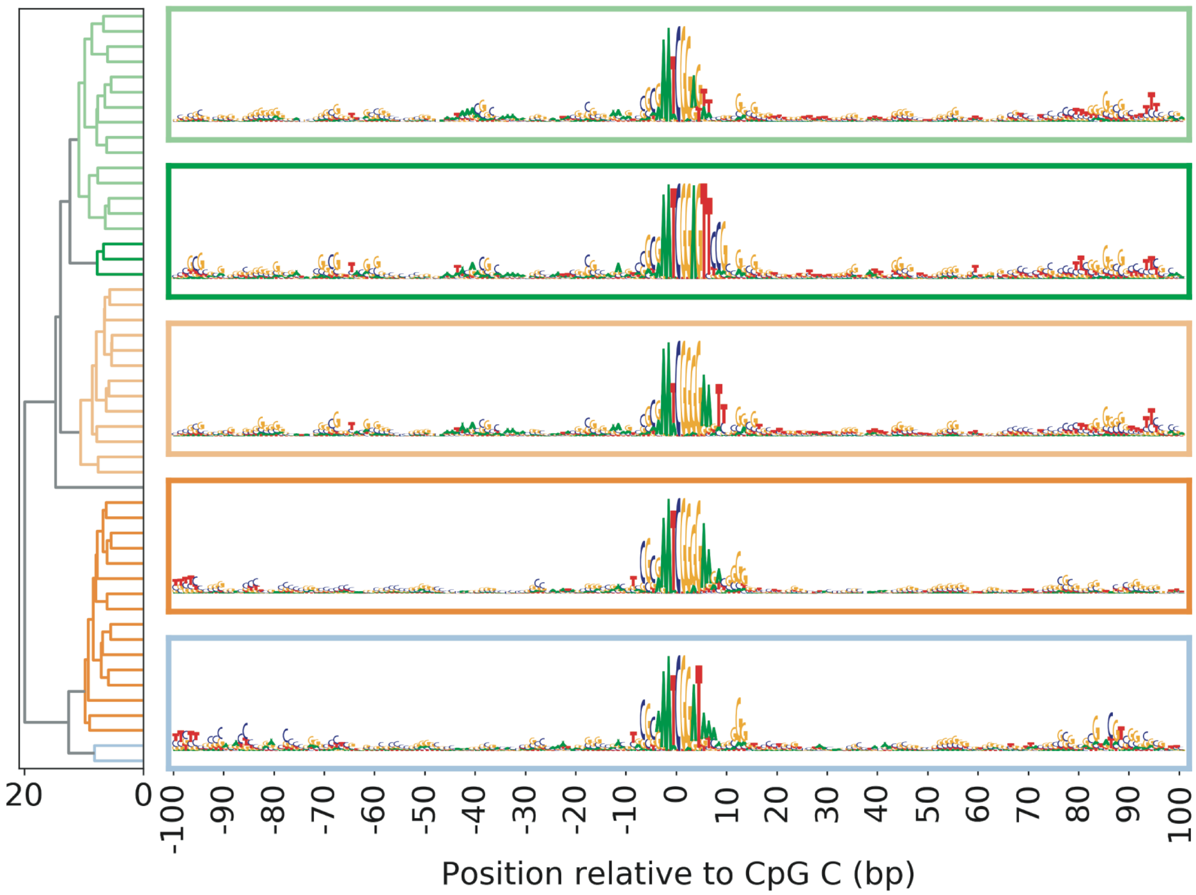
20
0
-100 -90 -80 -70 -60 -50 -40 -30 -20 -10 0 10 20 30 40 50 60 70 80 90 100
Position relative to CpG C (bp)

(D)

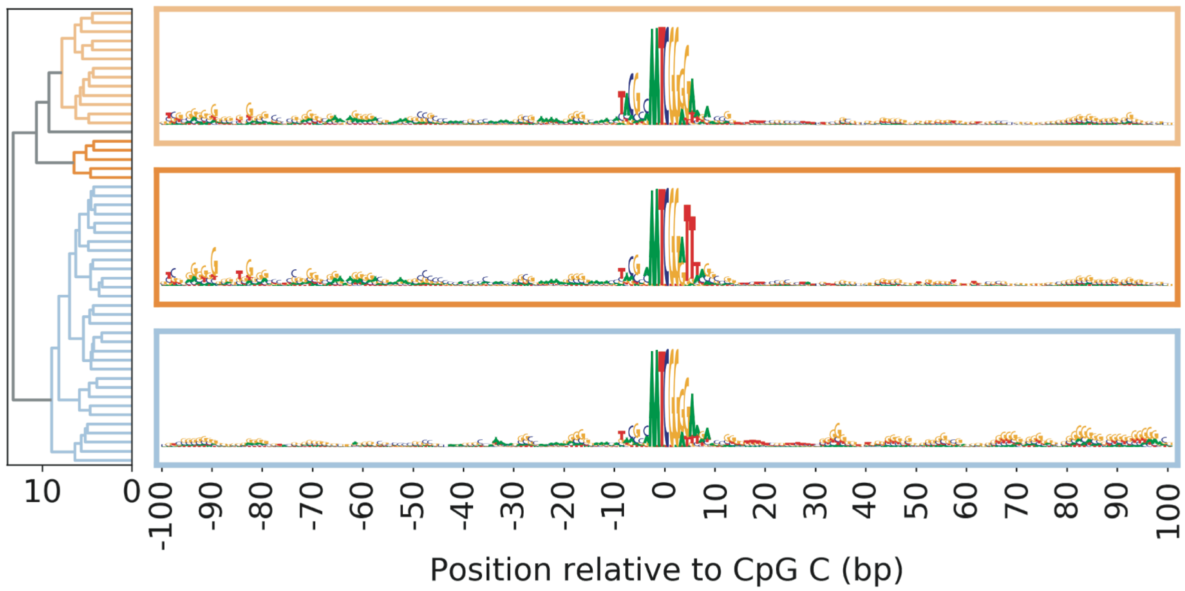
10
0
-100 -90 -80 -70 -60 -50 -40 -30 -20 -10 0 10 20 30 40 50 60 70 80 90 100
Position relative to CpG C (bp)

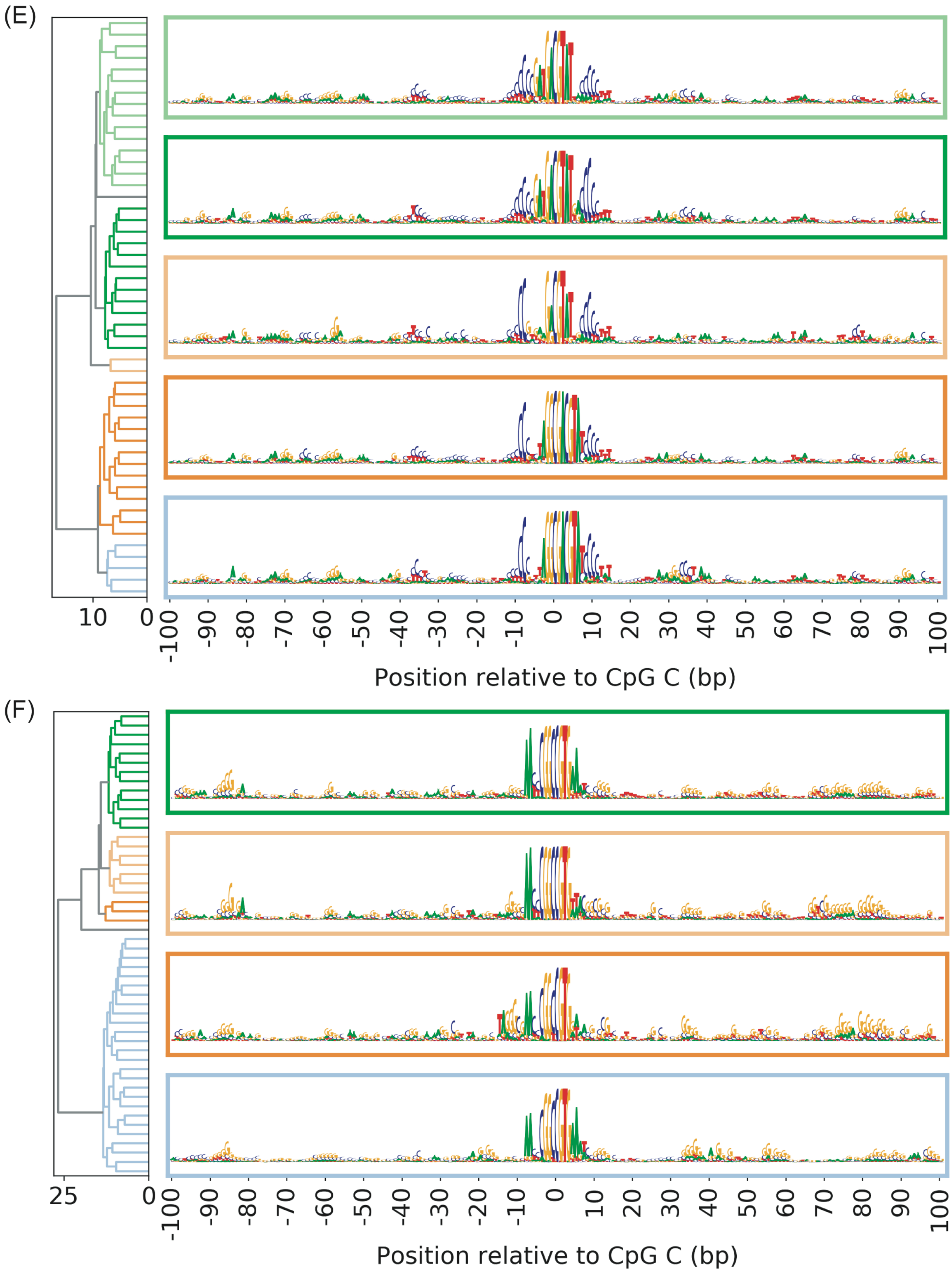


**Figure S23**: **Clustering of sequences predicted to minimize methylation** in **(A)** 3A1, **(B)** 3A1-3L, **(C)** 3A2, **(D)** 3A2-3L, **(E)** 3B1 and **(F)** 3B1-3L. The sequences were sampled in 50 iteration of SA interpretation for each of six conditions.

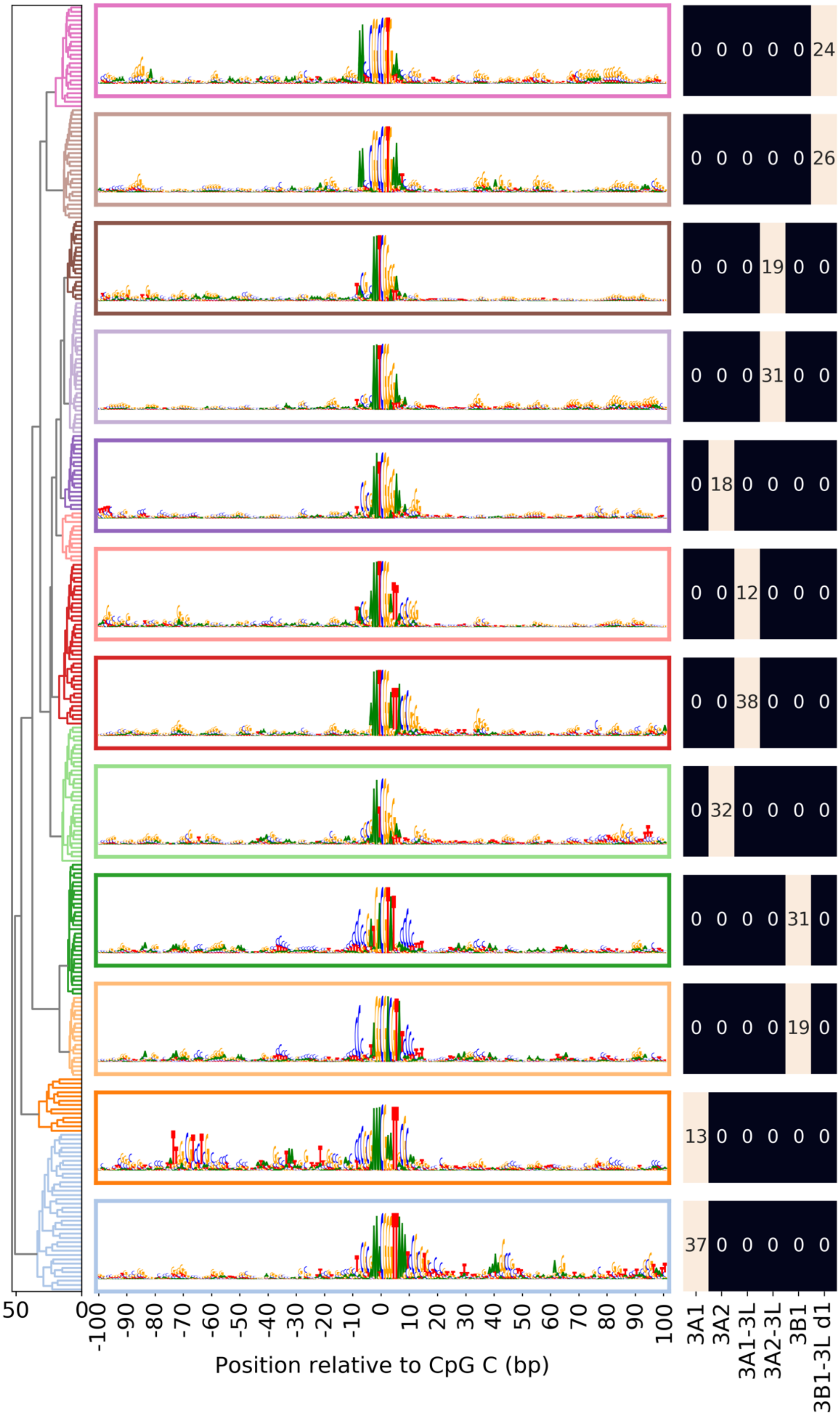


**Figure S24: Motif logos representing distinct patterns of sequence aversions identified in 300 iterations of SA interpretation.** The table at the right shows composition of each cluster in terms of SA results from each knock-in condition.

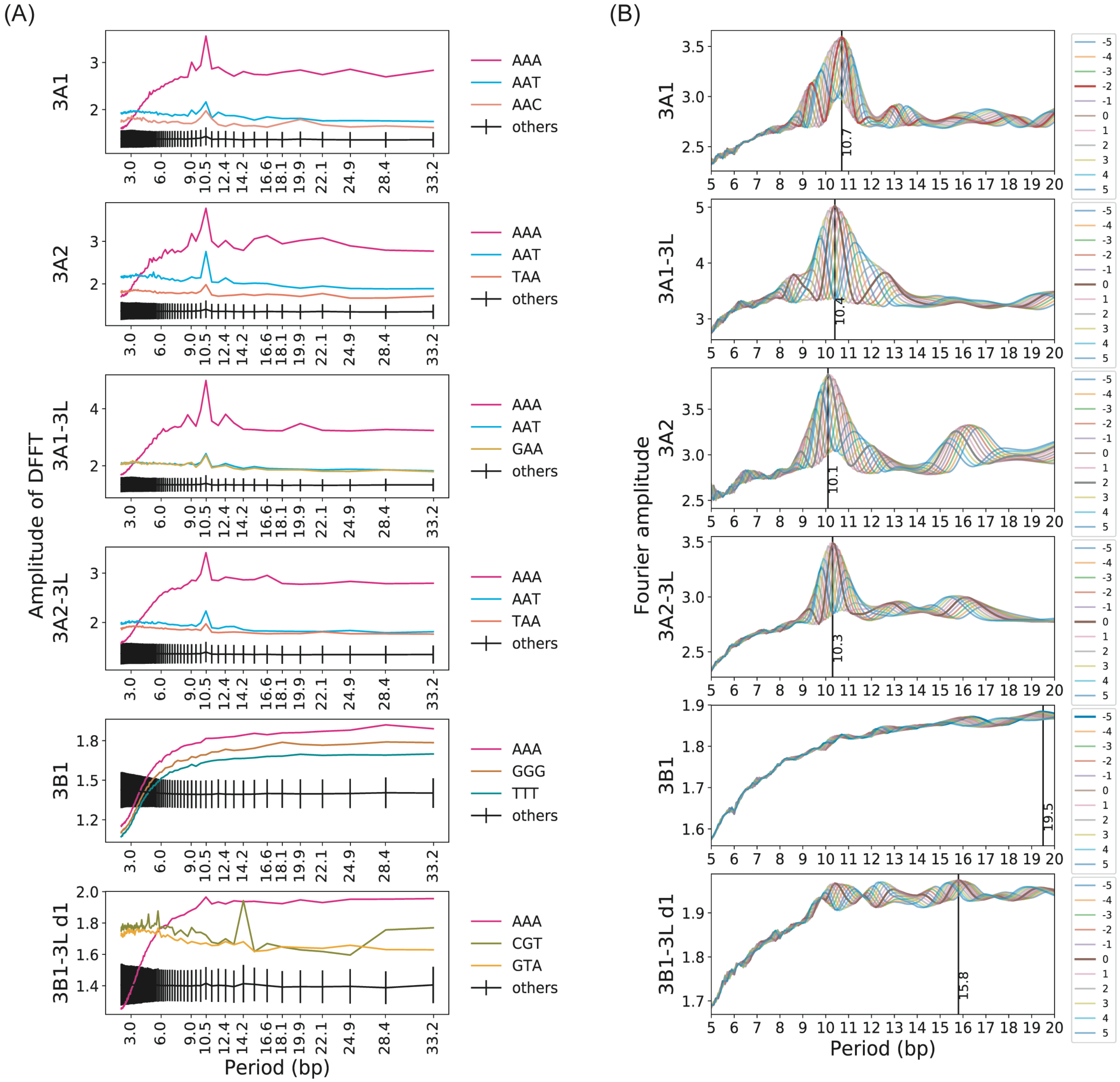


**Figure S25: Periodicity of trinucleotides in motifs predicted by SA to be preferred by DNMTs and their phase difference. (A)** DFFT applied to trinucleotide counts in individual sequences sampled by SA and then averaged. The top three trinucleotides with highest amplitude at 10.5 bp are shown individually, and amplitudes of the remaining 3-mers are summarized by their mean and standard deviations (Supplementary Methods: "Fourier analysis of periodic trinucleotides"). **(B)** Dot product between counts of each top trinucleotides in panel (A) and discrete complex exponential functions with period varying from 5 to 20 bp. Different colored curves show the amplitude of dot product after shifting the location of trinucleotide counts 3' of central CpG by the indicated amount of bases, while fixing the location of trinucleotide counts 5' of the central CpG (Figure S22).

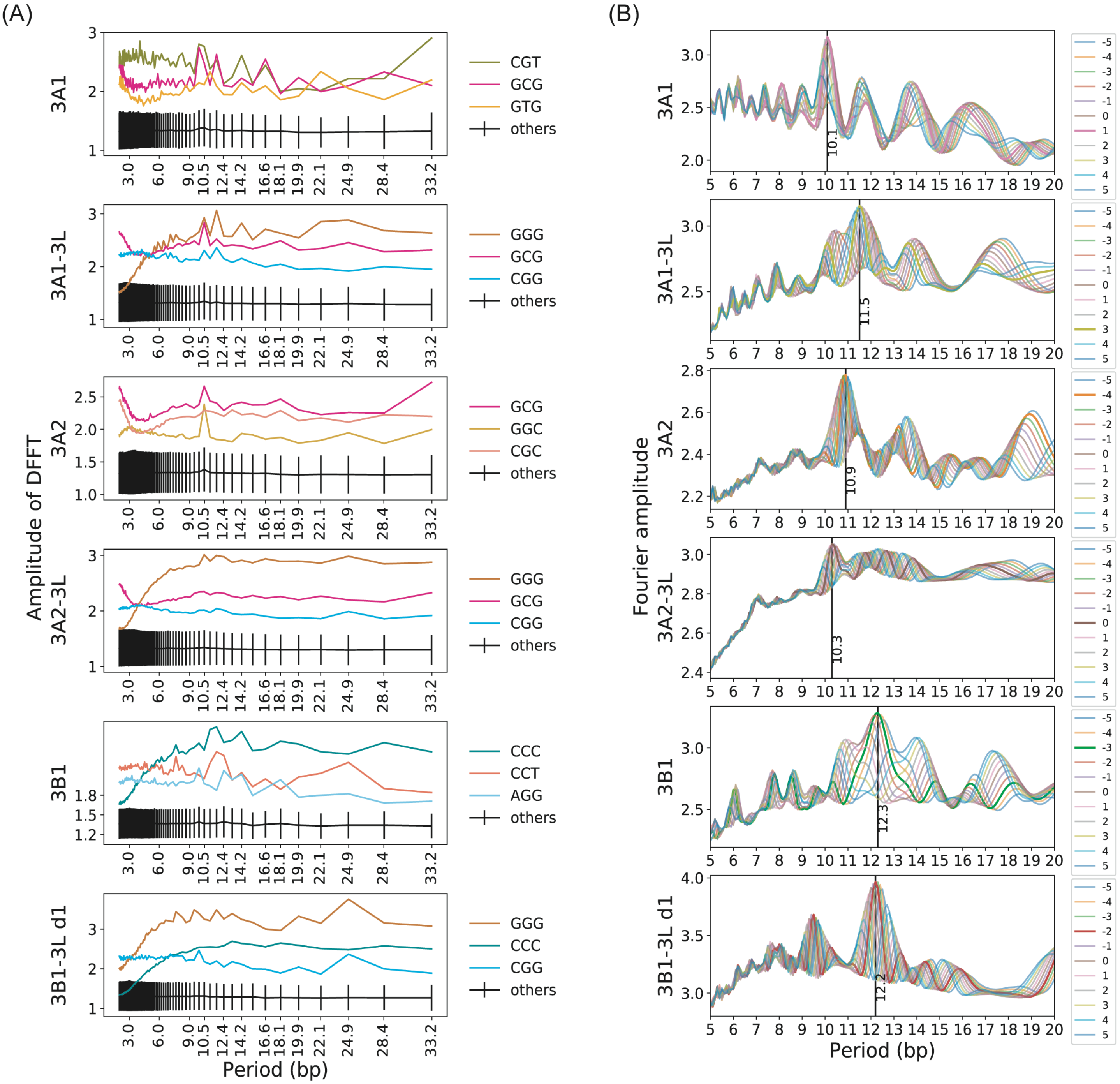


**Figure S26: Periodicity of trinucleotides in motifs predicted by SA to be avoided by DNMTs and their phase difference. (A)** DFFT applied to trinucleotide counts in individual sequences sampled by SA and then averaged. The top three trinucleotide with highest amplitude at 10.5 bp are shown individually, and amplitudes of the remaining trinucleotide are summarized by their mean and standard deviations (Supplementary Methods: "Fourier analysis of periodic trinucleotides"). **(B)** Dot product between counts of each top trinucleotides in panel (A) and discrete complex exponential functions with period varying from 5 to 20 bp. Different colored curves show the amplitude of dot product after shifting the location of trinucleotide counts 3' of central CpG by the indicated amount of bases, while fixing the location of trinucleotide counts 5' of the central CpG (Figure S24).

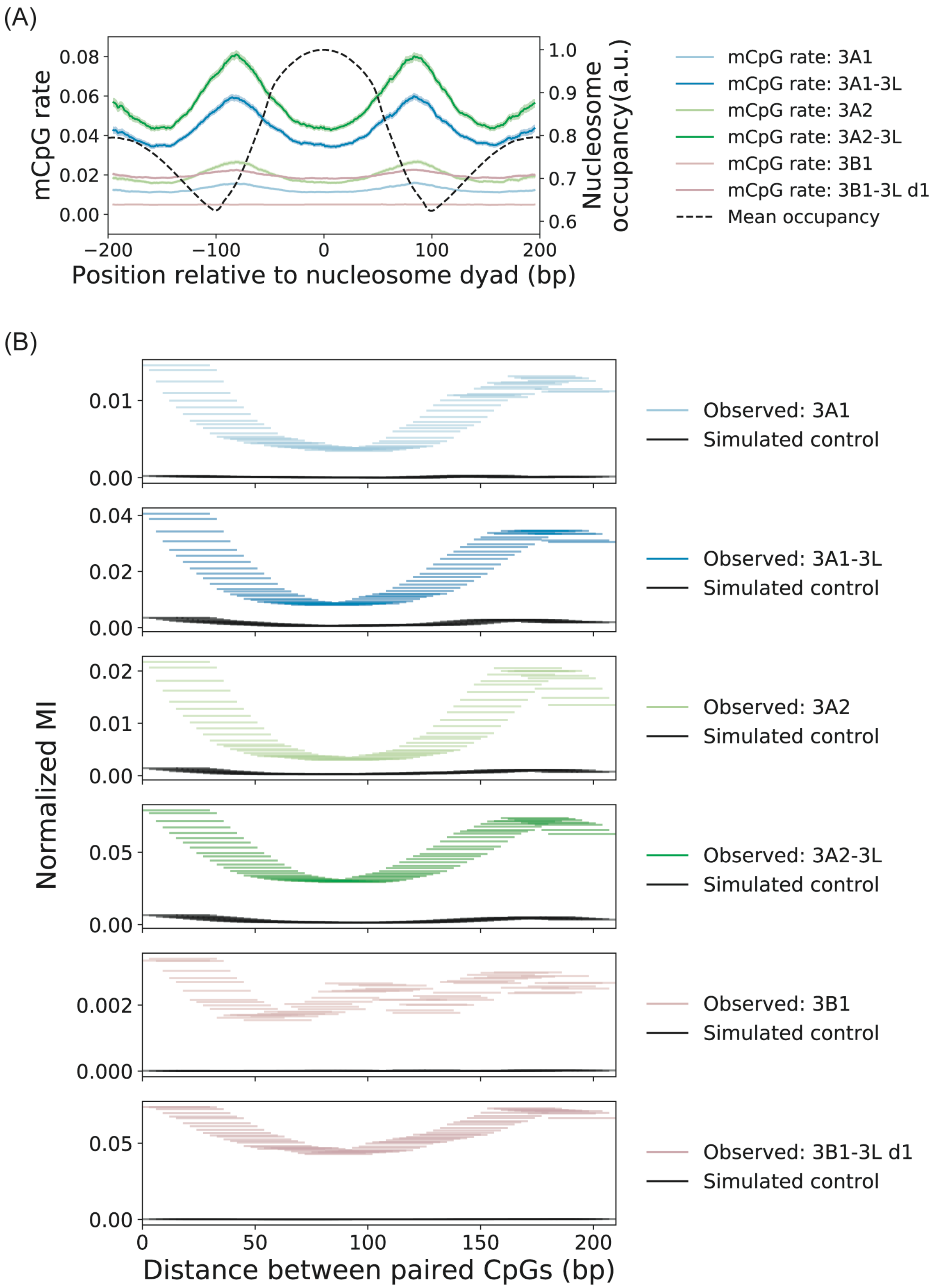


**Figure S27: (A)** Dyad-aligned mCpG rates and nucleosome occupancy. Shading indicates 95% of confidence interval. **(B)** Mutual information (MI) of methylation status at two distinct CpG sites as a function of their separation distance, normalized by marginal entropy. MI was estimated from the empirical joint distribution of methylation status at CpG pairs separated by a genomic distance within each indicated horizontal window (30 bp window width). Simulated negative control is based on independent sampling of binary methylation status from the site-specific mCpG rates.
